# TOPOVIBL-REC114 interaction regulates meiotic DNA double-strand breaks

**DOI:** 10.1101/2021.11.30.470517

**Authors:** Alexandre Nore, Ariadna B Juarez-Martinez, Julie Clément, Christine Brun, Boubou Diagouraga, Corinne Grey, Henri Marc Bourbon, Jan Kadlec, Thomas Robert, Bernard de Massy

## Abstract

Meiosis requires the formation of programmed DNA double strand breaks (DSBs), essential for fertility and for generating genetic diversity. DSBs are induced by the catalytic activity of the TOPOVIL complex formed by SPO11 and TOPOVIBL. To ensure genomic integrity, DNA cleavage activity is tightly regulated, and several accessory factors (REC114, MEI4, IHO1, and MEI1) are needed for DSB formation in mice. How and when these proteins act is not understood. Here, we show that REC114 is a direct partner of TOPOVIBL, and identified their conserved interacting domains by structural analysis. We then analysed the role of this interaction by monitoring meiotic DSBs in female and male mice carrying point mutations in TOPOVIBL that decrease or disrupt its binding to REC114. In these mutants, DSB activity was strongly reduced genome-wide in oocytes, and only in sub-telomeric regions in spermatocytes. In addition, in mutant spermatocytes, DSB activity was delayed in autosomes. These results provide evidence that REC114 is a key member of the TOPOVIL catalytic complex, and that the REC114/TOPOVIBL interaction ensures the efficiency and timing of DSB activity.

## Introduction

Sexual reproduction relies on the specialized cell division of meiosis to generate haploid cells that eventually differentiate into gametes. In most taxa, proper segregation of the homologous chromosomes (homologs) depends on homologous recombination, which physically connects homologs through at least one reciprocal exchange (crossover) between each homolog pair ^1^. Meiotic recombination initiates at the onset of prophase I by the formation of hundreds of programmed DNA double strand breaks (DSBs) at preferred DNA sites, named hotspots ^2^. DSBs are formed by the collective action of a conserved set of proteins: the TOPOVIL complex and its accessory partners ^3–6^.

The TOPOVIL complex is evolutionarily related to the TopoVI type IIB topoisomerase ^7^, and is composed of two conserved subunits: the SPO11 catalytic subunit and TOPOVIBL. TOPOVIBL shares homology with the GHKL-ATPase domain, with the central transducer domain and to a lesser extent, with the regulatory C-terminal domain (CTD) of the TopoVIB subunit of TopoVI ^5, 6, 8–11^. The TOPOVIL meiosis-specific activity is finely regulated in time (i.e. must be turned on and off at precise time windows) and in space (i.e. active at specific chromosomal locations) during meiotic prophase. However, the molecular mechanisms underlying this complex regulation remain largely elusive.

In *Saccharomyces cerevisiae*, Rec102, the identified TOPOVIBL homolog, shares only partial similarity with TopoVIB and not with its GHKL-ATPase domain ^5^. Interaction and biochemical studies suggest that Rec104, which is also essential for DSB formation in yeast, could replace the GHKL domain and that the Rec102/Rec104 complex fulfills the function of TOPOVIBL ^12–14^. The *S. cerevisiae* accessory partners, Rec114, Mei4 and Mer2 interact with DNA that mediates the formation of condensate-like RMM protein clusters ^15^. These condensates might recruit the Spo11/Rec102/Rec104/Ski8 complex to DNA through an interaction of Rec114 with Rec102/Rec104 ^12,15,16^. The finding that the Rec114 residues involved in this interaction are essential for DSB formation supports this hypothesis ^15^.

In mammals, DSB sites are determined by PRDM9 that recognizes specific DNA motifs and modifies chromatin upon binding to these sites ^17–19^. In the mouse, the accessory proteins REC114, MEI4, IHO1 (orthologues of *S. cerevisiae* Rec114, Mei4 and Mer2 respectively) and MEI1 are essential for DSB formation, and localize as foci on chromosome axes at meiotic prophase onset. It has been proposed that IHO1, MEI4 and REC114 directly control TOPOVIL through its recruitment or activation ^20–24^. MEI4 and REC114 form a stable complex, and structural analyses revealed that REC114 N-terminus forms a Pleckstrin homology (PH) domain ^23, 25^ harbouring exposed conserved residues potentially involved in protein-protein interactions. This suggests that REC114 acts as a regulatory platform. In line with this hypothesis, it was recently reported that ANKRD31 directly interacts with REC114 PH domain, and is involved in regulating DSB number and localization ^25, 26^. However, how these accessory proteins participate in TOPOVIL activity remains to be determined.

Here, using structural analysis, we found that the mouse REC114 PH domain directly interacts with a conserved C-terminal peptide of TOPOVIBL, identifying its CTD as a regulatory unit of the TOPOVIL complex. Accordingly, in mice where TOPOVIBL interaction with REC114 was disrupted, we observed meiotic DSB formation defects in both sexes associated by reduced fertility. In females, DSB formation was drastically reduced. In males, DSB formation was delayed, but DSB levels were not reduced excepted for sub-telomeric regions. On the X and Y chromosomes, which recombine specifically in the distal sub-telomeric pseudo-autosomal region (PAR) region, DSB activity reduction led to chromosome synapsis defects. These results provide evidence that REC114 is part of the TOPOVIL catalytic complex and acts to regulate the level and timing of DSB formation.

## Results

### TOPOVIBL forms a stable complex with REC114

Using yeast two-hybrid assays we identified a specific interaction between mouse TOPOVIBL and REC114. The deletion analysis showed that the N-terminal PH domain of REC114 was required for this interaction because TOPOVIBL did not bind to REC114 lacking the first 39 aa (40-259) (Figs. 1a and 1b). This interaction also required TOPOVIBL C-terminus as indicated by the absence of binding upon deletion of its last 29aa (TOPOVIBL 1-550) (Figs. 1a and 1c). TOPOVIBL 1-550 could still interact with SPO11β, indicating that the deletion of the 29aa did not disrupt TOPOVIBL folding (Fig. 1c). We obtained similar results *in vitro* using Strep-tag pull-down assays (Figs. 1d and 1e). While it was difficult to produce full-length TOPOVIBL, we could express and purify a 6xHis-SUMO fusion TOPOVIBL protein that lacks part of its transducer domain (construct 1-385; Fig. 1d, lane 1) and also the 6xHis-TOPOVIBL CTD (450-579) (Fig. 1d, lane 2). REC114 clearly interacted with TOPOVIBL 450-579 (Fig. 1d, lane 5), but not with TOPOVIBL 1-385 (Fig. 1d, lane 4). REC114 N-terminal domain (REC114 1-159), which includes the PH domain, was sufficient for the interaction with TOPOVIBL 450-579 (Fig. 1e, lane 2). In agreement, the REC114 PH domain and TOPOVIBL 450-579 co-eluted in a single peak during size exclusion chromatography (Extended data Fig. 1a).

**Fig. 1.**
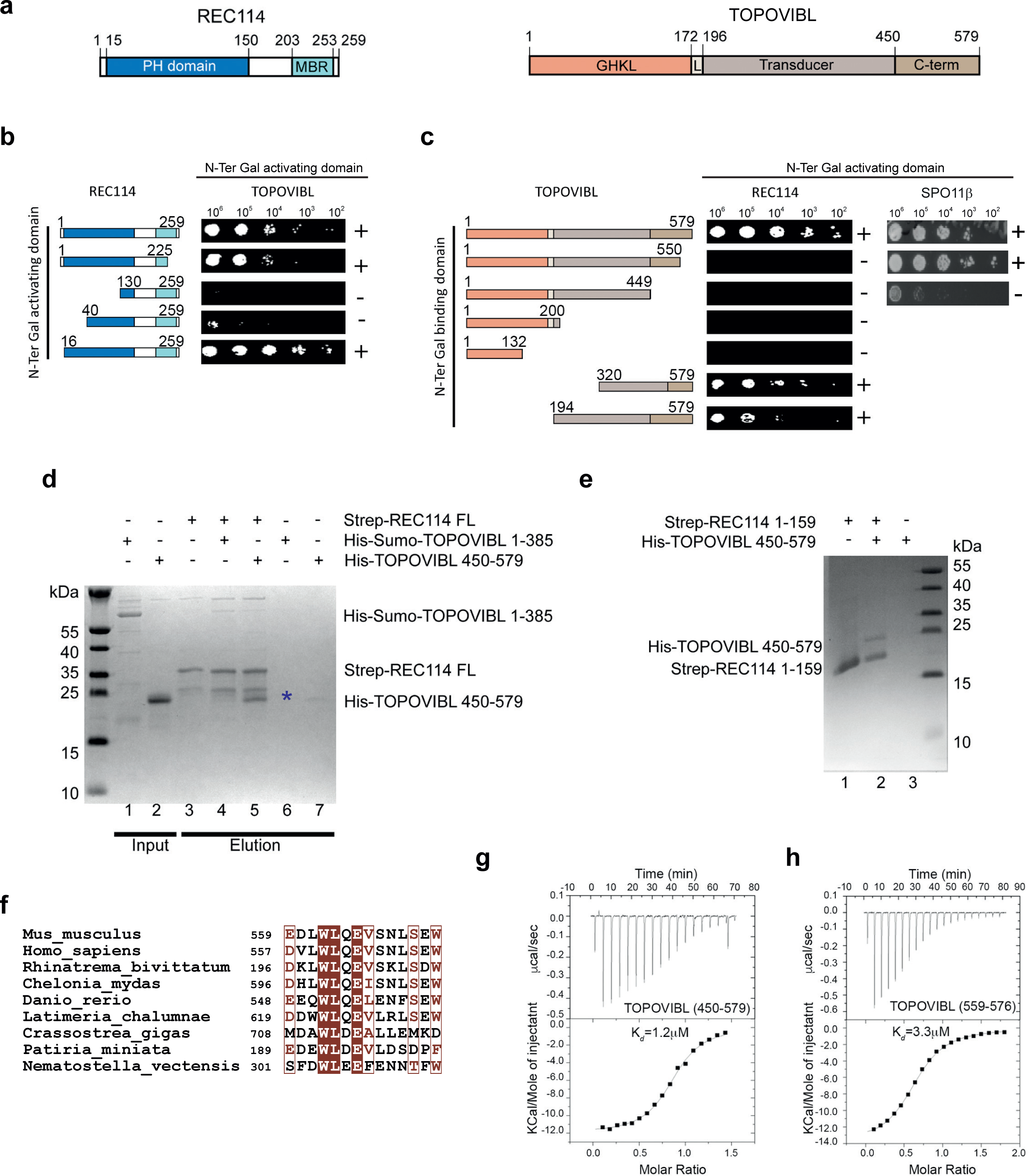
The C-terminal region of TOPOVIBL interacts with the N-terminal domain of REC114. **a.** Schematic representation of the domain structure of mouse REC114 (left) and TOPOVIBL (right). PH: Pleckstrin homology domain, MBR: MEI4 Binding Region, GHKL: Gyrase, HSP90, Histidine Kinase, MutL domain, L: linker, C-term: C-terminal. **b.** Yeast two-hybrid assays showed that REC114 N-terminal domain is required for the interaction with TOPOVIBL. Growth (+ or -) was assayed on medium lacking leucine, tryptophane and histidine and with 5mM 3-amino-1,2,4-triazole. **c.** Yeast two-hybrid assays indicated that TOPOVIBL last 29 residues are required for the interaction with REC114 but not with SPO11β. Growth (+ or -) was assayed on medium lacking leucine, tryptophane and histidine. **d.** Strep-tag pull-down experiments with REC114 and TOPOVIBL. The indicated TOPOVIBL domains were first purified on Ni^2+^ columns (lanes 1,2) and then loaded onto Strep-Tactin columns with or without previously bound full-length Strep-REC114. TOPOVIBL 1-385 is not retained by REC114 (lane 4), whereas TOPOVIBL C-terminal region (450-579) is sufficient for the interaction with REC114 (blue star, lane 5). **e.** The REC114 PH domain (1-159) is sufficient for the interaction with TOPOVIBL. Ni^2+^ column-purified TOPOVIBL 450-579 is co-eluted with Strep-REC114 1-159 (PH domain, lane 2), but is not retained by free Strep-Tactin resin (lane 3). **f.** Sequence alignment of the 14-aa conserved motif at TOPOVIBL C-terminus in metazoans Brown letters: equivalent amino acids; white letters: identical amino acids. **g.** ITC measurement of the interaction affinity between REC114 15-159 and TOPOVIBL 450-579. **h.** ITC measurement of the interaction affinity between REC114 15-159 and TOPOVIBL 559-576.

The structure prediction analysis and AlphaFold model (AF-J3QMY9-F1) suggested that TOPOVIBL CTD did not to contain any known globular domain and was partially intrinsically disordered. However, it indicated the presence of a putative helix at its C-terminus. Interestingly, within the last C-terminal residues, we detected high conservation among metazoan species (Fig. 1f). As our yeast two-hybrid assays suggested that the corresponding C-terminal 29 residues of TOPOVIBL (residues 551 to 579) were important for the interaction with REC114 (Fig. 1c), we hypothesized that this predicted conserved helix (residues 559-572) might represent the REC114 binding region. Using isothermal titration calorimetry (ITC), we showed that REC114 PH domain bound to TOPOVIBL CTD (450-579) with a dissociation constant (Kd) of 1.2 μM (Fig. 1g). Moreover, the interaction between a TOPOVIBL peptide spanning residues 559-576 and REC114 PH domain was in the same range (Kd: 3.3 μM) (Fig. 1h). These results demonstrate that this highly conserved motif in TOPOVIBL CTD (559-576 aa) is sufficient for interaction with REC114.

We then determined the crystal structure of the complex formed by the REC114 PH domain (residues 15-159) and the TOPOVIBL 559-576 peptide by X-ray crystallography (2.3Å resolution, Rfree of 24.9%, and R-factor of 22.6%) (Extended data Table 1, Extended data Fig. 1b). The structure of the TOPOVIBL-bound REC114 PH domain (two perpendicular antiparallel β-sheets followed by a C-terminal helix) was essentially the same as in its unbound form ^23^. The TOPOVIBL C-terminal peptide folds into a single helix that interacts with the REC114 PH domain β-sheet formed of strands β1, β2 and β6-β8, burying 731 Å2 of surface area (Figs. 2a and 2b). The interaction surface on REC114 is formed of highly conserved surface residues (Fig. 2c). In the N-terminal part of the TOPOVIBL peptide, L561, W562 and V566 pack against a hydrophobic surface of REC114 (Fig. 2d). Specifically, W562 is located in a hydrophobic pocked formed by aliphatic side chains of K95, V97, L104 and M115 of REC114. In the central part of the TOPOVIBL helix, L569 binds to a hydrophobic groove in REC114 formed by V97, R99, C102, L104, and R117 (Fig. 2e). The well-conserved R99 and R117 residues form several hydrogen bonds with main-chain carbonyls in the TOPOVIBL helix, and R117 forms a salt bridge with E571. The C-terminal part of the TOPOVIBL peptide forms a 310 helix where W572 inserts into another hydrophobic pocket of REC114 made of R24, V53, C102 and R117 (Fig. 2f). W572 also forms cation-π interactions by packing against the ammonium groups of the two arginine residues and also main-chain hydrogen bonds with R117 and Q119. Finally, L573 packs against a hydrophobic surface around M110 (Fig. 2f).

**Fig. 2.**
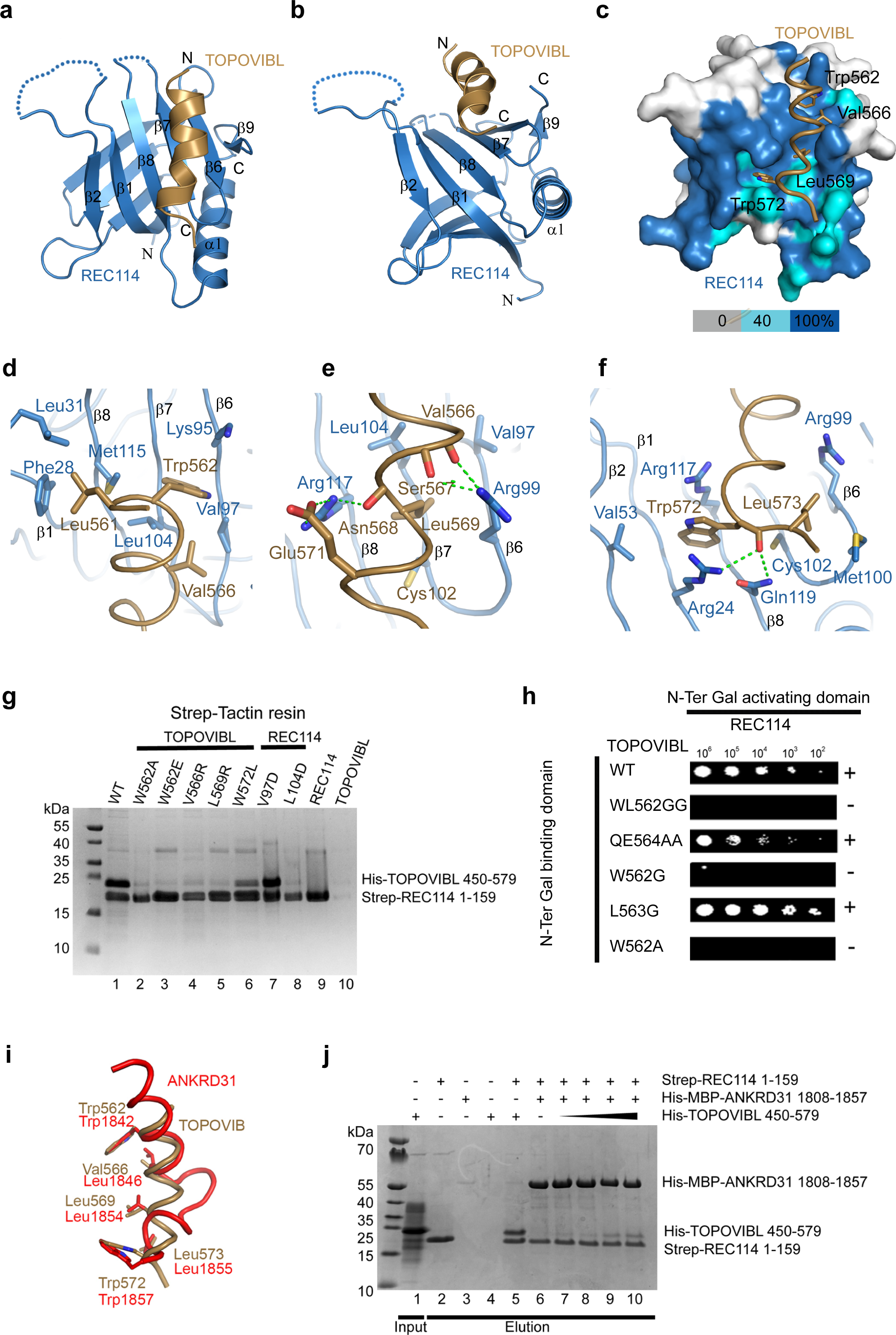
Structure of the REC114-TOPOVIBL complex. **a.** Ribbon representation of the overall structure of the REC114-TOPOVIBL complex. The REC114 PH domain is in blue, and the TOPOVIBL peptide in brown. Alpha helices (α) and beta sheets (β) are labelled. **b.** REC114-TOPOVIBL structure rotated 60° around the horizontal axis, compared to panel a. **c.** Surface representation of REC114 to highlight the conserved surface residues. Sequence conservation is represented from grey to blue according to the colour scale bar below. TOPOVIBL is shown as a cartoon and the key interacting residues as sticks. **d.** Details of the interaction between the N-terminal part of the TOPOVIBL helix (brown) and REC114 (blue). TOPOVIBL W562 inserts into a hydrophobic pocket on the β-sheet formed by strands β1 and β6-β8. **e.** The central part the TOPOVIBL helix (brown) forms several hydrogen bonds (green dotted lines) with conserved REC114 residues. L569 interacts with another hydrophobic cavity formed by β6-β8. **f.** The C-terminal W572 residue of TOPOVIBL forms hydrophobic and charged interactions with REC114. **g.** SDS-PAGE analysis of Strep-tag pull-down experiments with the indicated TOPOVIBL and REC114 proteins. Wild-type (WT) and mutants of His-TOPOVIBL were first purified on Ni^2+^ column and loaded on Strep-REC114 bound Strep-Tactin resin. **h.** Essential role of TOPOVIBL W562 in the interaction with REC114 shown by yeast two-hybrid assays. Growth (+ or -) was assayed on medium lacking leucine, tryptophane and histidine and with 5mM 3-amino-1,2,4-triazole. **i.** The key TOPOVIBL and ANKRD31 residues that interact with the REC114 β-sheet (β1, β6-β8) are in similar positions. **j.** ANKRD31-TOPOVIBL competition assay. TOPOVIBL 450-579 was purified on a Ni^2+^ column (lane 1). Addition of 1 mg of TOPOVIBL 450-579 to Strep-REC114 1-159 is sufficient to form an apparently stoichiometric complex (lane 5). When REC114 is bound to ANKRD31 1808-1857 (lane 6), addition of increasing amount of TOPOVIBL 450-579 (1 to 20mg) is not sufficient to form TOPOVIBL-REC114 complexes (lanes 7-10).

Most of the TOPOVIBL residues involved in the interaction with REC114 are highly conserved across species (Fig. 1f). We showed the interaction conservation by modelling the complex from different vertebrate species using AlphaFold ^27^ (Extended data Fig. 1c). To identify the TOPOVIBL residues required for the interaction with REC114, we mutated several candidate residues (W562, V566, L569 and W572) based on structure and conservation. In pull-down assays with the Strep-tagged REC114 PH domain, mutations in W562, V566 and L569 reduced the interaction (Fig. 2g, lanes 2-5), and the W572L mutation reduced the interaction compared with the wild-type residue (Fig. 2g, lane 6). We also mutated two hydrophobic residues of the REC114 β-sheet (V97D, L104D). V97D had little effect, whereas L104D abolished the interaction with TOPOVIBL (Fig. 2g, lanes 7,8). We obtained similar results in pull-down experiments using His-tagged TOPOVIBL (Extended data Fig. 1d). We confirmed TOPOVIBL W562 role in the interaction by ITC measurement, because the peptide containing the W562A mutation did not bind to REC114 PH domain (Extended data Fig. 1e). Gel filtration indicated that W562A in TOPOVIBL 450-579 did not disrupt the protein folding (Extended data Fig. 1f). Moreover, in yeast two-hybrid assays using full-length proteins we showed that W562A and W562G led to loss of any detectable interaction (Fig. 2h). We conclude that the TOPOVIBL W562A mutation specifically decreases the interaction with REC114, and we selected this mutation for the *in vivo* studies (see below).

Mouse REC114 directly interacts also with MEI4 ^23^ and ANKRD31 ^25^. The REC114/MEI4 interaction does not involve the REC114 PH domain but the C-terminal domain (203-254) ^23^. In agreement, we could show by size exclusion chromatography the simultaneous interaction of REC114 with MEI4 and TOPOVIBL (Extended data Fig. 1g). Conversely, the REC114/ANKRD31 interaction involves the REC114 PH domain. The crystal structure of the ANKRD31-REC114 complex ^25^ shows that the ANKRD31 interacting fragment (45 residues) is significantly longer than that of TOPOVIBL and covers a larger surface on REC114 (∼1800 Å2) packing against both of its β-sheets (Extended data Figs. 1h and 1i). The C-terminal part of the ANKRD31 peptide forms two helices and packs against the same surface as TOPOVIBL, and both peptides interact with equivalent REC114 residues (Fig. 2i). This structural comparison indicates mutually exclusive binding of these two proteins to REC114, possibly with higher affinity for ANKRD31. Mutation of REC114 L104 reduced the binding to both ANKRD31 ^25^ and to TOPOVIBL (Fig. 2g, lane 8). Similarly, the ANKRD31 W1842A mutation disrupted the interaction with REC114 ^25^, as did the corresponding W562A mutation in TOPOVIBL. As in our hands SUMO or MBP fusion ANKRD31 1808-1857 proteins aggregated unless bound to REC114, we could not determine its Kd for REC114. To confirm the mutually exclusive nature of the ANKRD31/TOPOVIBL binding to REC114, we performed ITC measurements of the interaction of REC114 with TOPOVIBL 450-579 in the absence or presence of saturating amounts of ANKRD31 1808-1857. While TOPOVIBL 450-579 normally interacted with REC114 with a Kd of 1.2 μM, we did not observe any binding when REC114 was pre-saturated with ANKRD31 1808-1857 (Extended data Fig. 1j). In pull-down assays, the interaction with TOPOVIBL 450-479 was significantly reduced when REC114 was first bound to ANKRD31 1808-1857 (Fig. 2j). These results indicate a mutually exclusive binding of TOPOVIBL and ANKRD31 to REC114 *in vitro*.

### *Top6bl* mutations lead to a DSB activity reduction in oocytes

To evaluate the biological significance of the TOPOVIBL-REC114 interaction, we generated two mutant alleles of mouse *Top6bl*: i) *Top6bl^W562A^* harbouring the point mutation W562A that weakens the interaction between TOPOVIBL and REC114; and ii) *Top6bl^Δ17Ct^* harbouring a truncation of TOPOVIBL C-terminal helix that interacts with REC114 (Extended data Fig. 2 for details on mutant generation).

We first investigated DSB formation during meiotic prophase of oocytes from embryonic ovaries (16 days post-coitum, dpc), when leptonema and zygonema are predominant ^28^. To follow DSB formation, we analysed the phosphorylated form of H2AX (γH2AX) that appears at chromatin domains around DSB sites upon DSB formation ^29^. In wild-type oocytes, γH2AX was present over large chromatin domains at leptonema and zygonema and mostly disappeared at pachynema (Fig. 3a). In both *Top6bl* mutants, γH2AX intensity was strongly reduced at leptonema and zygonema: by 5.2- and 2.5-fold, respectively, in *Top6bl^W562A/W562A^,* and by 8-fold and 7.6-fold, respectively, in *Top6bl^Δ17Ct/Δ17Ct^* oocytes, compared with wild-type oocytes (Figs. 3a and 3b). To determine whether this could be due to delayed DSB formation, we monitored γH2AX levels at a later developmental stage (18dpc) in *Top6bl^Δ17Ct/Δ17Ct^* mice and found a reduction by 7.8-fold (Extended data Fig. 3a). These findings suggest that the two *Top6bl* mutations lead to a decrease of DSB activity.

**Fig. 3.**
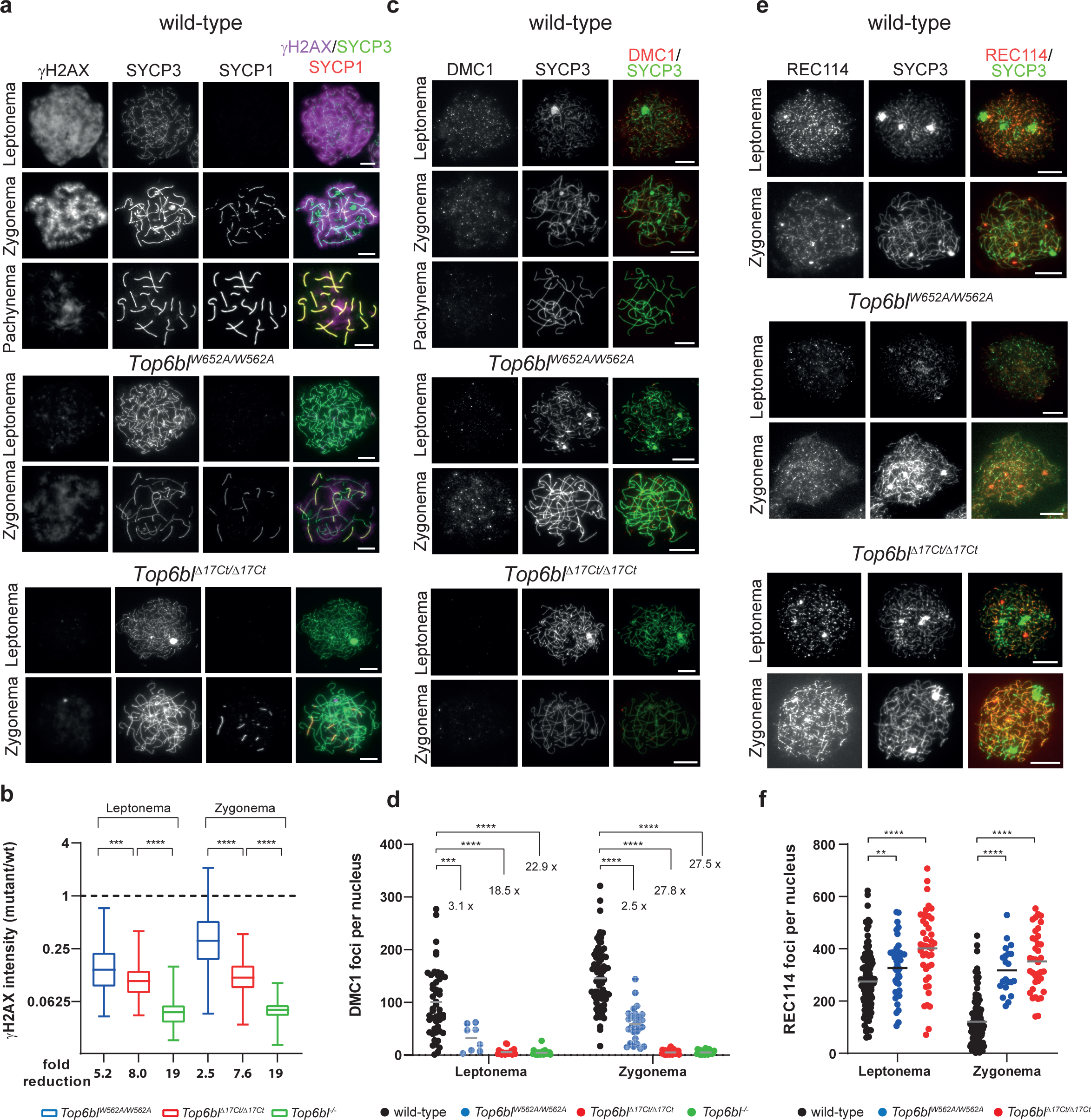
In *Top6bl^W562A/W562A^* and *Top6bl ^Δ17Ct/Δ17Ct^* mice, meiotic DSB activity is decreased in oocytes. **a.** Immunostaining of γH2AX, SYCP3 and SYCP1 in oocytes from 16 dpc wild-type (+/+), *Top6bl^W562A/W562A^,* and *Top6bl ^Δ17Ct/Δ17Ct^* ovaries. Scale bar, 10μm. **b.** Quantification of γH2AX signal intensity in leptotene and zygotene (or zygotene-like) nuclei of oocytes from 16 dpc wild-type (+/+ or *Top6bl ^+/Δ17Ct^*), *Top6bl^W562A/W562A^*, *Top6bl ^Δ17Ct/Δ17Ct^*, and *Tob6bl^-/-^* mice (n=5, 2, 2, and 1 mouse/genotype). Number of nuclei at leptonema: 126, 123, 73, and 23; number of nuclei at zygonema: 241, 132, 64, and 33 for each genotype. Ratios (mean ± SD) of the integrated intensity between the mean values in mutant nuclei and in wild-type nuclei are plotted. P values were determined using the two-tailed unpaired Mann-Whitney test. The fold reduction is the ratio of the wild-type to mutant mean values. **c.** Immunostaining of DMC1 and SYCP3 in oocytes from 16 dpc or 17 dpc wild-type (+/+), *Top6bl^W562A/W562A^* and *Top6bl ^Δ17Ct/Δ17Ct^* ovaries. Scale bar, 10μm. **d.** Quantification of DMC1 foci. DMC1 axis-associated foci were counted in leptotene and zygotene nuclei of oocytes from 16 and 17 dpc wild-type (+/+ or *Top6bl ^+/W562A^*), *Top6bl^W562A/W562A^*, *Top6bl ^Δ17Ct/Δ17Ct^*, and *Top6bl^-/-^* mice. Number of nuclei at leptonema: 50, 8, 22, and 28; number of nuclei at zygonema: 79, 28, 29, and 27 for each genotype, respectively. Grey bars show the mean values. P values were determined using the two-tailed unpaired Mann-Whitney test. The fold difference relative to wild-type is shown. **e.** Immunostaining of REC114 and SYCP3 in oocytes from 16dpc wild-type (+/+), *Top6bl^W562A/W562A^*, and *Top6bl ^Δ17Ct/Δ17Ct^* ovaries. Scale bar, 10μm. **f.** Quantification of axis-associated REC114 foci in leptotene and zygotene oocytes from 15 dpc wild-type (+/+), *Top6bl^W562A/W562A^*, and *Top6bl ^Δ17Ct/Δ17Ct^* mice (n=1 mouse/genotype). Number of nuclei at leptonema: 51 and 43; number of nuclei at zygonema: 58 and 39 in wild-type and *Top6bl ^Δ17Ct/Δ17Ct^* oocytes, respectively. Grey bars show the mean values. P values were determined using the two-tailed unpaired Mann-Whitney test.

We confirmed this hypothesis by quantifying DSB repair through the detection of DMC1 and RPA2. DMC1 binds to resected DSB ends and catalyses homologous strand exchange for DSB repair. RPA is recruited to resected DSB ends before and also after strand exchange during second end capture for repair ^2, 30^. In wild-type oocytes, we detected DMC1 and RPA foci that colocalized with the chromosome axis at leptonema and zygonema. The number of DMC1 foci was decreased in both *Top6bl* mutants at leptonema (3.1-fold in *Top6bl^W562A/W562A^*and 18.5-fold in *Top6bl^Δ17Ct/Δ17Ct^* mice) (Figs. 3c and 3d). DMC1 level in *Top6bl^Δ17Ct/Δ17Ct^* mice was close to the background level because DMC1 foci at leptonema and zygonema were reduced by 22.9-fold in *Top6bl^-/-^*mice where DSB formation is abolished (Fig. 3d). The reduction of DMC1 foci was also observed at zygonema (Fig. 3d). We obtained similar results for RPA2 foci (Extended data Figs. 3b and 3c). In wild-type oocytes, DSB repair promotes interactions between homologues that are stabilized by the recruitment and assembly of several proteins, including SYCP1 and SYCP3, to form the synaptonemal complex (Fig. 3a)^31^. In both *Top6bl^W562A/W562A^* and *Top6bl^Δ17Ct/Δ17Ct^* mice, we observed only short stretches of synapsis and very few nuclei with full synapses. In 16dpc wild-type ovaries, 42.5%, 32.5% and 20% of oocytes were in leptonema, zygonema and pachynema (n=315), respectively, compared with 61.9%, 37.7% and 0.4% (n = 496) in *Top6bl^W562A/W562A^,* and 57.6%, 42.4% and 0% (n=151) in *Top6bl^Δ17Ct/Δ17Ct^* ovaries. Overall, these results are consistent with a reduced DSB activity that affects synapsis formation between homologues.

As formation of meiotic DSBs depends on the pre-DSB proteins IHO1, REC114, MEI4, ANKRD31 and MEI1 ^21, 23–26, 32^, and because the two *Top6bl* mutations disrupt the interaction with REC114, we assessed REC114 cytological localization in *Top6bl^W562A/W562A^* and *Top6bl^Δ17Ct/Δ17Ct^* mice. In wild-type oocytes, REC114, IHO1, MEI4 and ANKRD31 form several hundred foci on chromosome axes at leptonema and they progressively disappear as DSBs form ^22–26^. The number of REC114 axis-associated foci was significantly higher in *Top6bl^W562A/W562A^* and particularly in *Top6bl^Δ17Ct/Δ17Ct^* mice at leptonema and especially at zygonema compared with wild-type oocytes (Figs. 3e and 3f). This higher number of foci is consistent with the decreased DSB activity because REC114 foci are displaced from chromosome axes upon DSB formation ^23^. Indeed, in spermatocytes, in the absence of meiotic DSB activity, such as in *Spo11^-/-^* mice, REC114 (and MEI4) accumulate on unsynapsed axes ^22, 23^. We verified that this accumulation of REC114 at zygonema in the absence of meiotic DSB activity also applies to oocytes, by analyzing *Top6bl^-/-^*mutant (Extended data Fig. 3f). ^23^We obtained similar results by analysing MEI4 and ANKRD31 foci in *Top6bl^W562A/W562A^* and *Top6bl^Δ17Ct/Δ17Ct^* oocytes (Extended data Figs. 3d, 3e, 4a and 4b). Overall, these analyses demonstrate that loading of REC114, MEI4 and ANKRD31 is not altered by the *Top6bl^W562A^* and *Top6bl^Δ17Ct^* mutations.

**Fig. 4.**
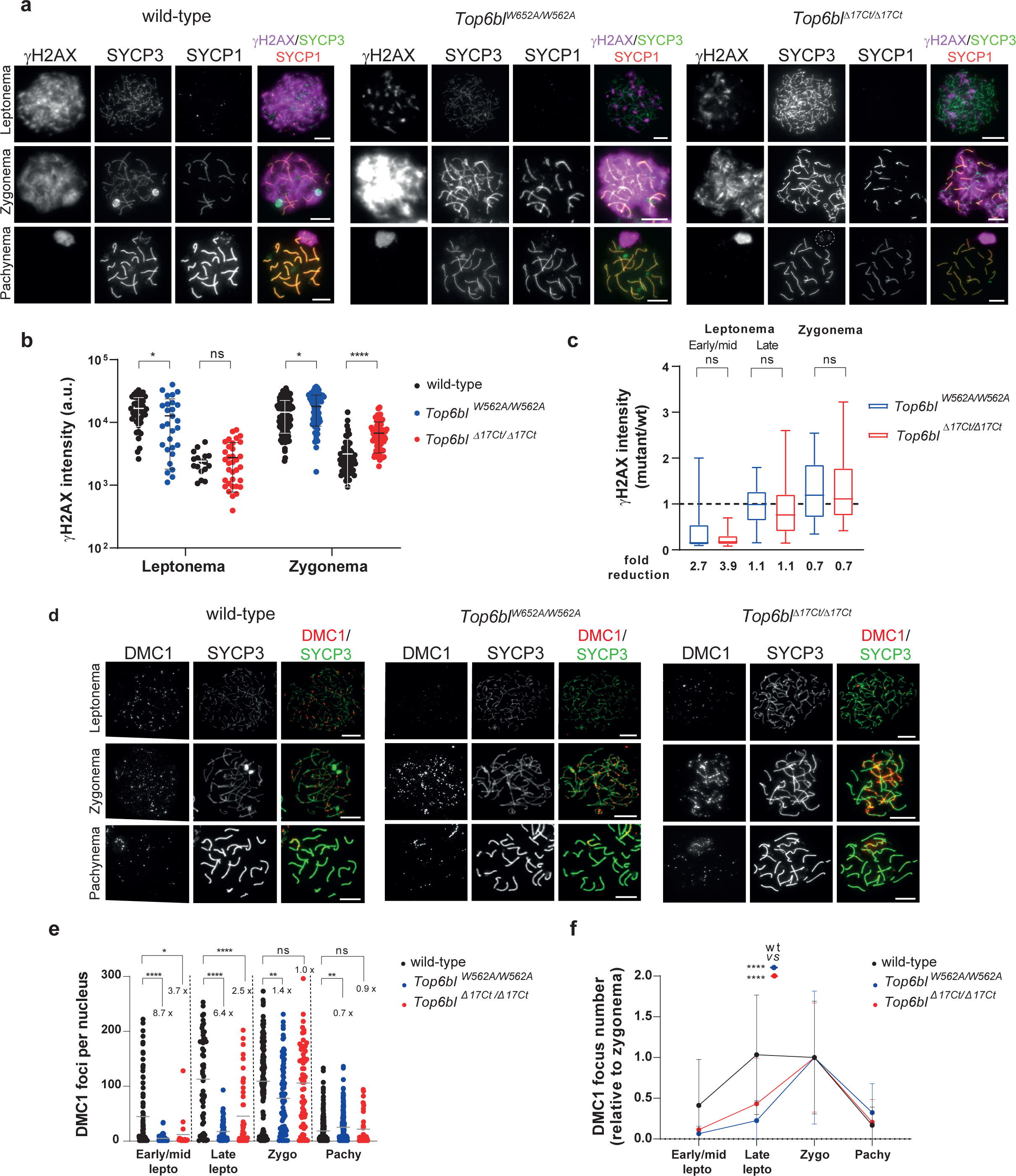
Efficient but delayed DSB formation in *Top6bl* mutant spermatocytes. **a.** Immunostaining of γH2AX, SYCP3 and SYCP1 in spermatocytes from 12 dpp wild-type (+/+), *Top6bl^W562A/W562A^*, and *Top6bl ^Δ17Ct/Δ17Ct^* mice. A white dotted circle highlights the unsynapsed X and Y chromosomes at pachynema (SYCP3 staining) in *Top6bl ^Δ17Ct/Δ17Ct^* mice. Scale bar, 10μm. **b.** Quantification of γH2AX intensity (mean ± SD; a.u., arbitrary units) in leptotene and zygotene spermatocytes from 12 dpp wild-type, *Top6bl^W562A/W562A^*, and *Top6bl ^Δ17Ct/Δ17Ct^* mice (n=1 mouse per genotype). Each mutant had a wild-type control performed in parallel. Number of nuclei in the 8 groups plotted on the x axis: 44, 28, 17, and 34 at leptonema; 132, 61, 77, and 66 at zygonema. P values were determined using the two-tailed unpaired Mann-Whitney test. **c.** Quantification of γH2AX intensity in *Top6bl^W562A/W562A^* and *Top6bl ^Δ17Ct/Δ17Ct^* spermatocytes relative to wild-type at early/mid, late leptotene, and zygotene. Stages were defined as described in Methods and shown in Extended data Fig. 6a. Number of nuclei: 19, 201 at early/mid, 33, 290 at late leptotene, and 88, 333 at zygotene in *Top6bl^W562A/W562A^* and *Top6bl ^Δ17Ct/Δ17Ct^* mice, respectively. The fold reduction is the ratio of the mean values in wild-type and mutant samples. P values were determined using the two-tailed unpaired Mann-Whitney test. **d.** Immunostaining of DMC1 and SYCP3 in spermatocytes from 14 dpp wild-type (+/+), *Top6bl^W562A/W562A^*, and *Top6bl ^Δ17Ct/Δ17Ct^* mice. Scale bar, 10μm. **e.** Quantification of DMC1 foci. Axis-associated DMC1 foci were counted in leptotene (early/mid and late), zygotene, and pachytene nuclei of spermatocytes from wild-type (+/+), *Top6bl^W562A/W562A^*, and *Top6bl ^Δ17Ct/Δ17Ct^* mice (n=3 wild-type, and n=2 mice per mutant genotype). Number of nuclei: 95, 50 and 18 at early/mid leptotene, 55, 63 and 45 at late leptotene, 120, 92 and 66 at zygotene, and 261, 207 and 36 at pachytene for each genotype, respectively. Grey bars show the mean values. P values were determined using the two-tailed unpaired Mann-Whitney test. **f.** Variation of DMC1 focus number during prophase. For each genotype (wild-type, *Top6bl^W562A/W562A^*, and *Top6bl ^Δ17Ct/Δ17Ct^*), the number of DMC1 foci at the indicated stages was normalized to the mean number at zygonema (set to 1). Mean values ± SD are shown. Statistical significance between wild-type and each mutant (blue *Top6bl^W562A/W562A^*; red *Top6bl ^Δ17Ct/Δ17Ct^*) was tested at late leptonema using the two-tailed unpaired Mann-Whitney test.

These meiotic prophase defects affected oogenesis: follicle number was strongly reduced in ovaries of *Top6bl^W562A/W562A^*and particularly *Top6bl^Δ17Ct/Δ17Ct^* mice (Extended data Figs. 5a, 5b and 5d). Indeed, *Top6bl^Δ17Ct/Δ17Ct^* mice were sterile, whereas *Top6bl^W562A/W562A^* mice were sub-fertile (Extended data Fig. 5c). We conclude that the TOPOVIBL-REC114 interaction is essential for meiotic DSB formation in oocytes.

**Fig. 5.**
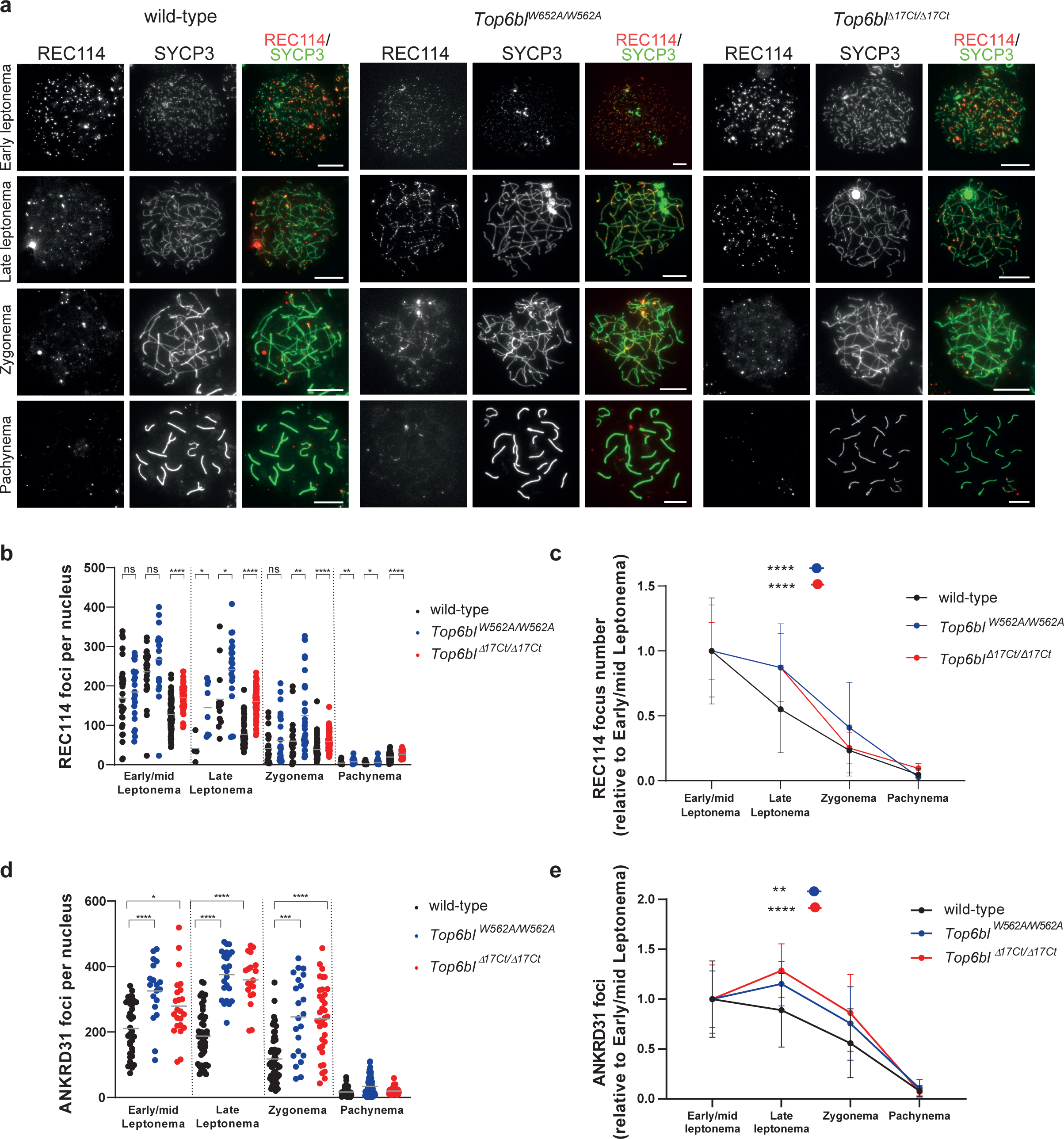
Efficient REC114 and ANKRD31 loading in *Top6bl* mutant spermatocytes. **a.** Immunostaining of REC114 and SYCP3 in spermatocytes from 12-14dpp wild-type (+/+), *Top6bl^W562A/W562A^*, and *Top6bl ^Δ17Ct/Δ17Ct^* mice. Scale bar, 10μm. **b.** Quantification of axis-associated REC114 foci in early/mid and late leptotene, zygotene and pachytene spermatocytes from 12-14dpp wild-type (+/+ or *Top6bl ^+/Δ17Ct^* : n=4), *Top6bl^W562A/W562A^* (n=2), and *Top6bl^Δ17Ct /Δ17Ct^* (n=2) mice. Each mutant is compared with a wild-type control in parallel to control for variations between experiments. Results from *Top6bl^W562A/W562A^* are from two independent experiments. Number of nuclei: wild-type: 31 early/mid L, 4 late L, 26 Z, 51 P; *Top6bl^W562A/W562A^*: 22 early/mid L, 8 late L, 26 Z, 24 P; wild-type 28 early/mid L, 12 late L, 24 Z, 46 P; *Top6bl^W562A/W562A^* 18 early/mid L, 21 late L, 32 Z, 38 P; wild-type: 84 early/mid L, 59 late L, 84 Z, 75 P; *Top6bl ^Δ17Ct/Δ17Ct^* : 76 early/mid L, 68 late L, 79 Z, 61 P. Grey bars show the mean values. P values were determined using the two-tailed unpaired Mann-Whitney test. **c.** Variation of REC114 foci during prophase. The number of foci at early/mid leptonema, late leptonema, zygonema and pachynema relative to the mean number at early/mid leptonema was plotted for wild-type, *Top6bl^W562A/W562A^*, and *Top6bl ^Δ17Ct/Δ17Ct^* mice. The mean values ± SD are shown. Statistical significance between wild-type and each mutant (*Top6bl^W562A/W562A^* and *Top6bl ^Δ17Ct/Δ17Ct^*) was tested at late leptonema using the two-tailed unpaired Mann-Whitney test. **d.** Quantification of axis-associated ANKRD31 foci in early/mid, late leptotene, zygotene and pachytene spermatocytes from 12-14 dpp wild-type (+/+ or *Top6bl ^+/Δ17Ct^* : n=2), *Top6bl^W562A/W562A^* (n=1), and *Top6bl^Δ17Ct /Δ17Ct^* (n=1) mice. Mean number of nuclei per genotype at early/mid leptotene (37, 19, and 23), late leptotene (51, 23, and 19), zygotene (61, 22, and 35), and pachytene (100, 61, and 31). Grey bars show the mean values. P values were determined using the two-tailed unpaired Mann-Whitney test. **e.** Variation of ANKRD31 foci during prophase. The number of foci at early/mid leptonema, late leptonema, zygonema, and pachynema relative to the mean number at early/mid leptonema (set at 1) was plotted for wild-type, *Top6bl^W562A/W562A^*, and *Top6bl ^Δ17Ct/Δ17Ct^* mice. The mean values ± SD are shown. Statistical significance between wild-type and each mutant (*Top6bl^W562A/W562A^* and *Top6bl ^Δ17Ct/Δ17Ct^*) was tested at late leptonema using the two-tailed unpaired Mann-Whitney test.

### In *Top6bl* spermatocytes, DSBs are delayed genome-wide and decreased in sub-telomeric regions

Unlike female meiosis, DSB activity was efficient in spermatocytes from both *Top6bl* mutants, as indicated by the detection and quantification of γH2AX at late leptonema and zygonema (Figs. 4a-4c). Conversely, at early/mid leptonema, γH2AX levels were lower in *Top6bl^W562A/W562A^* and *Top6bl^Δ17Ct/Δ17Ct^* than wild-type spermatocytes, suggesting a delay in DSB formation (Fig. 4c).

DMC1 and RPA2 foci also appeared later in both mutants compared with wild-type spermatocytes (Figs. 4d-4f and Extended data Figs. 6b-6d). The number of DMC1 and RPA2 foci was lower in *Top6bl^W562A/W562A^* and *Top6bl^Δ17Ct/Δ17Ct^* than in wild-type spermatocytes, particularly at early/mid leptonema (8.7- and 3.7-fold reduction of DMC1 foci and 11.7- and 7.4-fold reduction of RPA2 foci in *Top6bl^W562A/W562A^* and *Top6bl^Δ17Ct/Δ17Ct^*, respectively, compared with wild-type spermatocytes). Conversely, at zygonema and pachynema, the number of DMC1 and RPA2 foci was similar in wild-type and mutant spermatocytes (Fig. 4e and Extended data Fig. 6b). These findings suggest an efficient but delayed formation of DMC1 and RPA2 foci, and efficient DSB repair in *Top6bl^W562A/W562A^* and *Top6bl^Δ17Ct/Δ17Ct^* spermatocytes.

The localization of the DSB axis-associated proteins REC114, ANKRD31 and MEI4 at early/mid leptonema and the number of foci were similar or higher than in wild-type spermatocytes (Figs. 5a, 5b, 5d and Extended data Figs. 7a-7c). The number of foci gradually decreased from early/mid leptonema in wild-type spermatocytes, but only after late leptonema (ANKRD31) or after early/mid leptonema (REC114) and with a slower kinetic in the *Top6bl* mutants (Figs. 5c and 5e). These kinetic alterations are compatible with the observed delayed DSB activity because these axis-associated proteins disassemble from the axis upon DSB formation ^23, 25, 26^.

To directly evaluate DSB activity and to map DSB sites, we monitored DMC1 enrichment by DMC1 chromatin-immunoprecipitation (ChIP), followed by ssDNA enrichment (DMC1-Single Strand DNA Sequencing, SSDS) ^33^ in *Top6bl^Δ17Ct/Δ17Ct^* mice. We identified 16780 DSB hotspots in *Top6bl^Δ17Ct/Δ17Ct^*, among which 13261 (79%) overlapped with wild-type hotspots (Fig. 6a). This indicated efficient PRDM9-dependent DSB localization in the mutant, as confirmed also by the absence of significant signal at PRDM9-independent hotspots ^34^( Extended data Fig. 8). The hotspot intensity was similar in *Top6bl^Δ17Ct/Δ17Ct^* and wild-type samples for most hotspots (Fig. 6b). 3519 peaks were however specific to *Top6bl^Δ17Ct/Δ17Ct^* mice (Fig. 6a). At these weak hotspots, we however detected a weak enrichment for DMC1-SSDS in wild-type samples, ∼1.7-fold lower than in *Top6bl^Δ17Ct/Δ17Ct^* mice (Extended data Fig. 9a). PRDM9^B6^-specific H3K4me3 enrichment, evaluated in wild-type mice, also indicated a weak PRDM9 activity at these sites (Extended data Fig. 9a). We asked if we could also detect any increase of intensity in *Top6bl^Δ17Ct/Δ17Ct^* among the subset of the weakest hotspots identified in wild-type, but this was not the case (bin 1 from Extended data Fig. 9a). A differential analysis of DMC1-SSDS signal intensity at common hotspots (using the DESeq2 R package, see Methods) showed that in *Top6bl^Δ17Ct/Δ17Ct^* mice, the signal was decreased (by 1.2- to 24-fold) in 12% of hotspots, and increased (by 1.45- to 3.3-fold) in 2% of hotspots (Fig. 6c). Visual analysis highlighted that in *Top6bl^Δ17Ct/Δ17Ct^* mice, the DMC1-SSDS signal was decreased at hotspots located near the q-arm telomeres, the telomeres distal to centromeres (Fig. 6d and Extended data Figs. 10 and 11). We quantified this sub-telomeric phenotype by different approaches. First, by quantifying the ratio of the mutant/wild-type signal along the chromosome arms, we observed a ≥2-fold decrease in the last few megabases proximal to the q-arm telomeres (Fig. 6e). Second, in the ten hotspots closest to the q-arm telomeres, located within 4Mb of the telomere, DMC1-SSDS signal intensity at most hotspots for each chromosome was lower in *Top6bl^Δ17Ct/Δ17Ct^* than wild-type samples (blue for autosomes, red for the X chromosome) (Fig. 6b). Third, quantification of this effect at each chromosome showed a reduced DMC1-SSDS signal within the ten q-arm telomeric adjacent hotspots in *Top6bl^Δ17Ct/Δ17Ct^* mice at most chromosomes (Fig. 6f and Extended data Fig. 9c). Fourth, using DESeq, we evaluated the effect of the distance from telomeres, and found that the effect was most pronounced at hotspots located within 3Mb from telomeres and decreased when we tested larger intervals (Fig. 6g and Extended data Fig. 9d). In this analysis, the statistical significance also depended on the hotspot number in the tested region for each chromosome (see, for instance, chromosome 17, Extended data Fig. 9e). We conclude that in *Top6bl^Δ17Ct/Δ17Ct^* mice, the DMC1-SSDS signal is specifically reduced in the 3Mb sub-telomeric region of the q-arm of most chromosomes. This finding could indicate a DSB decrease or higher DMC1 turnover. This region-specific alteration may not affect homologous interactions between autosomes because these interactions should be ensured by DSB sites along the chromosome arms. However, on the X and Y chromosomes, which depend on recombination in the PAR and which is located proximal to the telomere, the decreased DSB activity in this region in *Top6bl^Δ17Ct/Δ17Ct^* mice (Fig. 6d) could influence their homologous interaction.

**Fig. 6.**
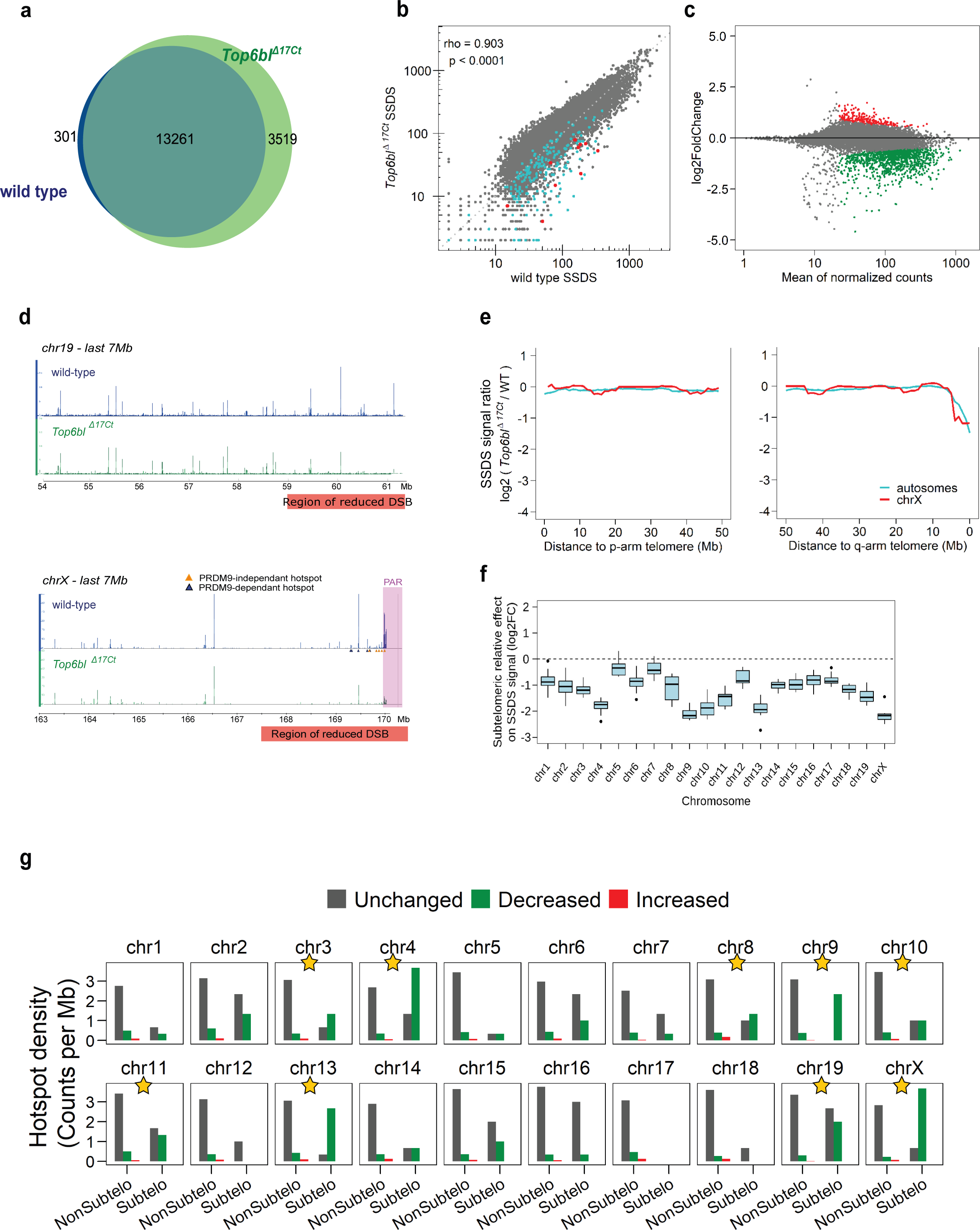
Proper DMC1-SSDS signal localisation and intensity, excepted for q-arm sub-telomeric regions in *Top6bl ^Δ17Ct/Δ17Ct^* spermatocytes. **a.** In *Top6bl^Δ17Ct/Δ17Ct^* spermatocytes, almost all wild-type hotspots and 20% of new hotspots are detected. The Venn diagram represents the DMC1-SSDS peaks in wild-type and *Top6bl^Δ17Ct/Δ17Ct^* mice. **b.** DMC1-SSDS signal correlation between wild-type and *Top6bl^Δ17Ct/Δ17Ct^* mice at wild-type hotspots. The Spearman rho and associated p-value are shown. Ten telomere-proximal hotspots are highlighted for each autosome (blue) and for the X chromosome (red). In the PAR, only part of the DMC1-SSDS signal, which covers a large domain (see panel d), is included within hotspots. **c.** MA plot of the DMC1-SSDS signal in *Top6bl^Δ17Ct/Δ17Ct^* compared with wild-type samples. The log2 of the ratios was calculated using DESeq2 for differential analysis and after log fold change shrinkage. Hotspot with significantly increased or decreased signal are highlighted in red (n=214) and green (n=1098), respectively (adjusted p-value<0.1). Unchanged hotspots are in grey (n=8043). The mean normalized count corresponds to the baseMean value from the DESeq2 analysis. **d.** DSB maps of the 7Mb telomere-proximal regions of the chromosomes 19 and X in wild-type (blue) and *Top6bl^Δ17Ct/Δ17Ct^* (green) mice. PRDM9-independent and -dependent hotspots are identified by orange and blue triangles, respectively. All chromosome ends are shown in Extended data Figs. 10 and 11. **e.** DMC1-SSDS signal intensity decreases in *Top6bl^Δ17Ct/Δ17Ct^* samples relative to wild-type samples in the q-arm sub-telomeric region (right panel). The DMC1-SSDS signal ratio within hotspots (log2-fold change estimated by DESeq2) was computed over 5Mb-windows with a 1Mb-step. Values were averaged for autosomes and plotted separately for the X chromosome. The same analysis was performed in the 50Mb adjacent to p-arm telomeres (left panel). As the genomic DNA sequence of p-arms has not been assembled, it is immediately flanked by centromeric and q-arm DNA sequences, where the signal can be mapped and quantified. **f.** The q-arm sub-telomeric region effect in *Top6bl^Δ17Ct/Δ17Ct^* mice. The averaged DMC1-SSDS signal ratio (*Top6bl^Δ17Ct/Δ17Ct^*/wild-type) of the last ten hotspots of a given chromosome was compared to the averaged DMC1-SSDS signal ratio of ten randomly chosen, non-telomeric consecutive hotspots in the same chromosome. Boxplots represents the log2 fold-change (FC) between these values (sub-telomeric/non-sub-telomeric) for ten randomizations. The control is shown in Extended data Fig. 8c. **g.** Decreased (green), increased (red), or unchanged (grey) hotspot density within the 3Mb sub-telomeric (Subtelo) region relative to the non-sub-telomeric regions (NonSubtelo). Decreased, increased and unchanged hotspots were determined from the DESeq2 analysis, as shown in panel c. The densities of decreased and unchanged hotspots in the two studied regions were compared using the Pearson’s Chi-square test, and p-values were adjusted for multiple testing using the Benjamini & Yekutieli method. Yellow stars indicate p-value <0.05. The Chi-square test results are provided in Extended data Table 2.

We tested this possibility by monitoring synapsis and bivalent formation on autosomes and on the X and Y chromosomes. Synapsis formation was normal on autosomes at pachynema in both mutants (Extended data Table 4). However, the X and Y chromosomes were frequently unsynapsed in pachytene nuclei (41% of *Top6bl^W562A/W562A^*, 72% of *Top6bl^Δ17Ct/Δ17Ct^* vs 18% in wild-type) (Fig. 7a). At metaphase, an increased frequency of nuclei with 21 DAPI staining bodies (Fig. 7b and 7d) and a decreased number of XY bivalent (Fig. 7c and 7e) was detected in both *Top6bl* mutants. We conclude that the decreased DMC1 signal on the X chromosome sub-telomeric region is compatible with decreased DSB activity rather than with rapid DMC1 turnover. We propose that the TOPOVIBL-REC114 interaction is specifically required at sub-telomeric regions for full DSB activity and therefore, is essential for X/Y chromosome synapsis and segregation.

**Fig. 7.**
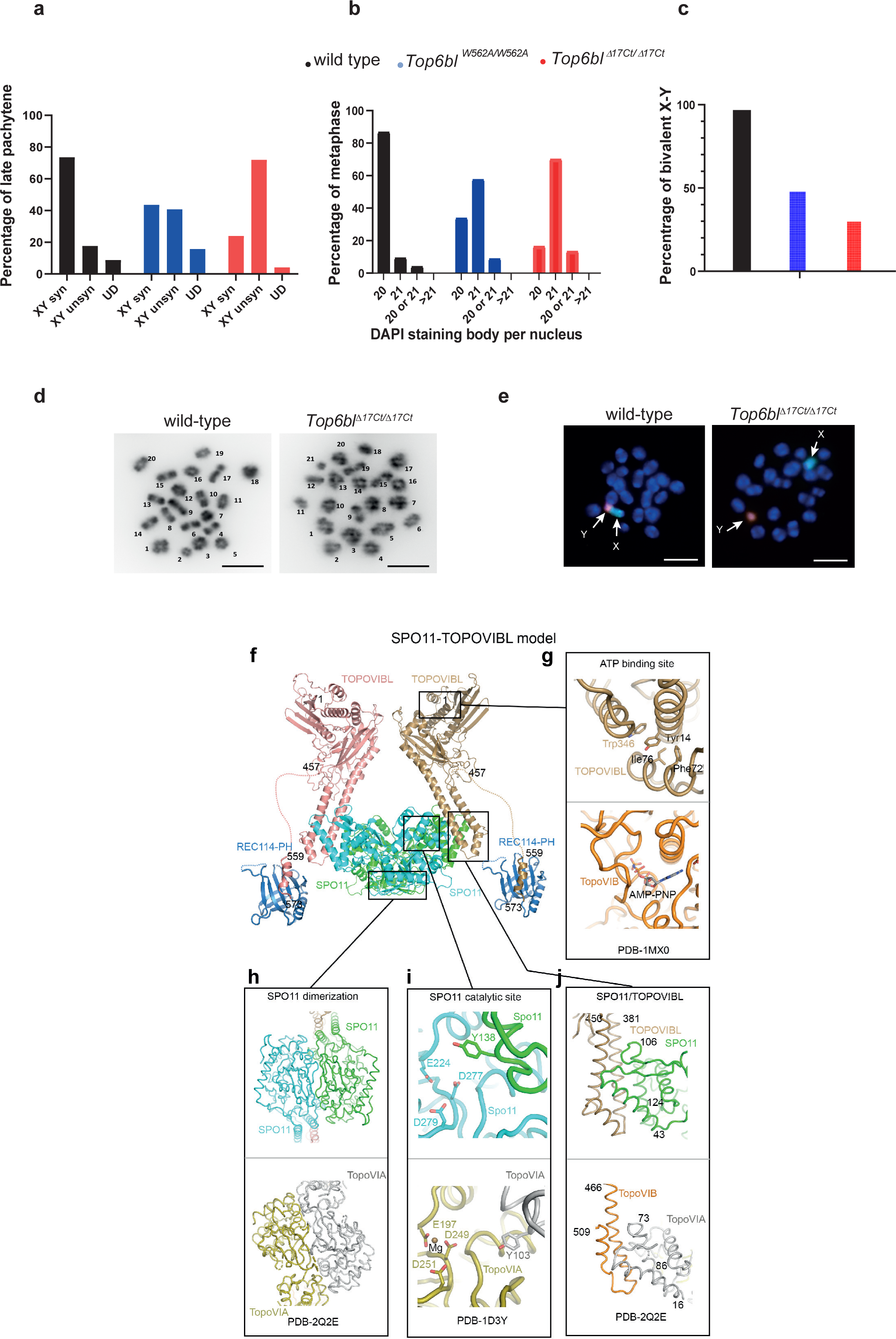
Defective XY chromosome synapsis in *Top6bl* mutants; model of the TOPOVIBL/SPO11/REC114 complex. **a.** Synapsis quantification between the X and Y chromosomes at pachynema. Synapsis formation was monitored on spreads from late pachytene spermatocytes from adult wild-type (+/+, *Top6bl^+/W562A^* or *Top6bl^+/Δ17Ct^*), *Top6bl^W562A/W562A^*, and *Top6bl ^Δ17Ct/Δ17Ct^* mice, identified by staining with γH2AX, SYCP3 and SYCP1. Number of nuclei: wild-type (227), *Top6bl^W562A/W562A^* (108), *Top6bl ^Δ17Ct/Δ17Ct^* (171). syn: synapsed; unsyn: unsynapsed; UD: undefined. The synapsed/unsynapsed ratios were significantly different between wild-type and *Top6bl^W562A/W562A^*and *Top6bl ^Δ17Ct/Δ17Ct^* spermatocytes (Pearson’s Chi-Square, 26.31 and 115.15, respectively). **b.** Quantification of bivalent formation at metaphase I. Percentage of metaphases with 20, 21, 20 or 21, or >21 DAPI-stained bodies per nucleus from adult wild-type (*Top6bl ^+/Δ17Ct^*), *Top6bl^W562A/W562A^*, and *Top6bl^Δ17Ct /Δ17Ct^* mice (n=2 mice per genotype). Number of nuclei: 75, 80, and 97, respectively. The number of nuclei with 20 and 21 bivalents was significantly different between wild-type and *Top6bl^W562A/W562A^* and between wild-type and *Top6bl ^Δ17Ct/Δ17Ct^* metaphases (Pearson’s Chi-Square: 44.4 and 78.8 respectively). **c.** Quantification of X and Y bivalents at metaphase I. The X and Y chromosomes were detected by FISH in metaphase spreads from adult wild-type (*Top6bl ^+/Δ17Ct^*), *Top6bl^W562A/W562A^*, and *Top6bl ^Δ17Ct/Δ17Ct^* mice (n=2 mice per genotype). Number of nuclei: 84, 144, and 164 for wild-type, *Top6bl^W562A/W562A^* and *Top6bl^Δ17Ct /Δ17Ct^*, respectively. The percentages of metaphase spreads with XY bivalents were significantly different between wild-type and *Top6bl^W562A/W562A^,* and between wild-type and *Top6bl ^Δ17Ct/Δ17Ct^* (Chi-Square Pearson: 54.2 and 96.9, respectively). **d.** Representative images of DAPI-stained wild-type and *Top6bl ^Δ17Ct/Δ17Ct^* metaphase spreads. Scale bar, 10μm. DAPI-stained bodies are numbered (arbitrarily): 20 are observed in wild-type, and 21 in *Top6bl ^Δ17Ct/Δ17Ct^* metaphase spreads. **e.** Representative images of a wild-type spermatocyte nucleus with the X and Y chromosomes forming a bivalent (left) and of a *Top6bl ^Δ17Ct/Δ17Ct^* spermatocyte nucleus with separated X and Y chromosomes (right). Blue, nuclei (DAPI staining); green, X chromosome probe; red, Y chromosome probe. Scale bar, 10μm. **f.** The structure of the complex formed by SPO11 and the C-terminal part of the TOPOVIBL transducer domain was modelled with AlphaFold ^27^. The full-length TOPOVIBL modelled structure (AF-J3QMY9-F1) was superimposed on the transducer domain. The structure of REC114 PH domain bound to TOPOVIBL C-terminus was determined in this study and is shown linked to the SPO11-TOPOVIBL complex via a long-disordered linker. **g.** According to the AlphaFold model, the putative ATP binding site of TOPOVIBL is degenerated. Secondary structure elements forming the ATP binding site in *S. shibatae* (PDB – 1MXO, lower panel), are organized differently in mouse TOPOVIBL. Consequently, a corresponding ATP binding site is not formed. Mouse TOPOVIBL also lacks the N-terminal “strap” region and the H2TH domain, both involved in TopoVIB dimerization, indicating that TOPOVIBL dimerization, if occurs, should differ from that of TopoVIB ^37^. **h.** The modelled SPO11 dimerization interface resembles the one described for archaeal TopoVIA (bottom), including the formation of a pseudo-continuous β-sheet by the two protomers ^38^. **i.** The modelled SPO11 catalytic site is equivalent to the one previously described for TopoVIA (PDB – 1D3Y, lower panel) where the two protomers contribute three negatively charged residues that co-ordinate the magnesium atom and the catalytic tyrosine ^38^. **j.** The modelled SPO11-TOPOVIBL interface is similar to that of the archaeal TopoVI complex (PDB - 2Q2E, lower panel), with the additional helix of the TOPOVIBL transducer domain ^9, 10^.

These molecular and cytological phenotypes should have specific consequences on meiotic prophase and downstream events, because unsynapsed X/Y chromosomes induce metaphase I arrest ^35, 36^. Indeed, *Top6bl^W562A/W562A^* and *Top6bl^Δ17Ct/Δ17Ct^* spermatocytes proceed through prophase like wild-type spermatocytes, but they arrested in metaphase and many cells were apoptotic, particularly in the *Top6bl^Δ17Ct^* mutant. This metaphase I arrest was correlated with reduced sperm production in both mutants and fertility loss in *Top6bl^Δ17Ct/Δ17Ct^* mice (Extended data Figs. 5e-5k).

## Discussion

A central question after the identification of the axis-associated proteins essential for meiotic DSB formation is to understand their function. Here, we found that TOPOVIBL CTD directly interacts with REC114 PH domain, and identified residues required for this interaction *in vitro*. We propose that TOPOVIBL can bind simultaneously to SPO11 through its transducer domain and to REC114 through the CTD. Both mouse TOPOVIBL and SPO11 structures have been modeled by the Alphafold structure prediction program ^27^ with high predicted accuracy (AlphaFold Protein Structure Database code-AF-J3QMY9 and AF-Q9WTK8). We further used AlphaFold ^27^to model the SPO11/TOPOVIBL complex and included REC114 (Fig. 7f). Compared to the TopoVIB structure ^37^, the TOPOVIBL ATP binding site seems degenerated ^5^ and likely does not allow ATP binding (Fig. 7g). TOPOVIBL also lacks both the N-terminal “strap” region involved in dimerization and the H2TH domain. Thus, the ATP mediated dimerization, observed for archaeal TopoVIB, is unlikely to occur in the case of TOPOVIBL. SPO11 however can be modeled as homodimer and its catalytic site, formed by the two protomers is equivalent to the one described for TopoVIA ^38^ (Figs. 7h and 7i). Finally, the modeled SPO11-TOPOVIBL interface resembles the archaeal TopoVI complex including an additional helix of the transducer domain (Fig. 7j). The REC114-TOPOVIBL complex, as characterized in this study, is connected by the flexible TOPOVIBL CTD to the SPO11/TOPOVIBL core (Fig. 7f). Although the exact REC114 effect on SPO11/TOPOVIBL organization/conformation is unknown, REC114 binding to TOPOVIBL could stabilize the SPO11/TOPOVIBL complex dimerization. This hypothesis is supported by the stoichiometry (2:1) of the Rec114/Mei4 complex in *S. cerevisiae* ^15^ and by the observation that REC114 could bind simultaneously to TOPOVIBL and MEI4 (Extended data Fig. 1g). Nevertheless, REC114 interactions are certainly more complex because its PH domain also interacts with ANKRD31 ^25^, thus likely competing with TOPOVIBL (Fig. 2j and Extended data Fig. 1j). Moreover, REC114 interacts with IHO1 in yeast-two hybrid assays ^24^. The interplay of these different interactions remains to be determined, but disrupting the interaction between TOPOVIBL and REC114 may have additional consequences on other interactions.

The observation of meiotic DSB defects in female and male mice harbouring *Top6bl* mutations indicates that *in vivo* REC114 acts by directly interacting with the SPO11/TOPOVIBL complex. The stronger phenotype of mice carrying the *Top6bl^Δ17Ct^* mutation (compared with *Top6bl^W562A^*) is explained by the absence of interaction in *Top6bl^Δ17Ct^*, whereas the interaction is only weakened in *Top6bl^W562A^*. It also indicates that the C-terminal helix is the major interaction site between TOPOVIBL and REC114. Moreover, the distinct phenotypes show that REC114 is a regulatory subunit of the activity, and not just an accessory factor of the TOPOVIL complex. The reduced DSB activity in oocytes fits exactly the simple interpretation that REC114 binding to TOPOVIBL is required for the catalytic activity. The male phenotype highlights REC114 double role in fine tuning DSB activity by regulating the timing of DSB formation genome-wide and DSB formation at sub-telomeric regions. It is remarkable that the delayed DSB formation with respect to axis formation in *Top6bl^Δ17Ct/Δ17Ct^* mice does not alter homologous synapsis, and seems to allow weak DSB sites to be more active than in wild-type. This could be explained if DSB sites have a window of opportunity to be active but can be turned off upon synapsis, as shown in *S. cerevisiae* ^39^. Delaying synapsis would thus allow weak DSB sites to be active. The sub-telomeric effect in *Top6bl^Δ17Ct/Δ17Ct^* mice potentially applies to all chromosomes and is not restricted to the four chromosomes (chr. 4, 9, 13 and X) shown to accumulate REC114, MEI4, IHO1 and ANKRD31 aggregates ^25, 26^. The property we detected is therefore distinct from those aggregates, which is also consistent with the observation that *Top6bl* mutants do not have any detectable defect in the formation of those aggregates (Extended data Fig. 7a). We do notice inter-chromosomal quantitative differences of the extent of decreased DSB activity in *Top6bl^Δ17Ct/Δ17Ct^* mice that may at least in part be due to inter-chromosomal differences in hotspot activity within sub-telomeric regions (Extended data Fig. 9e). Sub-telomeric regions display specific features (i.e. nuclear organization during meiosis ^40, 41^, architecture, organization and other epigenetic properties) that may influence the control of SPO11/TOPOVIBL activity. These regions are known to behave differently from chromosome arms for meiotic DSBs, based on their sensitivity to the expression of a GAL4BD-SPO11 fusion protein on DSB activity ^42^, to a lower DMC1/SPO11-oligonucleotides ratio ^43^ and to differential activities in male and female meiosis ^44^. It is also possible that mice expressing SPO11β-only (the long isoform of SPO11) and shown to have decreased DSB activity in the PAR, have a specific decrease of DSB activity near autosome ends ^36^. This would imply a specific regulation of SPO11 near chromosome ends, as the phenotypes due to the *Top6bl^Δ17Ct^* mutation suggest. As SPO11α (the short isoform of SPO11) is lacking the interaction domain with TOPOVIBL, it is not expected to be catalytically active. SPO11α may rather repress an inhibitor of TOPOVIL. One possibility is that this inhibitor interferes with the REC114-TOPOVIBL interaction. Such scenario, although certainly others can be envisioned, would fit with the potential similarity of phenotypes between *Spo11β*-only and *Top6bl^Δ17Ct/Δ17Ct^* male mice, including the XY synapsis defect. Overall, these observations imply that i) the TOPOVIBL-REC114 interaction is not essential for DSB activity in all genomic contexts. In *A. thaliana,* Rec114 role is dispensable ^45^ and it would be interesting to test if the absence of Rec114 leads to a DSB delay; ii) REC114 activity senses or responds to specific chromosomal features. Several studies have highlighted differences of recombination and/or chromosome organization between sexes as well along chromosomes but which links to DSB activity remain to be determined ^44, 46^.

Programming hundreds of DSBs in the genome is a challenge for the cell, and the current knowledge that the catalytic complex of SPO11/TOPOVIBL requires several other proteins is coherent with the need to regulate these events. Here, we described the central role of REC114 in SPO11/TOPOVIBL activity through its direct interaction with TOPOVIBL, thus highlighting a first level of this regulation. Moreover, the DSB program must be executed in two very different cell types (oocytes and spermatocytes), and our findings show that the REC114-TOPOVIBL interaction is sensitive to these differences. Other components of the meiotic DSB machinery and their potential multiple interactions should contribute to SPO11/TOPOVIBL regulation. Additional directed-mutagenesis studies will unravel them and will identify the complex(es) active *in vivo*.

## Supporting information

Nore_Juarez_supmarerial

## Acknowledgments

We thank the following Biocampus facilities from Montpellier for their service: the Réseau des Animaleries de Montpellier (RAM) for animal care, the Réseau d’Histologie Expérimentale de Montpellier (RHEM) for histology, and Montpellier Resources Imagerie (MRI) for microscopy. We thank all lab members for discussion and Pauline Auffret for support in bioinformatic analysis. We thank Scott Keeney for the anti-REC114 antibody, Attila Toth for the anti-ANKRD31 and anti-IHO1 antibodies. We thank Caroline Mas for assistance with ITC and the staff of the ESRF-EMBL (European Synchrotron Radiation Facility-European Molecular Biology Laboratory) Joint Structural Biology Group, particularly Andrew McCarthy, for access to and help with the ESRF beamlines. We thank the EMBL high-throughput crystallization facility (HTX).

## Methods

### Resource availability

Further information and requests for resources and reagents should be directed to and will be fulfilled by the lead contact Bernard de Massy.

### Mouse strains

Mice were in the C57BL/6J background. Mice carrying the homozygous mutant alleles Top6bl<em1(W562A)BdM> and Top6bl<em2(delta17)BdM> were named *Top6bl^W562A/W562A^* and *Top6bl^Δ17Ct/Δ17Ct^*, respectively. *Top6bl^-/-^* mice carry the Gm960^em2Arte^ allele, a null allele due to a 5bp deletion in *Top6bl* ^5^. All experiments were carried out according to the CNRS guidelines and were approved by the ethics committee on live animals (project CE-LR-0812 and 1295).

### Generation of mutant mice by CRISPR/Cas9

Mutant mice were created at the Jackson Laboratory using the CRISPR-Cas9 technology with three different guides and two different donor oligos (Extended data Table 3). Guides were selected to minimize off-target effects. The donor oligos were designed to change the W562 codon TGG (W) to GCG (A), and to introduce a silent mutation (A to G) to generate a *Pst*I restriction site. The Top6bl<em1(W562A)BdM> allele, named *Top6bl^W562A^*, is the result of homologous recombination. The Top6bl<em2(delta17)BdM> allele, named *Top6bl^Δ17Ct^*, is the result of non-homologous repair and has a 4bp deletion (Extended data Fig. 2). Founders were backcrossed with C57BL/6J animals to obtain heterozygous animals. The predicted TOPOVIBL protein expressed from each mutant allele and the genotyping strategies are shown in Extended data Fig. 2.

### Yeast two-hybrid assays: clones, assays, western blotting

All plasmids used in yeast two-hybrid assays were cloned with the Gateway® Gene Cloning Technology (Invitrogen) and transformed in the AH109 and Y187 haploid yeast strains (Clontech). AH109 and Y187 cells were transformed with Gal4 DNA binding domain (GBD) fusion plasmids derived from pAS2 and Gal4 activation domain (GAD) fusion plasmids that were obtained from pGAD. Purified colonies of diploid strains were streaked on synthetic medium (SD) plates lacking leucine and tryptophan (-LW), or leucine, tryptophan and histidine (-LWH), or leucine, tryptophan and histidine with 5 mM amino-triazole (-LWH+3AT), or leucine, tryptophan, histidine and adenine (-LWHA). Dilution assays were performed by spotting cells on -LW, -LWH, -LWH+3AT and -LWHA plates that were incubated at 30°C for 3 days. For verification of protein expression, protein extracts were prepared and analysed by western blotting, as previously described ^5^, with anti-GAD (1:3000; UPSTATE-06-283) and anti-GBD (1:1000; SIGMA; G3042) antibodies.

### Protein expression, purification and crystallization

His-tagged mouse REC114 15-159 was expressed in *E. coli* BL21-Gold (DE3) cells (Agilent) from the pProEXHTb expression vector (Invitrogen). The protein was first purified by affinity chromatography using the Ni^2+^ resin. After His-tag cleavage with the TEV protease, it was purified through a second Ni^2+^ column and size-exclusion chromatography. The pure protein was concentrated to 20 mg ml^-1^ in a buffer containing 20 mM Tris, pH 8.0, 200 mM NaCl and 5 mM β-mercaptoethanol, and supplemented with a three-fold molar excess of TOPOVIBL peptide (559-EDLWLQEVSNLSEWLNPG-576). The complex was crystallized using the hanging drop vapor diffusion method at 20°C. The best diffracting crystals grew within seven days in a solution containing 1.6M MgSO4, 100mM MES (pH 6.5), and 10% (v/v) dioxane. For data collection at 100 K, crystals were snap-frozen in liquid nitrogen with a solution containing mother liquor and 25% (vol/vol) glycerol.

### Data collection and structure determination

Crystals of the mouse REC114-TOPOVIBL complex belong to the space group *P*6122 with the unit cell dimensions *a*, *b* = 108.7 Å and *c* = 83. Å. The asymmetric unit contains one REC114-TOPOVIBL dimer and has a solvent content of 68%. A complete native dataset was collected to a 2.5 Å resolution, partially extending to 2.26Å on the ESRF beamline ID30B. The data were processed using autoPROC ^47^. Phases were obtained by molecular replacement using PHASER ^48^ with the crystal structure of the REC114 PH domain (PDB code: 6HFG) as search model. The initial map was improved using the prime-and-switch density modification option of RESOLVE ^49^. After manual model rebuilding with COOT ^50^, the structure was refined using Refmac5 ^51^ to a final *R*-factor of 22.6% and *R*free of 24.9% (Extended data Table 1) with all residues in the allowed (95.2% in favored) regions of the Ramachandran plot, as analysed by MOLPROBITY ^52^. A representative part of the 2*F*o − *F*c electron density map covering the TOPOVIBL-REC114 interface is shown in Extended data Fig. 1b.

### Pull-down assays

Full-length REC114 and its variants were cloned as Strep-tag fusions into pRSFDeut-1 (Novagen). TOPOVIBL 1-385 was cloned as a 6xHis-SUMO fusion protein in pETM11, and TOPOVIBL 450-579 and its mutated versions as 6xHis fusion proteins in pProEXHTb. Proteins were expressed individually in *E. coli* BL21Gold (DE3) cells. For analysis of the structure-based mutants, half of each culture was used for Strep-tag pull-down assays and the other half for His-tag pull-down assays. For Strep-tag pull-down assays, TOPOVIBL proteins were affinity-purified using Ni^2+^ resin. Following cell disruption, the Strep-REC114-containing supernatant was added to a Strep-Tactin XT resin that was then extensively washed. The REC114-bound resin was then divided for individual pull-down experiments. Equal amounts of TOPOVIBL proteins were added and incubated with the REC114-bound resin with agitation at 5°C for 10 min. Columns were then washed with a buffer containing 100mM Tris pH 8, 150mM NaCl, 5mM β-mercaptoethanol. Bound proteins were eluted by addition of 50mM of D-Biotin, and analysed on 15% SDS-PAGE. The His-tag pull-down experiments were performed in a similar way, but REC114 proteins were first purified on Strep-Tactin XT resin and then added in equal amounts to the TOPOVIBL-bound Ni^2+^ resin.

#### Competition assay

TOPOVIBL 450-579 was purified using Ni^2+^ resin as described above. ANKRD31 1808-1857 was cloned as 6xHis-MBP fusion in pETM41. Strep-REC114 1-159 and ANKRD31 1808-1857 were individually expressed in *E. coli* BL21Gold (DE3) cells. Following cell disruption, supernatants containing soluble Strep-REC114 1-159 and ANKRD31 1808-1857 were mixed and loaded onto Strep-Tactin XT resin. The resin bound with the REC114 1-159-ANKRD31 1808-1857 complex was then extensively washed and divided into 1ml fractions for individual competition assays by adding an increasing amount of purified His-tagged TOPOVIBL 450-579 (1mg, 5mg, 10mg and 20mg). After 10 min incubation at 5°C, columns were washed with a buffer containing 100mM Tris pH 8, 150mM NaCl, 5mM β-mercaptoethanol, and bound proteins were eluted by adding 50mM of D-Biotin, and analysed on 12% Tris-Tricine SDS PAGE.

### Isothermal Titration Calorimetry (ITC)

ITC experiments were performed at 25°C using an ITC200 microcalorimeter (MicroCal). Experiments included one 0.5µl injection and 18-20 injections of 1.5-2µL of 0.3-1.8 mM TOPOVIBL (TOPOVIBL 450-579, TOPOVIBL 559-576 or TOPOVIBL 559-576 W562A) into the sample cell that contained 30-40 µM REC114 15-159 in 20 mM Tris (pH 8.0), 200 mM NaCl, 5% glycerol and 5mM β-mercaptoethanol. The initial data point was deleted from the data sets. Binding isotherms were fitted with a one-site binding model by nonlinear regression using the Origin software, version 7.0 (MicroCal).

### Preparation of mouse protein extracts, immunoprecipitation and western blotting

Whole cell protein extracts were prepared from eight frozen testes collected at 14 dpp for each genotype. After protein extraction by homogenizing cells with a Dounce homogenizer in HNTG buffer (150 mM NaCl, 20 mM HEPES pH7.5, 1% Triton X100, 10% glycerol, 1 mM MgCl, Complete protease Inhibitor (Roche 11873580001)), followed by sonication, benzonase (250 U) was added at 4°C for 1h. After centrifugation (16000 g, 4°C, 10 min) to remove debris, immunoprecipitation was performed with 5 µg of homemade anti-TOPOVIBL antibody. For each immunoprecipitation, 3.5 mg of whole cell protein extract and 50 µl of Protein A Dynabeads (Invitrogen 10001D) were used. Then, immunoprecipitates were resuspended in 40 µl of Laemmli buffer and TOPOVIBL immunoprecipitation was assessed by western blotting with a homemade affinity-purified anti-TOPOVIBL (1/1000) antibody followed by an anti-rabbit LC mouse monoclonal secondary antibody (Jackson ImmunoResearch 211-032-171).

### RT-PCR assays

Total RNA was extracted with the miRNeasy Mini Kit (Qiagen) according to the manufacturer’s instructions. For RT-PCR, first-strand DNA was synthesized using oligo d(T)18 (Ambion), SuperScriptIII (Invitrogen), and total RNA (1-2 µg) from 16dpc ovaries.

The open reading frames of *Top6bl* and *Spo11* were amplified using standard PCR conditions and the primer pairs Oli63/Oli70 and Spo11:116U22/Spo11:655L22, respectively (Extended data Table 3). PCR cycling conditions were: 3 min at 94°C, 35 cycles of 30 sec at 94°C, 30 sec at 54°C, and 2 min or 30 sec at 72°C, followed by 5 min at 72°C. *Top6bl* ORF was then digested with the *Eci*I enzyme.

### Histological analysis of paraffin sections and TUNEL assay

Mouse testes or ovaries were fixed in Bouin’s solution for Periodic Acid Schiff (PAS) staining of testes and haematoxylin eosin staining of ovaries. Fixation was in 4% paraformaldehyde/1X PBS for immunostaining and TUNEL assay. Testes and ovaries were embedded in paraffin and cut in 3µm-thick sections. Sections were scanned using the automated tissue slide-scanning tool of a Hamamatsu NanoZoomer Digital Pathology system. TUNEL assay was performed with the DeadEnd Fluorometric TUNEL System (Promega), according to the manufacturer’s protocol.

### Spermatozoid counting

After dissection of the epididymis caudal part from adult testes (2 month-old), spermatozoids were extracted from the epididymis by smashing or crushing the tissue in PBS. After homogenization by pipetting, 10µl of the soluble part was diluted in 1mL of water, and spermatozoids were counted.

### Immunocytology

Spread from spermatocytes and oocytes were prepared with the dry down technique, as described ^53^, and immunostaining was performed as described ^54^.

### Antibodies

Guinea pig anti-SYCP3 ^54^, rabbit anti-SYCP1 (Abcam, 15090), rabbit anti-DMC1 (Santa Cruz, H100), anti-RPA2 (Abcam), anti-MEI4 ^21^, anti-REC114 ^23^, anti-REC114 (gift from S. Keeney), anti-ANKRD31 ^26^, anti-IHO1 ^24^, and mouse monoclonal anti-phospho-histone H2AX (Ser139) (γH2AX) (Millipore, 05-636) antibodies were used for immunostaining. Homemade affinity purified anti-TOPOVIBL antibody: rabbits were injected with full-length mouse His-TOPOVIBL protein prepared from *E. coli* inclusion bodies. Rabbit serum was purified by affinity using His-TOPOVIBL purified from inclusion bodies.

### Metaphase spread preparation

Tubules from decapsulated testes were pulled apart in 1% trisodium citrate and transferred into a 15 ml tube. After pipette homogenization and 1min sedimentation, the cell-containing supernatant was transferred in a new 15 ml tube. Following the same procedure, the tubule pieces were rinsed twice with 3ml of 1% trisodium citrate. The cell solution was centrifuged at 180g for 10min, and the pellet resuspended in 100µl of supernatant. Then, 3 ml of methanol: acetic acid: chloroform (3:1:0.05) solution was added drop by drop to the cell solution (rolling the first drops down the sides of the tube while flicking the tube). Cells were then centrifuged at 180g for 10 min and resuspended in 100µl of supernatant, and 3 ml of methanol: acetic acid (3:1) was added to the tube. After 10min of incubation at room temperature, cells were centrifuged again (180g for 10 min) and resuspend in ∼1 ml methanol: acetic acid (3:1). To prepare the slides, 40µl of the cell suspension was dropped from a height of ∼40cm onto a slide that was held titled at 45°. Slides were dried in a humid chamber.

### FISH for chromosome painting

X (D-1420-050-FI; D-1420-050-OR) and Y (D-1421-050-FI; D-1421-050-OR) chromosome-specific probes were used according to the manufacturer (Metasystems Probes).

### Image analysis

For focus quantification, all images were deconvoluted using the Huygens software. Image J was used to quantify foci that colocalized with the chromosome axis defined by SYCP3 staining.

For γH2AX quantification in oocytes, signal intensity was obtained using Cell Profiler on non-deconvoluted images. Integrated intensity was used for the analysis. For spermatocytes, both Cell Profiler and Image J quantifications were performed and gave similar results. The output of Image J integrated intensity is presented.

Staging criteria were as follows. Pre-leptotene nuclei had weak SYCP3 nuclear signal and no or very weak γH2AX signal; early leptotene nuclei were γH2AX-positive and with only short SYCP3 fragments; mid leptotene nuclei were γH2AX-positive and with short and long SYCP3 fragments; late leptotene were γH2AX-positive and with only long SYCP3 fragments; zygotene nuclei had partially synapsed homologs; and pachytene cells had all 19 autosomes fully synapsed (Extended data Fig. 6a).

### DMC1-SSDS analysis

#### Library preparation and sequencing

DMC1 ChIP-seq was performed as described in ^55^ using a goat anti-DMC1 antibody (Santa Cruz, C-20). Six *Top6bl^Δ17Ct/Δ17Ct^* testes and two wild-type testes from 12 to 25-week-old mice were used for each replicate. Sequencing was performed on a HiSeq 2500 instrument in paired-end mode (2x150bp).

#### DMC1-SSDS mapping and hotspot identification

After quality control and read trimming to remove adapter sequences and low-quality reads, DMC1 ChIP-SSDS reads were mapped to the UCSC mouse genome assembly build GRCm38/mm10. The previously published method ^33^ was used for DMC1-SSDS read mapping (i.e. the BWA modified algorithm and a customized script that were specifically developed to align and recover ssDNA fragments). A filtering step was performed on the aligned reads to keep only non-duplicated and high-quality uniquely mapped reads with no more than one mismatch per read. To identify meiotic hotspots from biologically replicated samples in DMC1-SSDS, the Irreproducible Discovery Rate (IDR) method was used, as done in our previous studies. This method was developed for ChIP-seq analysis and extensively used by the ENCODE and modENCODE projects ^56^. The framework developed by Qunhua Li and Peter Bickel’s group (https://sites.google.com/site/anshulkundaje/projects/idr) was followed. Briefly, this method allows testing the reproducibility within and between replicates by using IDR statistics. Following their pipeline, peak calling was performed using MACS version 2.0.10 with relaxed conditions (--pvalue=0.1 --bw1000 -- nomodel --shift400) for each of the two replicates, the pooled dataset, and pseudo-replicates that were artificially generated by randomly sampling half of the reads twice for each replicate and the pooled dataset. Then IDR analyses were performed and reproducibility was checked. Final peak sets were built by selecting the top N peaks from pooled datasets (ranked by increasing p values), with N defined as the highest value between N1 (the number of overlapping peaks with an IDR below 0.01, when comparing pseudo-replicates from pooled datasets) and N2 (the number of overlapping peaks with an IDR below 0.05 when comparing the true replicates, as recommended for the mouse genome). Hotspot centring and strength calculation were computed following the method described by Khil et al. ^33^. All read distributions and signal intensities presented in this work were calculated after pooling reads from both replicates, if not otherwise stated. When DSB maps were compared between mouse genotypes, the 1bp-overlaps were restricted to the central 400bp of hotspots (+/- 200bp around the peak centre). For correlation plots, the type 1 single-strand DNA signal was library-normalized (fragment per million).

#### Differential analysis of hotspot strength

To compare hotspot usage between *Top6bl^Δ17Ct/Δ17Ct^* and wild-type mice, DMC1-SSDS signal intensity was compared at the 13562 wild-type hotspots using DESeq2 ^57^. Among these hotspots, 4207 hotspots (31%) were filtered out with the default independent filtering option using the mean of normalized counts as filter statistic. The aim was to remove sites with too low counts (mean count below 22) that have zero or low chance of showing significant differences to increase the detection power for the other sites. For the 9355 tested hotspots, log fold change shrinkage was performed to correct data dispersion using the *apeglm* method^58^. The hotspots with increased or decreased DMC1-SSDS signal intensity were then determined using an adjusted p-value threshold of 0.1 and a log2 fold change value below or above zero, respectively. This led to the identification of 214 increased (1.2% of total hotspots, and 2% of the tested ones), 1098 decreased (8% of total hotspots, and 12% of the tested ones), and 8043 unchanged hotspots in *Top6bl^Δ17Ct/Δ17Ct^* mice (Fig. 6c).

#### Analysis of hotspot distribution at sub-telomeric regions

Visual inspection of DMC1-SSDS signal intensity along chromosomes and the localization of hotspots with decreased DMC1-SSDS signal intensity in *Top6bl^Δ17Ct/Δ17Ct^* suggested that much of the DMC1 signal decrease was located near the q-arm telomeres (telomeres distant from centromeres). To test whether this biased distribution was significant, each hotspot was annotated as sub-telomeric when within the sub-telomeric region defined with a variable size from 1 to 10Mb. For each sub-telomeric region, the numbers of unchanged, decreased and increased hotspots in the sub-telomeric versus the non-sub-telomeric region (i.e. the rest of the chromosome) were counted. Pearson’s Chi-square tests were computed (by taking into account or not the increased hotspots) and p-values were adjusted for multiple testing using the Benjamini & Yekutieli method. Megabase-normalized counts (hotspot density) were measured and plotted (Fig. 6g for sub-telomeric regions of 3Mb). For each chromosome, the sub-telomeric over non-sub-telomeric ratio of hotspot density was calculated for each unchanged, decreased or increased hotspot category (Extended data Fig. 9d). Alternatively, to evaluate hotspot activity without a fixed distance from the telomere, for each chromosome, the *Top6bl^Δ17Ct/Δ17Ct^*/wild-type signal ratio was averaged over the ten most q-arm telomeric hotspots, and then compared to the averaged signal ratio measured over another set of ten consecutive hotspots randomly chosen along the chromosome (excluding the last tens). Then, the ratio of these two mean values was computed. The procedure was repeated 10 times, each time with a different random set of 10 non-sub-telomeric hotspots. The sub-telomeric effects are presented in Fig. 6f as the distribution of these ratios. As control, these ratios were computed not between the last ten and ten non-sub-telomeric hotspots, but between two non-sub-telomeric random hotspot sets (procedure repeated 10 times) (Extended data Fig. 9c).

#### Statistical analysis

The statistical analyses of cytological observations were done with GraphPad Prism 9. The nonparametric Mann-Whitney test was used to compare number of foci, and the Pearson’s Chi square test to compare distributions, as indicated in the figure legends. The Chi square tests were performed at http://vassarstats.net/ and https://www.quantitativeskills.com/sisa/index.htm. Box plots (25-75 percentiles) show the median and 5-95 percentiles. Statistical tests for DMC1-SSDS data were done using R version 4.0.3. All tests and p-values (n.s., not significant. *P < 0.05, **P < 0.01, ***P < 0.001, ****P < 0.0001) are provided in the corresponding legends and/or figures.

#### Data deposition

The DMC1-SSDS raw and processed data for this study have been deposited under a confidential status in the European Nucleotide Archive (ENA) at EMBL-EBI (accession number PRJEB43730) (https://www.ebi.ac.uk/ena/browser/view/PRJEB43730) and will be released as soon as the present paper is accepted for publication. Data are available upon request during the reviewing process. Structural data has been submitted to PDB (accession number 7QWV).

#### Extended data

**Extended data Fig. 1a.** Superdex 200 gel filtration elution profiles of REC114 15-159 (blue), TOPOVIBL 450-579 (red), and their complex (green).

**b.** Representative part of the 2*F*o-*F*c electron density map covering the TOPOVIBL-REC114 interface contoured at 1.0 σ.

**c.** The REC114-TOPOVIBL complex models predicted by AlphaFold ^27^. The structure of the REC114 PH domain is shown as surface in blue. Key interacting residues of TOPOVIBL are shown as sticks. Most of the interactions observed in the mouse crystal structure are conserved in the predicted complexes of other species. In *D. rerio*, the residues corresponding to mouse Val566 is Leu555. This mutation seems to be compensated by the valine to glutamine mutation at position 92 in REC114.

**d.** His-tag pull-down assays using the proteins shown in Fig. 2g. Strep-tag purified REC114 (WT or mutant) was loaded on Ni^2+^ columns bound with WT or mutated TOPOVIBL proteins.

**e.** ITC measurement of the interaction affinity between REC114 15-159 and the TOPOVIBL 559-576 peptide harbouring the W562A mutation (the wild-type control is in Fig. 1g).

**f.** Comparison of the S200 gel filtration elution profiles of wild-type TOPOVIBL 450-579 and the W562A mutant. The two profiles are essentially identical, indicating that the W562A mutation does not affect the overall structure of this TOPOVIBL fragment.

**g.** Superose 6 gel filtration elution profiles of full-length REC114 in complex with MEI4 1-27 (blue), TOPOVIBL 450-579 (red) and their complex (green).

**h-i**. Comparison of the REC114-TOPOVIBL (h) and the ANKRD31-REC114 (i) structures.

**j**. ITC measurement of the interaction between REC114 15-159 and TOPOVIBL 450-579 indicates the absence of binding when REC114 is pre-saturated with ANKRD31 1808-1857.

**Extended data Fig. 2a.** *Top6bl*, *Top6bl^W562A^*and *Top6bl^Δ17Ct^* cDNA sequences. The top sequence corresponds to the 65 last nucleotides of the *Top6bl* open reading frame. Orange and blue letters mark exons 15 and 16, respectively. The TGG codon in green corresponds to the W562 residue in the protein sequence. Red arrows mark the position of the predicted CRISPR-Cas9 cleavage sites, corresponding to the three guide RNAs used. TAA in black: original stop codon.

The middle sequence corresponds to the repair product by homologous recombination (HR) following CRISPR-Cas9-mediated DSB formation. The TGG codon is replaced by the GCG codon (red letters). The G residue in purple corresponds to a silent substitution to create a *Pst*I restriction site for genotyping. TAA in black: original stop codon.

The bottom sequence corresponds to the product of the deletion of four nucleotides in the sequence following DSB formation and repair (grey crossed letters). TGA in black: new stop codon due to frameshift.

**b.** Protein sequences of the C-terminal part of TOPOVIBL, TOPOVIBL-W562A and TOPOVIBL-Δ17Ct. Blue letters correspond to the original residues, bold red A to the mutant substitution of W562 in the TOPOVIBL-W562A sequence, and red letters to the nine-residue substitution in the TOPOVIBL-Δ17Ct sequence. *: Stop.

**c.** Genotyping strategy. Following DNA extraction from tails of *Top6bl^+/+^, Top6bl^+/W562A^, Top6bl^+/Δ17Ct^, Top6bl^W562A/W562A^* and *Top6bl^Δ17Ct/Δ17Ct^*mice, PCR amplification with the primer pair specific for a 569 bp region containing the mutations was performed. PCR products were digested with *Pst*I or *Eci*I. The top table presents the expected size following digestion of the different alleles (red correspond to *Pst*I digestion, and dark blue to *Eci*I digestion). Bottom panel: DNA fragments separated on agarose gels.

**d.** TOPOVIBL immunoprecipitation. Following extraction of total protein from 14 dpp mouse testes obtained from *Top6bl^+/+^, Top6bl^W562A/W562A^ Top6bl^Δ17Ct/Δ17Ct^* and *Top6bl^-/-^* mice, TOPOVIBL was immunoprecipitated using the homemade anti-TOPOVIBL antibody. Six testes of each genotype were used for one immunoprecipitation. The immunoblot was revealed using the homemade anti-TOPOVIBL antibody. The red arrow shows the TOPOVIBL expected band. *corresponds to a non-specific band (immunoglobulin heavy chain).

**e.** Comparison of the S200 gel filtration elution profile of wild-type TOPOVIBL 450-579 and TOPOVIBL Δ17Ct 450-579. Both profiles are very similar, indicating that the Δ17Ct mutation does not affect the overall structure of this TOPOVIBL fragment.

**f.** Expression of *Top6bl* assessed by RT-PCR. RNA extracted from ovaries of 16 dpc *Top6bl ^Δ17Ct/Δ17Ct^* or *Top6bl ^+/Δ17Ct^* mice was incubated with or without reverse transcriptase (RT), amplified by PCR and incubated with or without *Eci*I restriction enzyme. The size of the fragments after *Eci*I digestion are shown for the *Top6bl ^+^* and *Top6bl ^Δ17Ct^* alleles. Stars indicate non-specific amplification products. Control PCR assays for monitoring the expression of the two *Spo11* splice variants (*Spo11α* and *β*) and of actin are shown.

**Extended data Fig. 3 a.** Quantification of γH2AX signal intensity in leptotene (L)/zygotene (Z) oocytes from 16 and 18 dpc wild-type and 18dpc *Top6bl ^Δ17Ct/Δ17Ct^* mice. Number of nuclei: 64, 9, and 167, respectively. Grey bars show the mean values. P values were determined using the two-tailed unpaired Mann-Whitney test. Progression into prophase at 18dpc was: L 46%, Z 54%, Pachynema (P) 0% in *Top6bl ^Δ17Ct/Δ17Ct^*, and L 2%, Z 8%, P 90% in wild-type oocytes.

**b.** Immunostaining of RPA2, SYCP3 and γH2AX in oocytes from 16 dpc wild-type (+/+), *Top6bl^W562A/W562A^*, and *Top6bl ^Δ17Ct/Δ17Ct^* ovaries. Scale bar, 10μm.

**c.** Quantification of RPA foci per nucleus in leptotene and zygotene oocytes from 16 dpc wild-type (+/+ or *Top6bl ^+/Δ17Ct^*), *Top6bl^W562A/W562A^*, *Top6bl ^Δ17Ct/Δ17Ct^*, and *Top6bl^-/-^*mice (n=4 wild-type and n=2 mice per mutant genotype). Number of nuclei per genotype at leptonema (157, 85, 14, and 29) and at zygonema (165, 55, 105, and 28). Grey bars show the mean value. P values were determined using the two-tailed unpaired Mann-Whitney test. Ratios of the wild-type and mutant mean values are shown.

**d.** Immunostaining of MEI4, SYCP3 and γH2AX in oocytes from 16 dpc wild-type (+/+), *Top6bl^W562A/W562A^*, and *Top6bl ^Δ17Ct/Δ17Ct^* ovaries. Scale bar, 10μm.

**e.** Quantification of axis-associated MEI4 foci in leptotene and zygotene oocytes from 15 and 16 dpc wild-type (+/+ or *Top6bl ^+/Δ17Ct^*), *Top6bl^W562A/W562A^*, and *Top6bl ^Δ17Ct/Δ17Ct^* mice (n=3 wild-type, n=1 *Top6bl^W562A/W562A^*, and n=2 *Top6bl ^Δ17Ct/Δ17Ct^* mice). Number of nuclei per genotype at leptonema (139, 91, and 42) and at zygonema (106, 24, and 23). Grey bars show the mean values. P values were determined using the two-tailed unpaired Mann-Whitney test.

**f.** Quantification of axis-associated REC114 foci in leptotene and zygotene oocytes from 16 dpc wild-type (+/+) and *Top6bl^-/-^*mice (n=1 mouse/genotype). Number of nuclei: 29 and 36 L, and 31and 29 Z, respectively. Grey bars show the mean values. P values were determined using the two-tailed unpaired Mann-Whitney test.

**Extended data Fig. 4 a.** Immunostaining of ANKRD31, SYCP3 and γH2AX in oocytes from 16 dpc wild-type (+/+), *Top6bl^W562A/W562A^*, and *Top6bl ^Δ17Ct/Δ17Ct^* ovaries. Scale bar, 10μm.

**b.** Quantification of axis-associated ANKRD31 foci in leptotene and zygotene oocytes from 16 and 18 dpc wild-type (+/+ or *Top6bl ^+/Δ17Ct^*), *Top6bl^W562A/W562A^*, and *Top6bl ^Δ17Ct/Δ17Ct^* mice (n=3 wild-type, n=1 *Top6bl^W562A/W562A^*, and n=2 *Top6bl ^Δ17Ct/Δ17Ct^* mice). Number of nuclei per genotype at leptonema (21, 74, and 56) and at zygonema (56, 57, and 108). Grey bars show the mean values. P values were determined using the two-tailed unpaired Mann-Whitney test.

**Extended data Fig. 5**

a. Quantification of primordial, primary, and growing follicles in ovaries from 20 dpp wild-type homozygous or heterozygous (+/+, *Top6bl ^+/Δ17Ct^*, *Top6bl ^+/W562A^*, +/ *Top6bl^-^*), *Top6bl^W562A/W562A^, Top6bl ^Δ17Ct/Δ17Ct^*, and *Top6bl^-/-^* mice. Each data point is the mean ± SD of at least 10 sections from one ovary. Number of mice: 6, 2, 4, and 2, respectively. P values were determined using the two-tailed unpaired Mann-Whitney test. Asterisks indicate the p values (see Methods).

b. Quantification of primordial, primary, and growing follicles in ovaries from 7-8-week-old wild-type (+/+ or *Top6bl ^+/W562A^*), *Top6bl^W562A/W562A^, Top6bl ^Δ17Ct/Δ17Ct^*, and *Top6bl^-/-^* mice. Each data point is the mean ± SD of 20 sections from one ovary. P values were determined using the two-tailed unpaired Mann-Whitney tests. Number of mice: 7, 5, 3, and 2 respectively.

c. Fertility tests*. Top6bl ^+/Δ17Ct^*, *Top6bl^W562A/W562A^*and *Top6bl ^Δ17Ct/Δ17Ct^* females (n= 5, 4 and 4 respectively), from 2.3 to 10.6 months of age, were mated with heterozygous males (*Top6bl ^+/Δ17Ct^* or *Top6bl ^+/W562A^*). Each symbol refers to a female. Only two of the four *Top6bl^W562A/W562A^* females were fertile (large circle, dark green, and small circle), and after the last progeny, they were not mated any longer. Breeding periods of non-productive females and for *Top6bl ^+/Δ17Ct^* females are shown.

d. Haematoxylin and eosin-stained sections of ovaries from 20 dpp wild-type (+/+), *Top6bl^W562A/W562A^* and *Top6bl ^Δ17Ct/Δ17Ct^* mice. Scale bars, 250 and 50 μm in the left and right panel, respectively. Primordial (PriF), primary (PF), and growing (GF) follicles are indicated.

e. Sperm count in *Top6bl* mutants. Sperm in epididymis was counted in 2-month-old mice. Each data point is one epididymis (number of mice= 7, 5, 3 and 1 for wild-type (+/+ or +/-), *Top6bl^W562A/W562A^*, *Top6bl ^Δ17Ct/Δ17Ct^*, and *Top6bl*^-/-^, respectively). The mean ± SD is shown. P values were determined using the two-tailed unpaired Mann-Whitney test.

f. Periodic acid-Schiff staining of testis sections from 2 month-old wild-type (+/+), *Top6bl^W562A/W562A^*, and *Top6bl ^Δ17Ct/Δ17Ct^* mice. Scale bar, 250μm. Zoomed images of stage XII tubules are shown on the right.

g. Testis weight in *Top6bl* mutants. Testis weight was measured in 2 month-old mice. Each data point is one testis (number of mice= 5, 5, 3, and 1 for wild-type (+/+), *Top6bl^W562A/W562A^*, *Top6bl ^Δ17Ct/Δ17Ct^,* and *Top6bl*^-/-^, respectively). The mean value ± SD is shown. P values were determined using the two-tailed unpaired Mann-Whitney test.

h. Staging of tubules in testis sections from 2-month-old wild-type and *Top6bl ^Δ17Ct/Δ17Ct^* mice (n=2 mice/genotype). Tubules were divided in five groups based on several criteria. I-VI: round spermatid with acrosomes, 0 to ∼25% of the perimeter of the spermatid, elongated spermatids on the lumen; VII-VIII: round spermatids with acrosomes, ∼50% of the perimeter; IX-X: elongated spermatids with acrosomes not hook-shaped; XI: elongated spermatids with a hook; XII: presence of metaphases. Number of staged tubules: 310 in wild-type and 366 in *Top6bl ^Δ17Ct/Δ17Ct^* mice. The horizontal bars show the mean value. The distributions are statistically different (Pearson’s Chi-Square: 36.028).

i. Detection of apoptosis and metaphase in testis sections. Sections were labelled with TUNEL (green) and H3pS10 (red), and stained with DAPI (blue). Scale bar, 100μm. White arrowheads indicate tubules positive for H3pS10 and TUNEL.

j. A stage XII tubule with metaphase spermatocytes (from *Top6bl ^Δ17Ct/Δ17Ct^* mice). Scale bar, 100μm.

Several metaphases with condensed chromosomes are seen, and one (arrow) shows relatively more intense nuclear staining, suggesting apoptosis.

k. Detection of apoptosis by TUNEL assay. Tubules were classified in three categories: <3, 3 to 5, and >5 apoptotic nuclei. The mean values ± SD of the 3 to 5 and >5 categories are shown. Number of 2 month-old mice: 4 wild-type (+/+ and *Top6bl ^+/Δ17Ct^*) and 2 mice per mutant. The percentage of tubules with <3, 3 to 5, and >5 apoptotic nuclei was significatively different between wild-type and *Top6bl^W562A/W562A^* and *Top6bl^Δ17Ct /Δ17Ct^* mice (Chi-square Pearson: 354.03 and 811.63, respectively).

**Extended data Fig. 6 a.** Identification of spermatocytes at early, mid, late leptonema and zygonema. Examples of wild-type nuclei at the different stages by staining with SYCP3 (green), SYCP1 (red) and γH2AX (magenta). Scale bar, 10μm.

**b.** Quantification of axis-associated RPA2 foci in early/mid and late leptotene, zygotene and pachytene spermatocytes from 12-14dpp wild-type (+/+ or *Top6bl ^+/Δ17Ct^*; n=3), *Top6bl^W562A/W562A^* (n=2), and *Top6bl^Δ17Ct /Δ17Ct^* (n=2) mice. Number of nuclei per genotype at early/mid leptotene (105, 36, and 41), late leptotene (91, 38, and 131), zygotene (135, 62, and 85), and pachytene (178, 119, and 80). Grey bars show the mean values. P values were determined using the two-tailed unpaired Mann-Whitney test.

**c.** RPA2 focus variation during prophase. For each genotype (wild-type, *Top6bl^W562A/W562A^*, and *Top6bl ^Δ17Ct/Δ17Ct^*), the number of RPA2 foci at the different stages was normalized to the mean number at zygonema (set to 1). The mean value ± SD is shown. Statistical significance between wild-type and each mutant (*Top6bl^W562A/W562A^*; *Top6bl ^Δ17Ct/Δ17Ct^*) was tested at late leptonema using the two-tailed unpaired Mann-Whitney test.

**d.** Immunostaining of RPA2, SYCP3 and γH2AX in spermatocytes from 12-14dpp wild-type (+/+), *Top6bl^W562A/W562A^*, and *Top6bl ^Δ17Ct/Δ17Ct^* spermatocytes. A white dotted circle highlights the unsynapsed X and Y chromosomes at pachynema (SYCP3 staining) in *Top6bl ^Δ17Ct/Δ17Ct^* mice. Scale bar, 10μm.

**Extended data Fig. 7 a.** Immunostaining of ANKRD31 and SYCP3 in spermatocytes from 12-14dpp wild-type (+/+), *Top6bl^W562A/W562A^*, and *Top6bl ^Δ17Ct/Δ17Ct^* mice. At zygonema, dotted white circles (merge panels) highlight ANKRD31 aggregates at the ends of some chromosome axes. Scale bar, 10μm.

**b.** Quantification of axis-associated MEI4 foci in early/mid, late leptotene and zygotene spermatocytes from 12-14 dpp wild-type (+/+ or *Top6bl ^+/Δ17Ct^*; n=2), *Top6bl^W562A/W562A^* (n=1), and *Top6bl ^Δ17Ct/Δ17Ct^* (n=2) mice. Number of nuclei: 97 early/mid L, 43 late L, 114 Z in wild-type; 26 early/mid L, 28 late L and 16 Z in *Top6bl^W562A/W562A^*; 77 early/mid L, 68 late L and 103 Z in *Top6bl ^Δ17Ct/Δ17Ct^*. Grey bars show the mean values. P values were determined using the two-tailed unpaired Mann-Whitney test.

**c.** Immunostaining of MEI4 and SYCP3 in spermatocytes from 12-14 dpp wild-type (+/+), *Top6bl^W562A/W562A^*, and *Top6bl ^Δ17Ct/Δ17Ct^* mice. Scale bar, 10μm.

**Extended data Fig. 8 a.** Overlap between wild-type and *Top6bl^Δ17Ct/Δ17Ct^* hotspots and the default hotspots used in the *PRDM9^KO^* mouse strain. The overlaps were restricted to the central 400bp around the estimated hotspot centre.

**b.** DMC1-SSDS signal (heatmap and average plots) from wild-type and *Top6bl^Δ17Ct/Δ17Ct^* mice at PRDM9^B6^-, PRDM9^RJ2^- and PRDM9^KO^-defined hotspots.

**Extended data Fig. 9 a.** H3K4me3 signal (heatmap and average plots) from B6 and RJ2 mice and DMC1-SSDS signal from *Top6bl^Δ17Ct/Δ17Ct^* and wild-type mice at wild-type hotspots (13562) and at hotspots specifically activated in *Top6bl^Δ17Ct/Δ17Ct^* (3519). Peaks are ranked in function of the H3K4me3 or DMC1 signal intensity.

**b.** DMC1-SSDS signal in *Top6bl ^Δ17Ct/Δ17Ct^*, *Ankrd31^-/-^* and their respective wild-type control mice. Signal is represented as boxplots computed for hotspots ranked according to increased DMC1-SSDS signal intensity and grouped into 10 bins (1 bin = 10% of the total hotspot number; bin 1 = weakest signal, bin 10= strongest signal).

**c.** Control for the sub-telomeric effect. Quantification was done as in Fig. 6f, but with ratios calculated between two random pools of 10 non-sub-telomeric consecutive hotspots.

**d.** Sub-telomeric/Non-sub-telomeric hotspot density ratios for unchanged, decreased, and increased hotspots, as defined by DESeq2 analysis, for sub-telomeric regions defined using various distances from the telomeric annotated ends (1-2-3-4-5-7-10Mb). The log2 ratio was plotted and showed excess (positive values) or lack (negative values) of hotspots at sub-telomeric regions compared with non-sub-telomeric regions.

**e.** Distribution of hotspots from annotated telomeric ends and according to their DESeq2 category (unchanged, decreased, or low signal) along the 3Mb window. Note the absence of increased hotspots within this sub-telomeric window size. The normalized wild-type DMC1-SSDS signal is shown on the y-axis.

**Extended data Fig. 10**

Browser windows of DMC1-SSDS in wild-type (blue) and *Top6bl^Δ17Ct/Δ17Ct^* (green) mice within the 10Mb q-arm telomere-proximal regions of chromosomes 1 to 10.

**Extended data Fig. 11**

Browser windows of DMC1-SSDS in wild-type (blue) and *Top6bl^Δ17Ct/Δ17Ct^* (green) mice within the 10Mb q-arm telomere-proximal regions of chromosomes 11 to X.

**Extended data Table 1**

Data collection and refinement statistics.

**Extended data Table 2.**

Statistics to test the biased distribution of hotspot densities at sub-telomeric vs. non-sub-telomeric regions in *Top6bl ^Δ17Ct^*samples: Pearson’s Chi square adjusted p-value for tests considering two (unchanged/decreased) or three (unchanged/decreased/increased) hotspot categories for different sizes of sub-telomeric regions. NA: not applicable due to the absence of hotspot with quantifiable signal.

**Extended data Table 3**

Oligonucleotides

**Extended data Table 4**

Percentage of pachytene spermatocytes with full synapsis

