## Supplementary material for "TOPOVIBL-REC114 interaction regulates meiotic DNA double-strand breaks": Nore_Juarez_supmarerial

**Supplementary Materials**

**Supplementary Figures (1-17)**

**Supplementary Tables (S1, S2, S3 and S4)**

**Supplementary discussion**

**Supplementary references**

**Supplementary Fig. 1**

**a.** Yeast two-hybrid assays between full-length TOPOVIBL and truncated REC114 variants and western blots to confirm TOPOVIBL and REC114 expression. GAD: GAL4 activation domain. GBD: GAL4 DNA binding domain. Growth was tested on -LW, -LWH, -LWH+5mM 3AT and -LWHA medium. 10µl of cells at the indicated concentration were spotted. Western blotting was performed with an anti-GDB (aGBD) or anti-GAD antibody (aGAD).

**b.** Yeast two-hybrid assays between full-length TOPOVIBL and the GAL4 activating domain, and between truncated REC114 variants and the GAL4 binding domain, with western blot analysis to confirm TOPOVIBL and REC114 expression.

**c.** Yeast two-hybrid assays between truncated TOPOVIBL variants and full-length REC114 with western blot analysis to confirm TOPOVIBL and REC114 expression

**d.** Yeast two-hybrid assays between truncated TOPOVIBL variants and the GAL4 activating domain with western blot analysis to confirm TOPOVIBL expression.

**e.** Yeast two-hybrid assays between TOPOVIBL-ΔC130/ΔC29 and full-length REC114 or GAL4 activating domain with western blot analysis to confirm TOPOVIBL and REC114 expression.

**f.** Yeast two-hybrid assays between truncated TOPOVIBL variants and SPO11β with western blot analysis to confirm TOPOVIBL and SPO11β expression.

**Supplementary Fig. 2**

**a.** SDS-PAGE analysis of fractions 1-12 of the gel filtration elution profiles of REC114 15-159, TOPOVIBL 450-579, and their complex. Proteins were expressed individually and purified on Ni2+ resin.

**b.** SDS-PAGE analysis of fractions 1-12 of the elution profiles of FL-REC114 bound to MEI4 1-27, TOPOVIBL 450-579, and their complex. REC114 and MEI4 we co-expressed and purified on Ni2+ resin.

**Supplementary Fig. 3**

Prediction of mouse TOPOVIBL secondary structure using PSIPRED [49].

**Supplementary Fig. 4**

Comparison of the S200 gel filtration elution profiles of wild-type TOPOVIBL 450-579 and the W562A mutant. The two profiles are essentially identical, indicating that the W562A mutation does not affect the overall structure of this TOPOVIBL fragment.

**Supplementary Fig. 5**

**a.** Yeast two-hybrid assays between TOPOVIBL point mutants and full length REC114 with western blot analysis to confirm TOPOVIBL and REC114 expression

**b.** Yeast two-hybrid assays between TOPOVIBL point mutants and Gal4-activating domain with western blot analysis to confirm TOPOVIBL expression.

**c.** Yeast two-hybrid assays between TOPOVIBL point mutants and SPO11β with western blot analysis to confirm TOPOVIBL and SPO11β expression.

**Supplementary Fig. 6**

**a-c**. Comparison of the REC114-TOPOVIBL complex structure (**a**) with the ANKRD31-REC114 (**b**) and EVL-ActA [50] (**c**) structures.

**f.** Expression of *Top6bl* assessed by RT-PCR. RNA extracted from ovaries of 16 dpc *Top6bl* $^{\Delta17Ct/\Delta17Ct}$ or *Top6bl* $^{+/\Delta17Ct}$ mice was incubated with or without reverse transcriptase (RT), amplified by PCR and incubated with or without *Eci*I restriction enzyme. The size of the fragments after *Eci*I digestion are shown for the *Top6bl* $^{+}$ and *Top6bl* $^{\Delta17Ct}$ alleles. Stars indicate non-specific amplification products. Control PCR assays for monitoring the expression of the two *Spo11* splice variants (*Spo11*$\alpha$ and $\beta$) and of actin are shown.

**Supplementary Fig. 8**

**a.** Quantification of γH2AX signal intensity in leptotene (L)/zygotene (Z) oocytes from 16 and 18 dpc wild-type and 18dpc *Top6bl* $^{\Delta17Ct/\Delta17Ct}$ mice. Number of nuclei: 64, 9, and 167, respectively. Grey bars show the mean values. P values were determined using the two-tailed unpaired Mann-Whitney test. Progression into prophase at 18dpc was: L 46%, Z 54%, Pachynema (P) 0% in *Top6bl* $^{\Delta17Ct/\Delta17Ct}$, and L 2%, Z 8%, P 90% in wild-type oocytes.

**b.** Quantification of axis-associated REC114 foci in leptotene and zygotene oocytes from 16 dpc wild-type (+/+) and *Top6bl*$^{-/-}$ mice (n=1 mouse/genotype). Number of nuclei: 29 and 36 L, and 31and 29 Z, respectively. Grey bars show the mean values. P values were determined using the two-tailed unpaired Mann-Whitney test.

**Supplementary Fig. 9**

Immunostaining of IHO1 and SYCP3 in oocytes from 16 dpc wild-type (+/-) (**a**), *Top6bl*$^{W562A/W562A}$(**b**), *Top6bl* $^{\Delta17Ct/\Delta17Ct}$ (**c**), and *Top6bl*$^{-/-}$ (**d**) mice. Scale bar, 10µm.

**Supplementary Fig. 10**

**a.** Quantification of primordial, primary, and growing follicles in ovaries from 20 dpp wild-type homozygous or heterozygous (+/+, *Top6bl* $^{+/\Delta17Ct}$, *Top6bl* $^{+/W562A}$, +/ *Top6bl*$^{-}$), *Top6bl*$^{W562A/W562A}$, *Top6bl* $^{\Delta17Ct/\Delta17Ct}$, and *Top6bl*$^{-/-}$ mice. Each data point is the mean ± SD of at least 10 sections from one ovary. Number of mice: 6, 2, 4, and 2, respectively. P values were determined using the two-tailed unpaired Mann-Whitney test. Asterisks indicate the p values (see Methods).

**b.** Haematoxylin and eosin-stained sections of ovaries from 20 dpp wild-type (+/+), *Top6bl*$^{W562A/W562A}$ and *Top6bl* $^{\Delta17Ct/\Delta17Ct}$ mice. Scale bars, 250 and 50 µm in the left and right panel, respectively. Primordial (PriF), primary (PF), and growing (GF) follicles are indicated.

**c.** Quantification of primordial, primary, and growing follicles in ovaries from 7-8-week-old wild-type (+/+ or *Top6bl* $^{+/W562A}$), *Top6bl*$^{W562A/W562A}$, *Top6bl* $^{\Delta17Ct/\Delta17Ct}$, and *Top6bl*$^{-/-}$ mice. Each data point is the mean ± SD of 20 sections from one ovary. P values were determined using the two-tailed unpaired Mann-Whitney tests. Number of mice: 7, 5, 3, and 2 respectively.

**d.** Fertility tests. *Top6bl* $^{+/\Delta17Ct}$, *Top6bl*$^{W562A/W562A}$ and *Top6bl* $^{\Delta17Ct/\Delta17Ct}$ females (n= 5, 4 and 4 respectively), from 2.3 to 10.6 months of age, were mated with heterozygous males (*Top6bl* $^{+/\Delta17Ct}$ or *Top6bl* $^{+/W562A}$). Each symbol refers to a female. Only two of the four *Top6bl*$^{W562A/W562A}$ females were fertile (large circle, dark green, and small circle), and after the last progeny, they were not mated any longer. Breeding periods of non-productive females and for *Top6bl* $^{+/\Delta17Ct}$ females are shown.

**Supplementary Fig. 11**

**a.** Quantification of axis-associated MEI4 foci in early/mid, late leptotene and zygotene spermatocytes from 12-14 dpp wild-type (+/+ or *Top6bl* $^{+/\Delta17Ct}$ ; n=2), *Top6bl*$^{W562A/W562A}$ (n=1), and *Top6bl* $^{\Delta17Ct/\Delta17Ct}$ (n=2) mice. Number of nuclei: 97 early/mid L, 43 late L, 114 Z in wild-type; 26 early/mid L, 28 late L and 16 Z in *Top6bl*$^{W562A/W562A}$; 77 early/mid L, 68 late L and 103 Z in *Top6bl* $^{\Delta17Ct/\Delta17Ct}$. Grey bars show the mean values. P values were determined using the two-tailed unpaired Mann-Whitney test.

**b.** Immunostaining of MEI4 and SYCP3 in spermatocytes from 12-14 dpp wild-type (+/+), *Top6bl*$^{W562A/W562A}$, and *Top6bl* $^{\Delta17Ct/\Delta17Ct}$ mice. Scale bar, 10µm.

**Supplementary Fig. 12**

**a.** Overlap between wild-type and *Top6bl*$^{\Delta17Ct/\Delta17Ct}$ hotspots and the default hotspots used in the *PRDM9*$^{KO}$ mouse strain. The overlaps were restricted to the central 400bp around the estimated hotspot centre.

**b.** DMC1-SSDS signal is not changed in the 50Mb adjacent to p-arm telomeres in *Top6bl*$^{\Delta 17Ct/\Delta 17Ct}$ samples relative to wild-type samples. The DMC1-SSDS signal ratio within hotspots (log2-fold change estimated by DESeq2) was computed over 5Mb-windows with a 1Mb-step. Values were averaged for autosomes and plotted separately for the X chromosome along the 50Mb. As the genomic DNA sequence of p-arms has not been assembled, it is immediately flanked by centromeric and q-arm DNA sequences, where the signal can be mapped and quantified.

**c.** Control for the sub-telomeric effect. Quantification was done as in Fig. 3h, but with ratios calculated between two random pools of 10 non-sub-telomeric consecutive hotspots.

**d.** H3K4me3 signal (heatmap and average plots) from B6 and RJ2 mice and DMC1-SSDS signal from *Top6bl*$^{\Delta 17Ct/\Delta 17Ct}$ and wild-type mice at wild-type hotspots (13562) and at hotspots specifically activated in *Top6bl*$^{\Delta 17Ct/\Delta 17Ct}$ (3519). Peaks are ranked in function of the H3K4me3 or DMC1 signal intensity.

**e.** DMC1-SSDS signal in *Top6bl*$^{\Delta 17Ct/\Delta 17Ct}$, *Ankrd31*$^{-/-}$ and their respective wild-type control mice. Signal is represented as boxplots computed for hotspots ranked according to increased DMC1-SSDS signal intensity and grouped into 10 bins (1 bin = 10% of the total hotspot number; bin 1 = weakest signal, bin 10= strongest signal).

**f.** Distribution of hotspots from annotated telomeric ends and according to their DESeq2 category (unchanged, decreased, or low signal) along the 3Mb window. Note the absence of increased hotspots within this sub-telomeric window size. The normalized wild-type DMC1-SSDS signal is shown on the y-axis.

**Supplementary Fig. 13**

Browser windows of DMC1-SSDS in wild-type (blue) and *Top6bl*$^{\Delta 17Ct/\Delta 17Ct}$ (green) mice within the 10Mb q-arm telomere-proximal regions of chromosomes 1 to 10.

**Supplementary Fig. 14**

Browser windows of DMC1-SSDS in wild-type (blue) and *Top6bl*$^{\Delta 17Ct/\Delta 17Ct}$ (green) mice within the 10Mb q-arm telomere-proximal regions of chromosomes 11 to X.

**Supplementary Fig. 15**

**a.** Sperm count in *Top6bl* mutants. Sperm in epididymis was counted in 2-month-old mice. Each data point is one epididymis (number of mice= 7, 5, 3 and 1 for wild-type (+/+ or +/-), *Top6bl*$^{W562A/W562A}$, *Top6bl*$^{\Delta 17Ct/\Delta 17Ct}$, and *Top6bl*$^{-/-}$, respectively). The mean ± SD is shown. P values were determined using the two-tailed unpaired Mann-Whitney test.

**b.** Periodic acid-Schiff staining of testis sections from 2 month-old wild-type (+/+), *Top6bl*$^{W562A/W562A}$, and *Top6bl*$^{\Delta 17Ct/\Delta 17Ct}$ mice. Scale bar, 250µm. Zoomed images of stage XII tubules are shown on the right.

**c.** Testis weight in *Top6bl* mutants. Testis weight was measured in 2 month-old mice. Each data point is one testis (number of mice= 5, 5, 3, and 1 for wild-type (+/+), *Top6bl*$^{W562A/W562A}$, *Top6bl* $^{\Delta 17Ct/\Delta 17Ct}$, and *Top6bl*$^{-/-}$, respectively). The mean value ± SD is shown. P values were determined using the two-tailed unpaired Mann-Whitney test.

**d.** Staging of tubules in testis sections from 2-month-old wild-type and *Top6bl* $^{\Delta 17Ct/\Delta 17Ct}$ mice (n=2 mice/genotype). Tubules were divided in five groups based on several criteria. I-VI: round spermatid with acrosomes, 0 to ~25% of the perimeter of the spermatid, elongated spermatids on the lumen; VII-VIII: round spermatids with acrosomes, ~50% of the perimeter; IX-X: elongated spermatids with acrosomes not hook-shaped; XI: elongated spermatids with an hook; XII: presence of metaphases. Number of staged tubules: 310 in wild-type and 366 in *Top6bl* $^{\Delta 17Ct/\Delta 17Ct}$ mice. The horizontal bars show the mean value. The distributions are statistically different (Pearson's Chi-Square: 36.028).

**e.** Detection of apoptosis and metaphase in testis sections. Sections were labelled with TUNEL (green) and H3pS10 (red), and stained with DAPI (blue). Scale bar, 100μm. White arrowheads indicate tubules positive for H3pS10 and TUNEL.

**f.** A stage XII tubule with metaphase spermatocytes (from *Top6bl* $^{\Delta 17Ct/\Delta 17Ct}$ mice). Scale bar, 100μm. Several metaphases with condensed chromosomes are seen, and one (arrow) shows relatively more intense nuclear staining, suggesting apoptosis.

**g.** Detection of apoptosis by TUNEL assay. Tubules were classified in three categories: <3, 3 to 5, and >5 apoptotic nuclei. The mean values ± SD of the 3 to 5 and >5 categories are shown. Number of 2 month-old mice: 4 wild-type (+/+ and *Top6bl* $^{+/\Delta 17Ct}$) and 2 mice per mutant. The percentage of tubules with <3, 3 to 5, and >5 apoptotic nuclei was significatively different between wild-type and *Top6bl*$^{W562A/W562A}$ and *Top6bl* $^{\Delta 17Ct/\Delta 17Ct}$ mice (Chi-square Pearson: 354.03 and 811.63, respectively).

**Supplementary Fig. 16**

Identification of spermatocytes at early, mid, late leptonema and zygonema. Examples of wild-type nuclei at the different stages by staining with SYCP3 (green), SYCP1 (red) and γH2AX (magenta). Scale bar, 10μm.

**Supplementary Fig. 17**

**a.** DMC1-SSDS signal correlation between wild-type and *Ankrd31*$^{-/-}$ mice at wild-type hotspots. The type 1 single-strand DNA signal was library-normalized (fragment per million). The Spearman rho and associated p-value are shown. Ten telomere-proximal hotspots are highlighted for each autosome (blue) and for chromosome X (red). The dotted line represents the x=y relationship.

**b.** MA plot of DMC1-SSDS signal change in *Ankrd31*$^{-/-}$ compared to wild-type samples. The log2 fold changes were estimated using DESeq2 for differential analysis and after log fold change shrinkage. Significantly increased and decreased hotpots are highlighted in red (n=3111) and green (n=3224), respectively (adjusted p-value<0.1). Unchanged hotspots are shown in grey (n=7145). The mean normalized count corresponds to the baseMean value from DESeq2 analysis.

**c.** On the left panel, the q-arm sub-telomeric relative effect in *Ankrd31*$^{-/-}$ mice was measured for each chromosome as follows: the averaged DMC1-SSDS signal ratio (*Ankrd31*$^{-/-}$/wild-type) of the last ten hotspots of a given chromosome was compared to the averaged DMC1-SSDS signal ratio of 10 non-telomeric randomly chosen consecutive hotspots in the same chromosome. Boxplots represents the log2 fold-change between these values (sub-telomeric/non-sub-telomeric) for 10 randomizations. The control of this measure is shown on the right panel and was calculated using two non-sub-telomeric sets of 10 consecutive hotspots.

**d.** Decreased (green), increased (red), and unchanged (grey) hotspot density within 3Mb sub-telomeric regions relative to non-sub-telomeric regions. Decreased, increased and unchanged hotspots were determined by DESeq2 analysis. The densities of decreased vs unchanged hotspots in the two regions were compared using the Pearson's Chi-square test, and p-values were adjusted for multiple testing using the Benjamini & Yekutieli method. Yellow stars indicate p-value <0.05. The Chi square test results are provided in Supplementary Table S3. See Supplementary discussion 2 for comparison with DMC1 SSDS analysis in *Top6bl*$^{\Delta 17Ct/\Delta 17Ct}$ mice.

**Supplementary Table S1**. Data collection and refinement statistics.

**Supplementary Table S2**. Statistics to test the biased distribution of hotspot densities at sub-telomeric vs. non-sub-telomeric regions in *Top6bl*$^{\Delta 17Ct}$ samples: Pearson's Chi square adjusted p-value for tests considering two (unchanged/decreased) or three (unchanged/decreased/increased) hotspot categories for different sizes of sub-telomeric regions.

**Supplementary Table S3**. Statistics to test the biased distribution of hotspot densities at sub-telomeric vs. non-sub-telomeric regions in *Ankrd31*$^{-/-}$ samples: Pearson's Chi square adjusted p-value for tests considering two (unchanged/decreased) or three (unchanged/decreased/increased) hotspot categories for different sizes of sub-telomeric regions.

**Supplementary Table S4**. Oligonucleotides

#### Supplementary discussion

##### 1. Interaction of TOPOVIBL and ANKRD31 to the PH domain of REC114.

*In vitro* binding assays show a mutually exclusive interaction of REC114 PH domain and either TOPOVIBL or ANKRD31 (Extended Data Fig. 2i). The crystal structure of the ANKRD31-REC114 complex [1] shows that the ANKRD31 interacting fragment (45 residues) is significantly longer than that of TOPOVIBL and covers a larger surface on REC114 (~1800 Å2) packing against both of its β-sheets (Supplementary Fig. 6a, b). The C-terminal part of the ANKRD31 peptide forms two helices and packs against the same surface as TOPOVIBL, and both peptides interact with equivalent REC114 residues (Extended Data Fig. 2h). Mutation of REC114 L104 reduced the binding to both ANKRD31[1] and to

TOPOVIBL (Fig. 1i, lane 8). Similarly, the ANKRD31 W1842A mutation disrupted the interaction with REC114[1], as did the corresponding W562A mutation in TOPOVIBL (Fig. 1i, lane 8). As in our hands SUMO or MBP fusion ANKRD31 1808-1857 proteins aggregated unless bound to REC114, we could not determine its Kd for REC114. All these properties suggest a competition for binding to the PH domain of REC114 *in vitro*, with the caveat that these conclusions are based on interactions between peptides, not full-length proteins. Further implications of the relevance *in vivo*, would require analyzing protein levels and to determine which components are limiting.

**2. Comparison of $Top6bl^{\Delta17Ct/\Delta17Ct}$ with $Ankrd31^{-/-}$ mice.**

*In vivo*, insight into the role of ANKRD31 was obtained from analyzing $Ankrd31^{-/-}$ mice [1,2]. These mice have multiple defects in DSB activity, some of which are discussed below, and are interesting to analyze as some are apparently similar to those we reported in *Top6bl* mutant mice. One striking similarity is the XY chromosome DSB and synapsis defect.

In $Ankrd31^{-/-}$ mice this phenotype is explained by the reduction of IHO1/REC114/MEI4/ANKRD31 aggregates on the PAR. This phenotype was not observed in *Top6bl* mutants which do not have any detectable defect in the formation of those aggregates (Extended Data Fig. 7). Despite this cytological difference, it is possible that in both mutant contexts the reduced DSB activity in the PAR is due to a reduction or loss of the interaction between REC114 and TOPOVIBL.

In $Ankrd31^{-/-}$ mice, the DMC1-SSDS signal is decreased not only in the PAR but also at the ends of chromosomes 4, 9 and 13 [1,2]. The four regions are characterized by the presence of the mo-2 minisatellite which may either contribute to the phenotype or be a consequence of other properties of these chromosome ends [3]. In $Top6bl^{\Delta17Ct/\Delta17Ct}$ mice, the DMC1-SSDS signal was also decreased at the ends of chr. 4, 9 and 13 extending the similarity of this phenotype between the two mutants (Fig. 3h, Extended data Fig. 8c, d, Supplementary Fig. 13, 14). As we showed in $Top6bl^{\Delta17Ct/\Delta17Ct}$ mice a reduction of DSB activity at most chromosome ends (Fig. 3h), we then examined DSB activity in $Ankrd31^{-/-}$ mice at all chromosome ends. This analysis revealed a reduction of DSB activity at most chromosome ends in $Ankrd31^{-/-}$: Using the same method as for $Top6bl^{\Delta17Ct/\Delta17Ct}$ analysis, based on DeSeq2, a near systematic reduction of DSB activity was detected at the ten most telomere-proximal hotspots in $Ankrd31^{-/-}$ mice (Supplementary Fig. 17a, c). Overall, the DMC1-SSDS signal was reduced in the 3Mb subtelomeric region of several chromosomes, with a significant difference for chromosomes 2, 4, 6, 9, 10, 11, 13, 15 and X (Supplementary Fig. 17d).

Quantitatively these phenotypes of reduced DSB activity in subtelomeric regions were not always identical in $Ankrd31^{-/-}$ and $Top6bl^{\Delta17Ct/\Delta17Ct}$ however, as some chromosomes showed significant effects in both mutants (4, 9, 10, 11, 13, X), and some only in one of the two mutants (3, 6, 8, 17, 19) (Extended Data Fig. 8 and Supplementary Fig. 7). Two distinct scenarios can be proposed: either the cause of the DSB activity reduction is distinct in $Ankrd31^{-/-}$ and $Top6bl^{\Delta17Ct/\Delta17Ct}$, or it is due to a common defect, the lack of interaction between TOPOVIBL and REC114, but with additional defects in *Ankrd31*$^{-/-}$ mice. These mice have indeed several additional DSB defects along chromosome arms with increased or decreased DMC1 SSDS signal (Supplementary Fig. 17b) [2] and with use of default (PRDM9-independent) sites. These defects are detected genome-wide and could also contribute to alterations of DSB activity near chromosome ends.

##### 3. Increased activity at weak hotspots in *Top6bl*$^{\Delta17Ct/\Delta17Ct}$ mice

A subset of very weak hotspots with slightly increased signal was detected in *Top6bl*$^{\Delta17Ct/\Delta17Ct}$ mice leading to an additional 3519 peaks not detected as peaks in wild-type (Fig. 3d). At these weak hotspots, we however detected a weak enrichment for DMC1 in wild-type samples, ~1.7-fold lower than in *Top6bl*$^{\Delta17Ct/\Delta17Ct}$ mice (Supplementary Fig. 12d). We conclude that these 3519 peaks are weak PRDM9-dependent hotspots detected as significantly enriched regions (*i.e.* peaks) in *Top6bl*$^{\Delta17Ct/\Delta17Ct}$ but not in wild-type samples. PRDM9$^{B6}$-specific H3K4me3 enrichment, evaluated in wild-type mice, also indicated a weak PRDM9 activity at these sites (Supplementary Fig. 12d). We asked if we could also detect any increase of intensity in *Top6bl*$^{\Delta17Ct/\Delta17Ct}$ at the weakest hotspots identified in wild-type, but this was not the case (bin 1 from Supplementary Fig. 12e). Therefore, the increased DMC1-SSDS signal could be due to the specifically low DSB activity of the 3519 *Top6bl*$^{\Delta17Ct/\Delta17Ct}$ -specific hotspots or to another specific property of these sites. As we did not identify any other specific (genomic, epigenetic) feature at these hotspots, we favor the hypothesis, that this phenotype is related to the genome-wide delay in DSB activity. Due to the down-regulation of DSB activity by homolog engagement, such a delay may give the opportunity for normally late DSBs to form at an increased level[4].

##### 4. Features of subtelomeric regions.

One unique property of telomeres is their attachment to the nuclear envelope [5,6]. Their nuclear attachment or functionally related events may have consequences on DSB activity regulation. If this property was related to the decreased DSB activity described above, both telomeric ends should behave similarly in *Top6bl*$^{\Delta17Ct/\Delta17Ct}$. However, due to the proximity of the centromere, information about the DSB activity adjacent to the p-arm telomere is inaccessible in DMC1-SSDS data due to the presence of repeated DNA.

Several studies have described the specific properties of chromosome ends, specifically in the context of meiotic DSB activity. In *S. cerevisiae*, persistence of DSB potential, linked to persistence of Hop1, is a manifestation of a telomere end effect [7]. It is not known whether this property applies to the mouse as well, and whether it could be linked to the observed phenotype. We did not detect in databases epigenetic modifications features specific to the 1-3Mb subtelomeric region. Interesting observations potentially linked with the phenotype of *Top6bl*$^{\Delta17Ct/\Delta17Ct}$ mice have been reported:

First, a specific decrease of the DMC1-SSDS signal within a few Mb from the q-arm telomeres has been observed in male mice that express a GAL4BD-SPO11 fusion protein instead of SPO11 [8]. This observation, which indicates a specific property of GAL4BD-SPO11 and/or its low abundance, highlights a specific control of DSB activity in subtelomeric regions in males. Second, in wild-type mice, the level of DMC1 SSDS signal relative to genome average is lower near chromosome ends [9]. This may be due to lower DSB activity and/or to increased DMC1 turnover. Indeed, DSB activity monitored by detecting the oligonucleotides covalently linked to SPO11 (SPO11-oligos) showed a decrease of the ratio between DMC1-SSDS and SPO11-oligos within the 5 Mb subtelomeric regions of autosomes [10], compatible with increased DMC1 turnover or reduced DMC1 loading. These data provide independent evidence for specific controls of DSB activity of these chromosome ends in wild-type mice. It is also possible that the XY phenotype of SPO11β-only (the long isoform of SPO11) mice, with decreased DSB activity in the PAR [11], actually applies to other chromosome ends. As we have observed in the context of the *Top6bl* mutants, reduction of DSB activity near chromosome ends has no detectable consequence on autosome synapsis, at least at the microscopic level. This would imply that the SPO11α isoform (the short SPO11 isoform) is not specifically required for recombination between X and Y but required for recombination near chromosomal ends. As SPO11α is lacking the interaction domain with TOPOVIBL, it is not expected to be catalytically active. It may rather repress an inhibitor of TOPOVIL. One possibility is that this inhibitor interferes with REC114-TOPOVIBL interaction. Such scenario, although certainly others can be envisioned, would fit with the potential similarity of phenotypes between *Spo11β*-only and *Top6bl*$^{\Delta17Ct/\Delta17Ct}$ male mice.

#### Table S1. Data collection and refinement statistics

| | **TOPVIBL-REC114** |
|---|---|
| **Data collection** | |
| Space group | $P6_122$ |
| Cell dimensions | |
| *a*, *b*, *c* (Å) | 108,6, 108,6, 83.4 |
| α β γ (°) | 90, 90, 120 |
| Resolution (Å) | *50-2.2 (2.85-2.7)*[a] |
| $R_{merge}$ (%) | *25.7 (546.2)* |
| I / σI | *12.39 (0.94)* |
| $CC_{½}$ (%) | *100 (37.5)* |
| Completeness (%) | *99.8 (99.6)* |
| Redundancy | *18.3 (19.3)* |
| **Refinement** | |
| Resolution (Å) | 94.03-2.2 |
| No. reflections | 12125 |
| $R_{work}$ / $R_{free}$ | 0.232/0.256 |
| *B*-factors | 60.5 |
| R.m.s. deviations | |
| Bond lengths (Å) | 0.005 |
| Bond angles (°) | 1.412 |

[a] Values in parentheses are for highest resolution shell.

**Table S2**

| | Unchanged-Decreased-Increased | | | | | Unchanged-Decreased | | | | |
|---|---|---|---|---|---|---|---|---|---|---|
| | Subtelo1M | Subtelo2M | Subtelo3M | Subtelo4M | Subtelo5M | Subtelo1M | Subtelo2M | Subtelo3M | Subtelo4M | Subtelo5M |
| chr1 | 0,19688 | 1 | 1 | 1 | 1 | 0,337954 | 0,707152 | 0,95226 | 1 | 0,30121 |
| chr2 | 0,19688 | 1 | 0,9538 | 1 | 1 | 0,118562 | 0,728138 | 0,170249 | 0,26614 | 0,26614 |
| chr3 | NA | 0,09331 | 0,00059 | 0 | 0,00062 | NA | 0,209354 | 0,000178 | 1,00E-06 | 8,70E-05 |
| chr4 | 2,00E-05 | 0 | 0 | 0 | 0 | 1,20E-05 | 0 | 0 | 0 | 0 |
| chr5 | NA | 0,09472 | 1 | 1 | 1 | NA | 0,214951 | 0,530377 | 1 | 1 |
| chr6 | 0,1378 | 0,00765 | 1 | 1 | 1 | 0,275433 | 0,00955 | 0,263484 | 0,426418 | 0,848232 |
| chr7 | NA | 1 | 1 | 1 | 1 | NA | 0,867484 | 1 | 1 | 0,92624 |
| chr8 | 0,00204 | 9,00E-05 | 0,00548 | 0,2626 | 1 | 0,005766 | 5,50E-05 | 0,001712 | 0,039004 | 0,37051 |
| chr9 | 2,00E-05 | 0 | 0 | 0 | 0 | 5,00E-06 | 0 | 0 | 0 | 0 |
| chr10 | 0,07514 | 0,00201 | 0,06632 | 0,68638 | 1 | 0,213463 | 0,003835 | 0,016055 | 0,106033 | 0,211257 |
| chr11 | 0,00041 | 0,00046 | 0,13714 | 0,58528 | 1 | 0,000331 | 0,000331 | 0,022116 | 0,06639 | 0,301525 |
| chr12 | NA | 1 | 1 | 1 | 1 | NA | 1 | 1 | 0,877914 | 0,708196 |
| chr13 | NA | 0,00056 | 0 | 0 | 0 | NA | 0,000478 | 0 | 0 | 0 |
| chr14 | 0,00156 | 0,20581 | 0,25 | 1 | 1 | 0,004174 | 0,094238 | 0,094238 | 0,886754 | 0,784146 |
| chr15 | 0,00041 | 0,58244 | 0,25 | 1 | 1 | 0,001482 | 0,189291 | 0,0572 | 0,170896 | 0,329774 |
| chr16 | 0,85779 | 1 | 1 | 1 | 1 | 0,596137 | 1 | 1 | 0,932596 | 1 |
| chr17 | NA | NA | NA | 1 | 0,02142 | NA | NA | NA | 0,622188 | 0,003701 |
| chr18 | NA | NA | 1 | 1 | 1 | NA | NA | 1 | 0,644324 | 0,875619 |
| chr19 | 0 | 0 | 0,00162 | 0,05963 | 0,26777 | 0 | 0 | 0,00019 | 0,004521 | 0,020866 |
| chrX | NA | 0 | 0 | 0 | 0 | NA | 0 | 0 | 0 | 0 |

**Table S3**

| | Unchanged-Decreased-Increased | | | | | Unchanged-Decreased | | | | |
|---|---|---|---|---|---|---|---|---|---|---|
| | Subtelo1M | Subtelo2M | Subtelo3M | Subtelo4M | Subtelo5M | Subtelo1M | Subtelo2M | Subtelo3M | Subtelo4M | Subtelo5M |
| chr1 | 0,1270438 | 0,15828855 | 0,12043018 | 0,10304826 | 0,01316773 | 0,8633 | 0,9237 | 0,9167 | 1 | 0,6562 |
| chr2 | 0,12682662 | 7,1955E-05 | 0 | 0 | 0 | 0,5843 | 0,004 | 0,0003 | 0,0002 | 0,0002 |
| chr3 | NA | 1 | 0,71112522 | 0,28593636 | 0,12739808 | NA | 1 | 1 | 1,00E+00 | 1,00E+00 |
| chr4 | 1,11E-02 | 0 | 0 | 0 | 0 | 2,24E-01 | 0 | 0 | 0 | 0 |
| chr5 | NA | 0,14812875 | 0,16640265 | 0,01585219 | 1,599E-05 | NA | 0,4886 | 0,7521 | 0,2018 | 0,001 |
| chr6 | 0,1270438 | 0,00840843 | 3,5977E-05 | 0,00032779 | 0,00204472 | 0,5843 | 0,0648 | 0,0015 | 0,0032 | 0,025 |
| chr7 | 1 | 0,32901843 | 0,2272917 | 0,00868691 | 0,00825266 | 1 | 1 | 1 | 0,1606 | 0,2065 |
| chr8 | 0,36200843 | 7,58E-01 | 0,45403898 | 0,09875076 | 0,00045332 | 1 | 1,00E+00 | 1 | 1 | 0,0759 |
| chr9 | 1,27E-01 | 0,00012952 | 0 | 0 | 0 | 7,31E-01 | 0,0059 | 0,0004 | 0,0002 | 0,0001 |
| chr10 | 1 | 0,41195078 | 0,00351679 | 7,1955E-05 | 1,599E-05 | 1 | 1 | 0,0292 | 0,0021 | 0,0006 |
| chr11 | 0,1270438 | 0,00840843 | 0,00074011 | 0,00077711 | 0,00099428 | 0,5843 | 0,0552 | 0,0106 | 0,013 | 0,0417 |
| chr12 | 1 | 1 | 0,04625385 | 0,09700534 | 0,94164587 | 1 | 1 | 0,4673 | 1 | 1 |
| chr13 | 0,1270438 | 0 | 0 | 0 | 0 | 0,5843 | 0 | 0 | 0 | 0 |
| chr14 | 0,72015086 | 0,23130145 | 0,06851895 | 0 | 0 | 1 | 1 | 0,7521 | 0,0001 | 0,0009 |
| chr15 | 0,22391028 | 7,1955E-05 | 0 | 0 | 0 | 1 | 0,004 | 0,001 | 0 | 0 |
| chr16 | 0,71792936 | 0,03847982 | 0,00421655 | 1,0279E-05 | 0 | 1 | 0,4886 | 0,1201 | 0,0035 | 0,0002 |
| chr17 | 1 | 1 | 1 | 1 | 0,24520595 | 1 | 1 | 1 | 1 | 0,6032 |
| chr18 | NA | 1 | 1 | 1 | 1 | NA | 1 | 1 | 1 | 1 |
| chr19 | 0,21142886 | 0,12138774 | 0,16640265 | 0,17873994 | 0,10121791 | 0,7313 | 0,4886 | 1 | 1 | 1 |
| chrX | 1 | 0,02995118 | 0,00357375 | 0,01154874 | 0,02394176 | 1 | 0,4886 | 0,103 | 0,3873 | 1 |

### Table S4

| | | |
|---|---|---|
| Guides for CirspR | 6509_Top6bl_sgRNA1 | CCCGCAGGACTTGTGGCTAC |
| | 6510_Top6bl_sgRNA2 | GCAGGACTTGTGGCTACAGG |
| | 6511_Top6bl_sgRNA3 | TCCTGTAGCCACAAGTCCTG |
| Donor for CrispR | 6512_Top6bl_donor1 | TTCCTTGGCTTGCTGGGCGTCTCAGAGAGAATGCTTAGGACCTGTGTCCCGGATTCAGCCACTCGGACAGATTGGATACCTCCTGCAGCGCCAAGTCCTGCGGGAGCAGAGGAGGCGGTGGTGAGGA |
| | 6513_Top6bl_donor2 | GCGGCCTGGAGGTACCGCAGCGAGAGCAGGGCGGGTGCGGGCCTGAGCTACCACCGCCGGGTCCTCACCACCGCCTCCTCTGCTCCCGCAGGACTTGGCGCTGCAGGAGGTATCCAATCTGTCCGAG |
| Genoptyping | oliAN 320 | CAACCATGGCAGGGAAGAATC |
| | oliAN 3201 | TACCGACAGTAAACGCAGCC |
| RT-PCR *Top6bl* | Oli63 | GGACCAGCTCTGCTACTTTT |
| | Oli70 | CAGAGAGAATGCTTAGGACC |
| RT-PCR *Spo11* | Spo11:116U22 | CTGTTGGCCATGGTGAAGAGAG |
| | Spo11:655L22 | TGTGCCGACTAACATTCAAGGA |

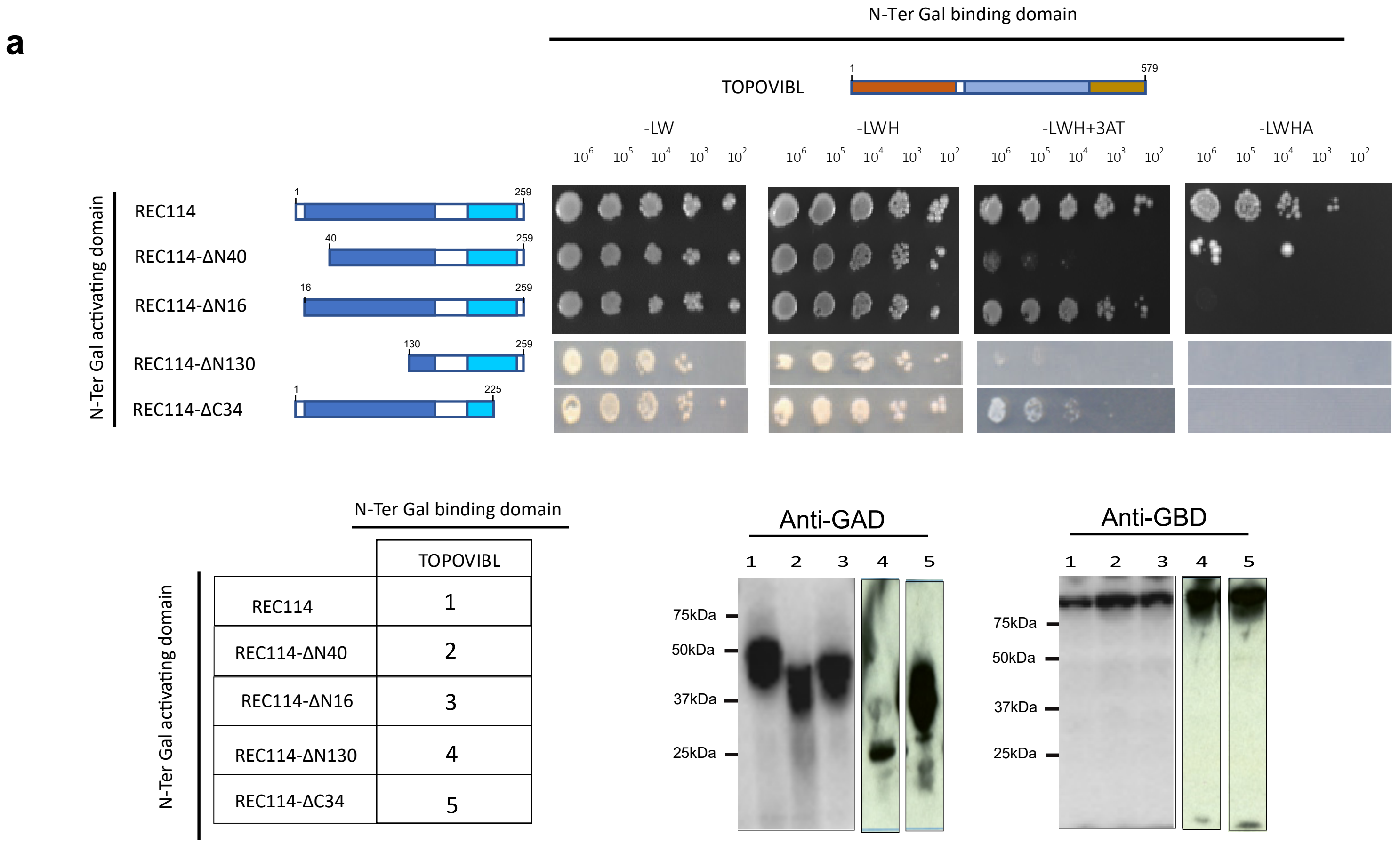

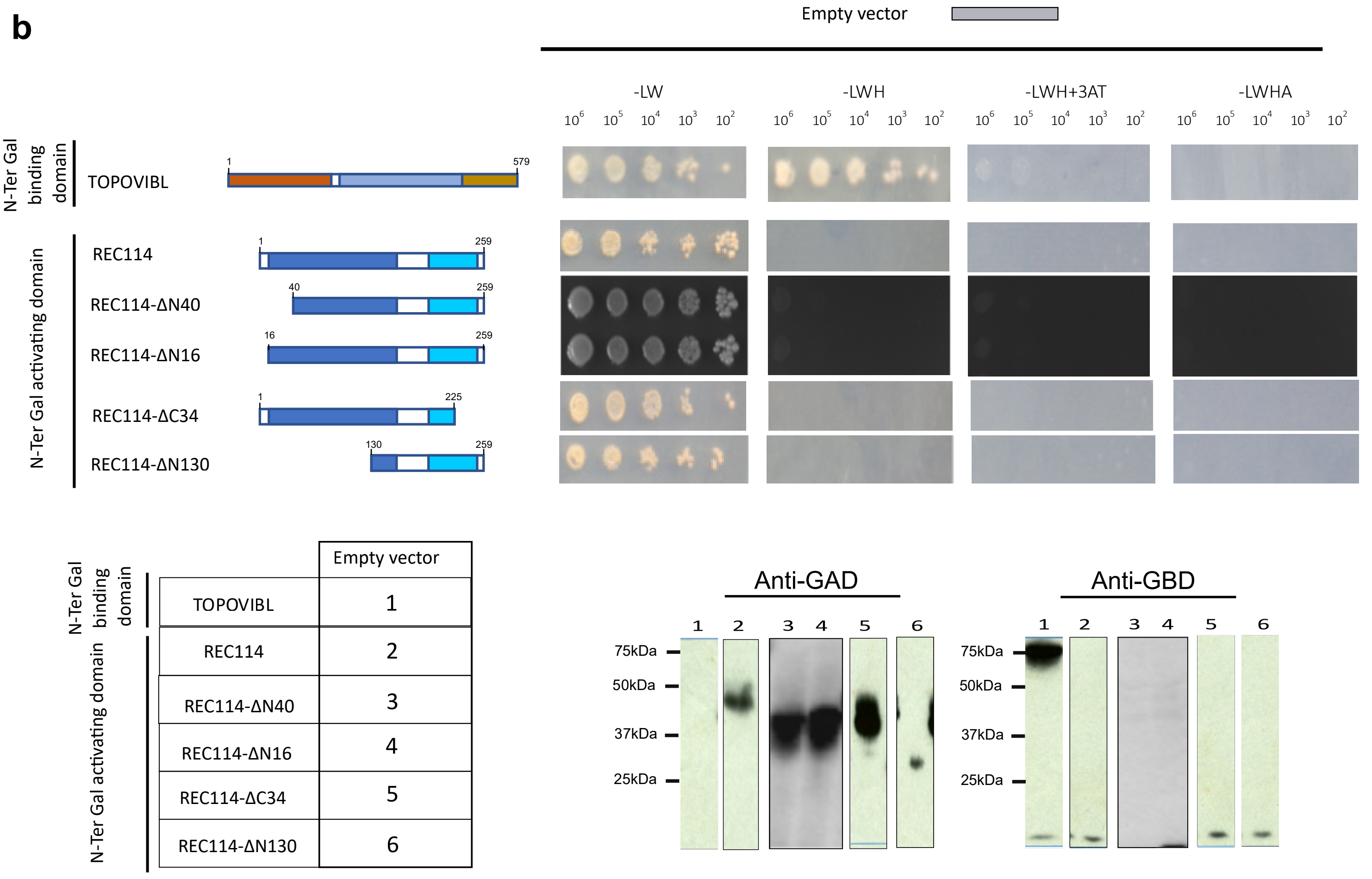

**Supplementary Figure 1-1**

**c**

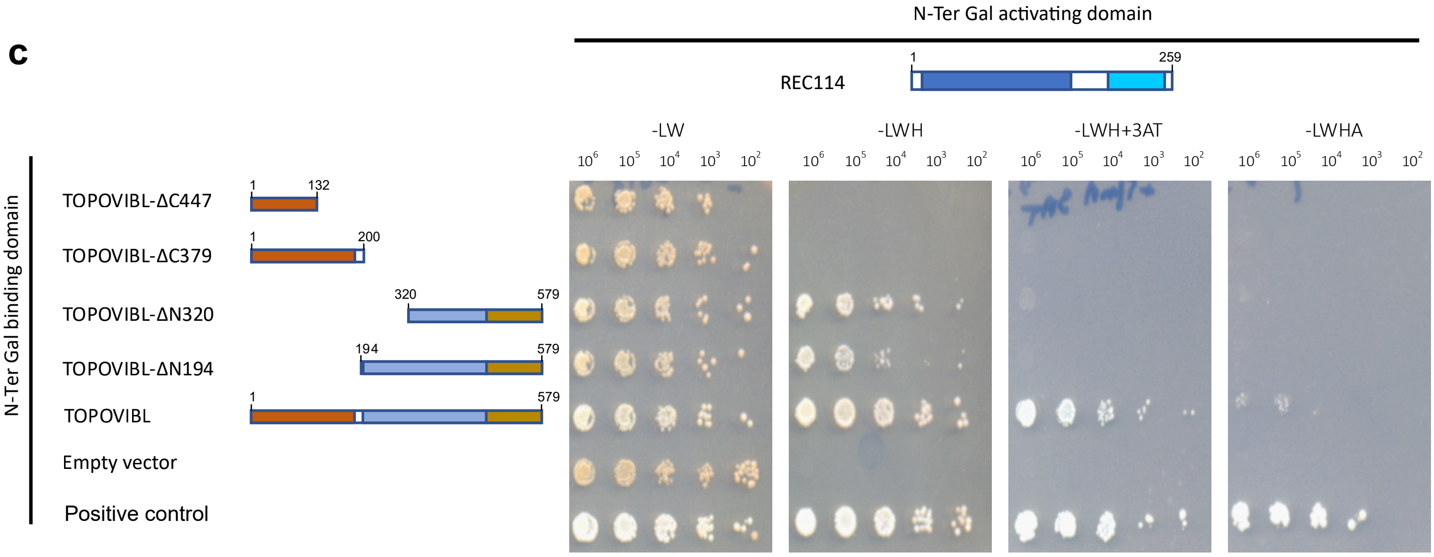

N-Ter Gal activating domain

N-Ter Gal binding domain

| | REC114-FL |
|---|---|
| TOPOVIBL-ΔC447 | 1 |
| TOPOVIBL-ΔC379 | 2 |
| TOPOVIBL-ΔN320 | 3 |
| TOPOVIBL-ΔN194 | 4 |
| TOPOVIBL | 5 |
| Empty vector | 6 |

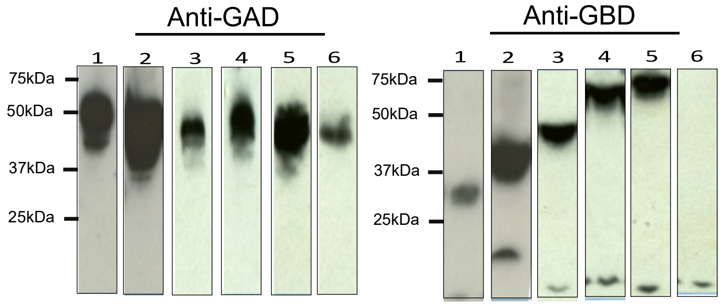

**d**

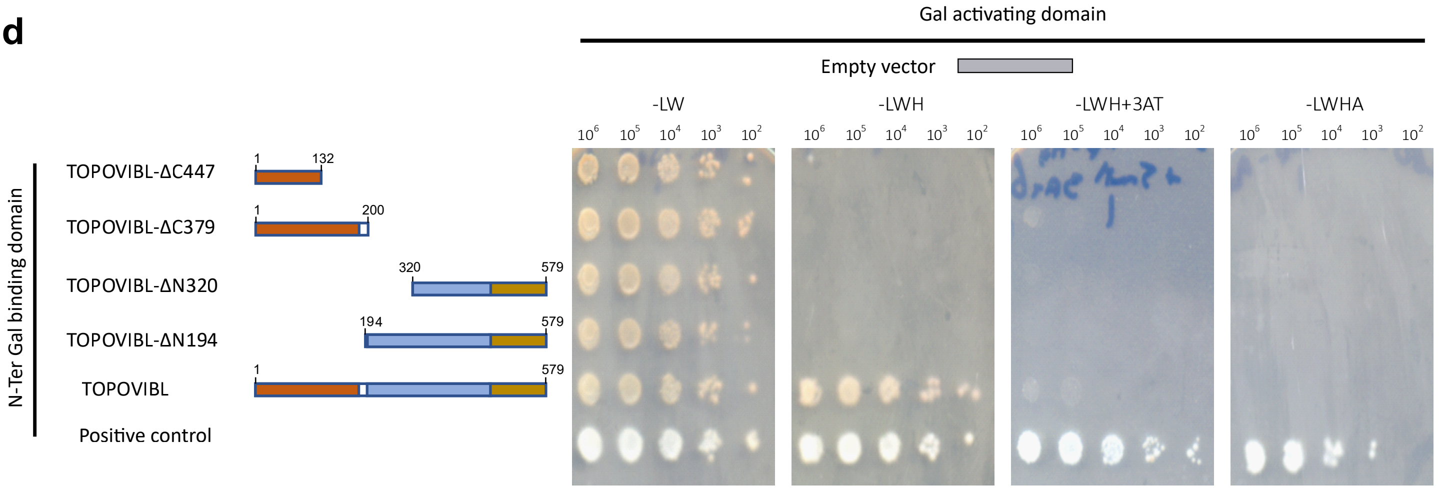

N-Ter Gal activating domain

N-Ter Gal binding domain

| | Empty vector |
|---|---|
| TOPOVIBL-ΔC447 | 1 |
| TOPOVIBL-ΔC379 | 2 |
| TOPOVIBL-ΔN320 | 3 |
| TOPOVIBL-ΔN194 | 4 |
| TOPOVIBL | 5 |

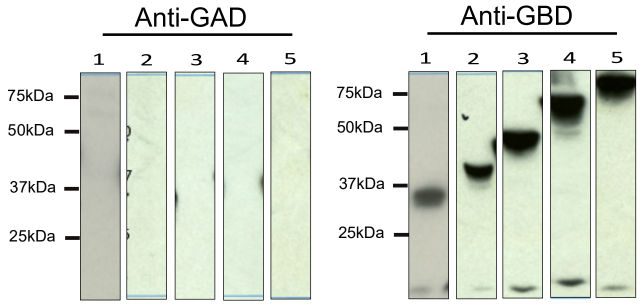

**Supplementary Figure 1-2**

**e**

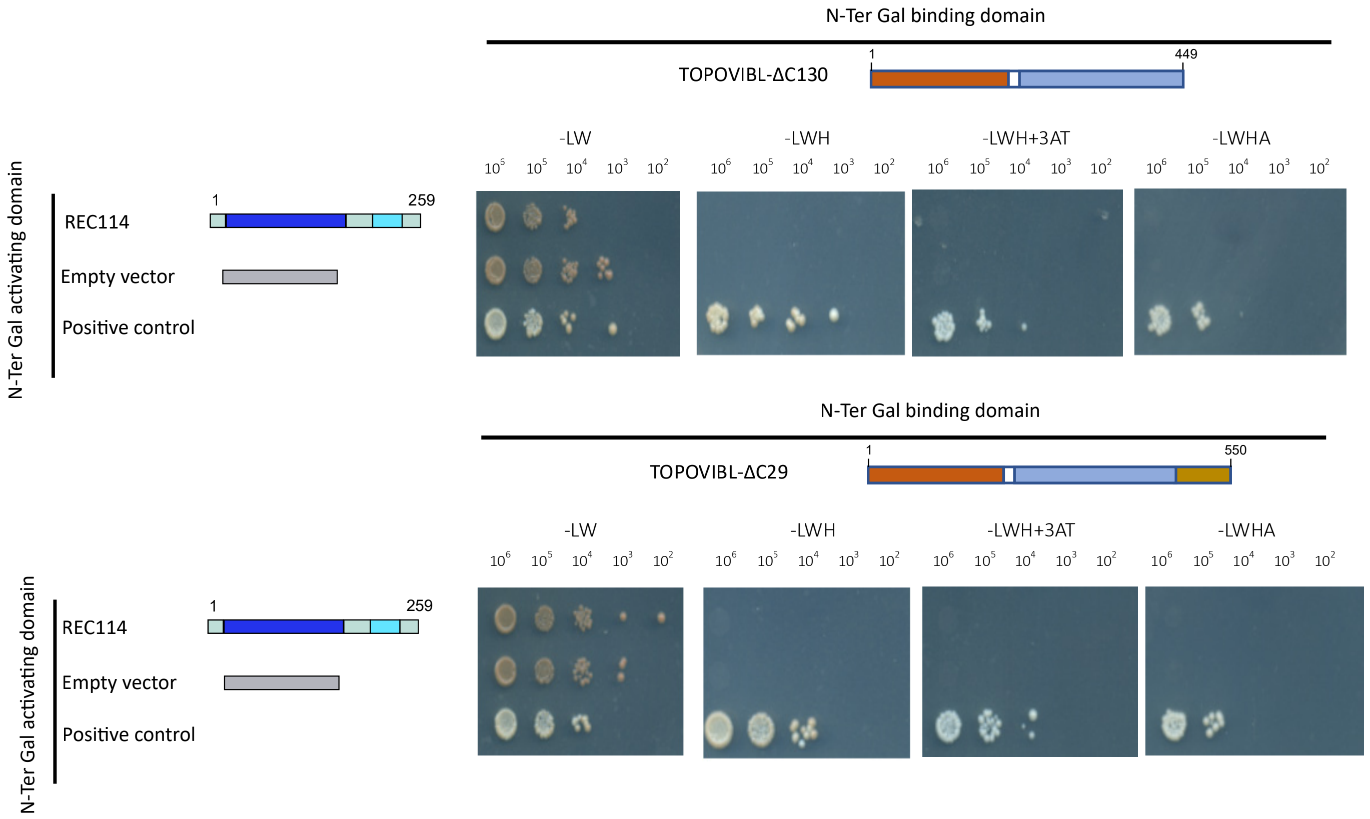

N-Ter Gal activating domain

| N-Ter Gal binding domain | REC114-FL | Empty vector |
|---|---|---|
| TOPOVIBL-ΔC130 | 1 | 2 |
| TOPOVIBL-ΔC29 | 3 | 4 |

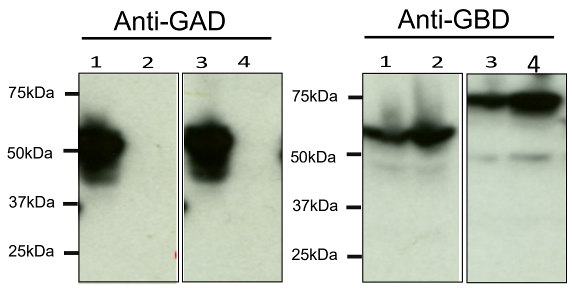

**f**

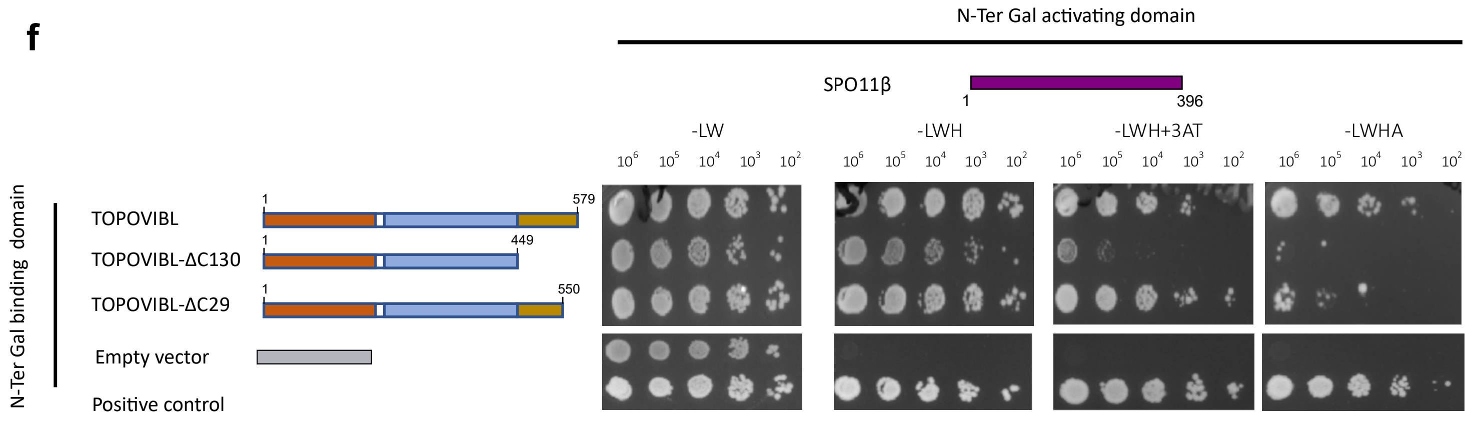

N-Ter Gal activating domain

| N-Ter Gal binding domain | SPO11β |
|---|---|
| TOPOVIBL | 1 |
| TOPOVIBL-ΔC29 | 2 |
| TOPOVIBL-ΔC130 | 3 |
| Empty vector | 4 |

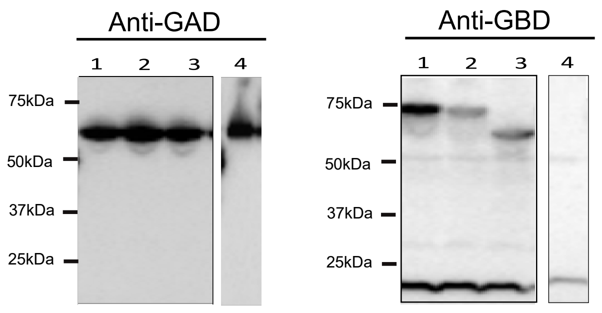

**Supplementary Figure 1-3**

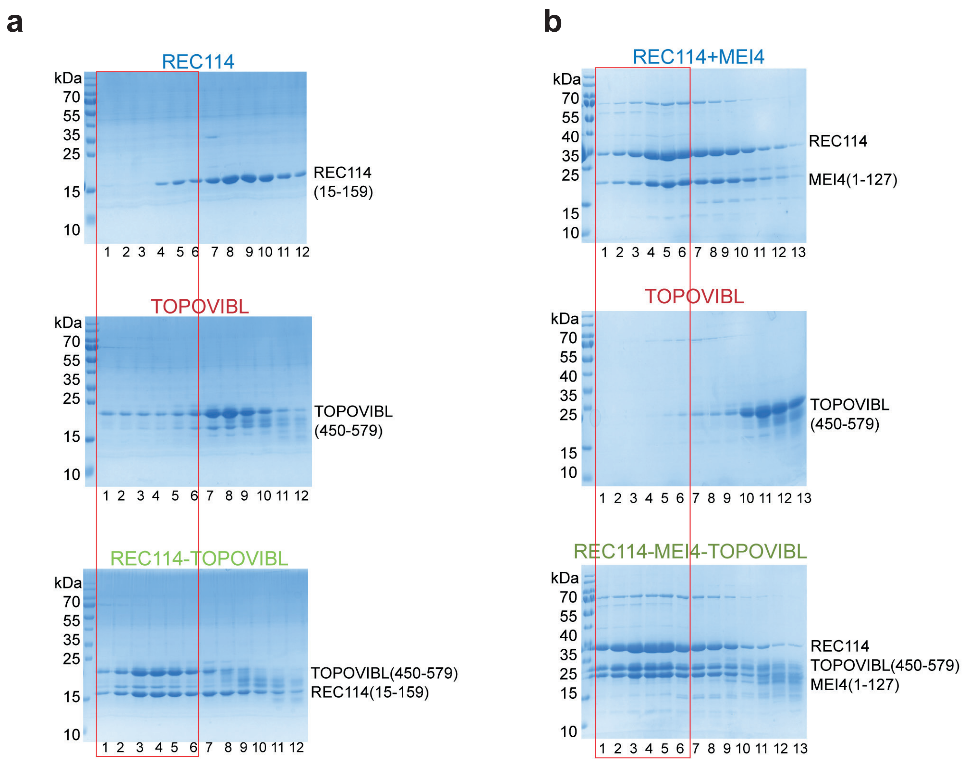

#### Supplementary Figure 2

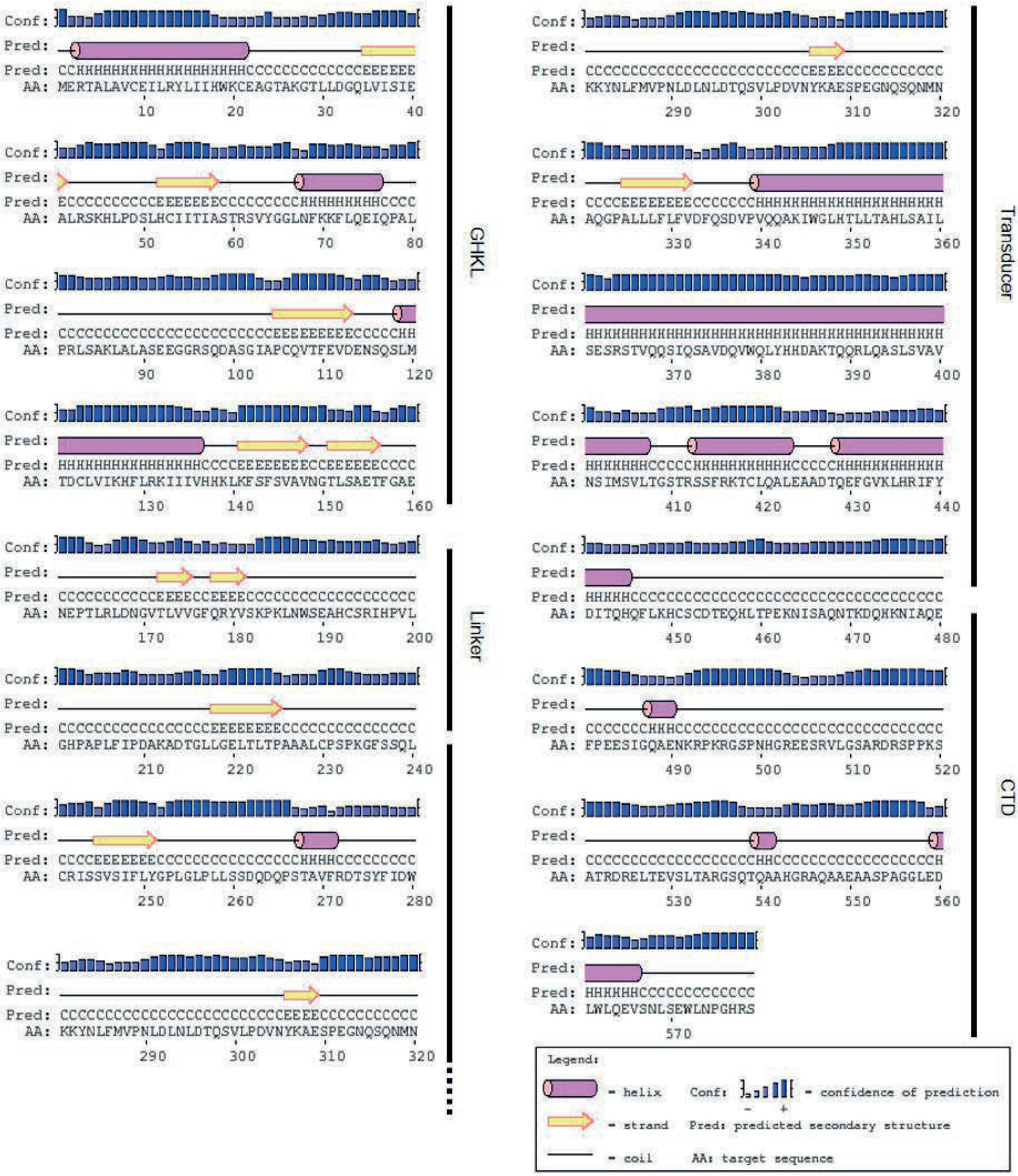

**Supplementary Figure 3**

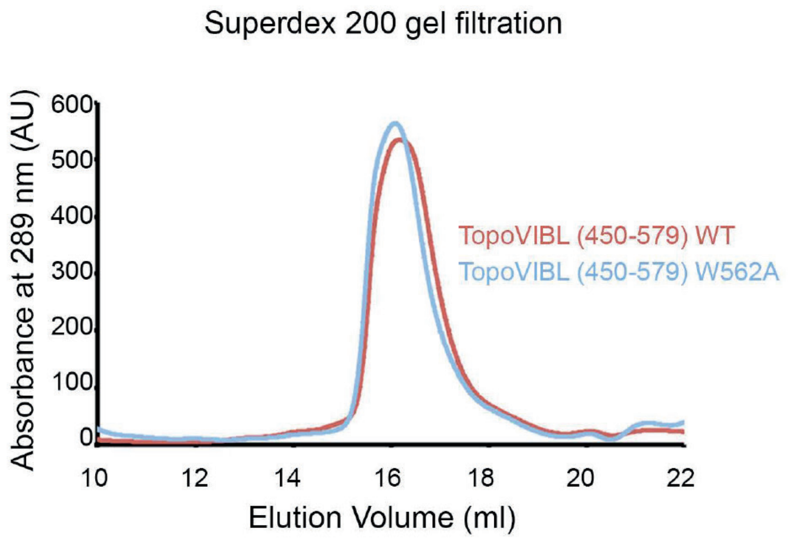

**Supplementary Figure 4**

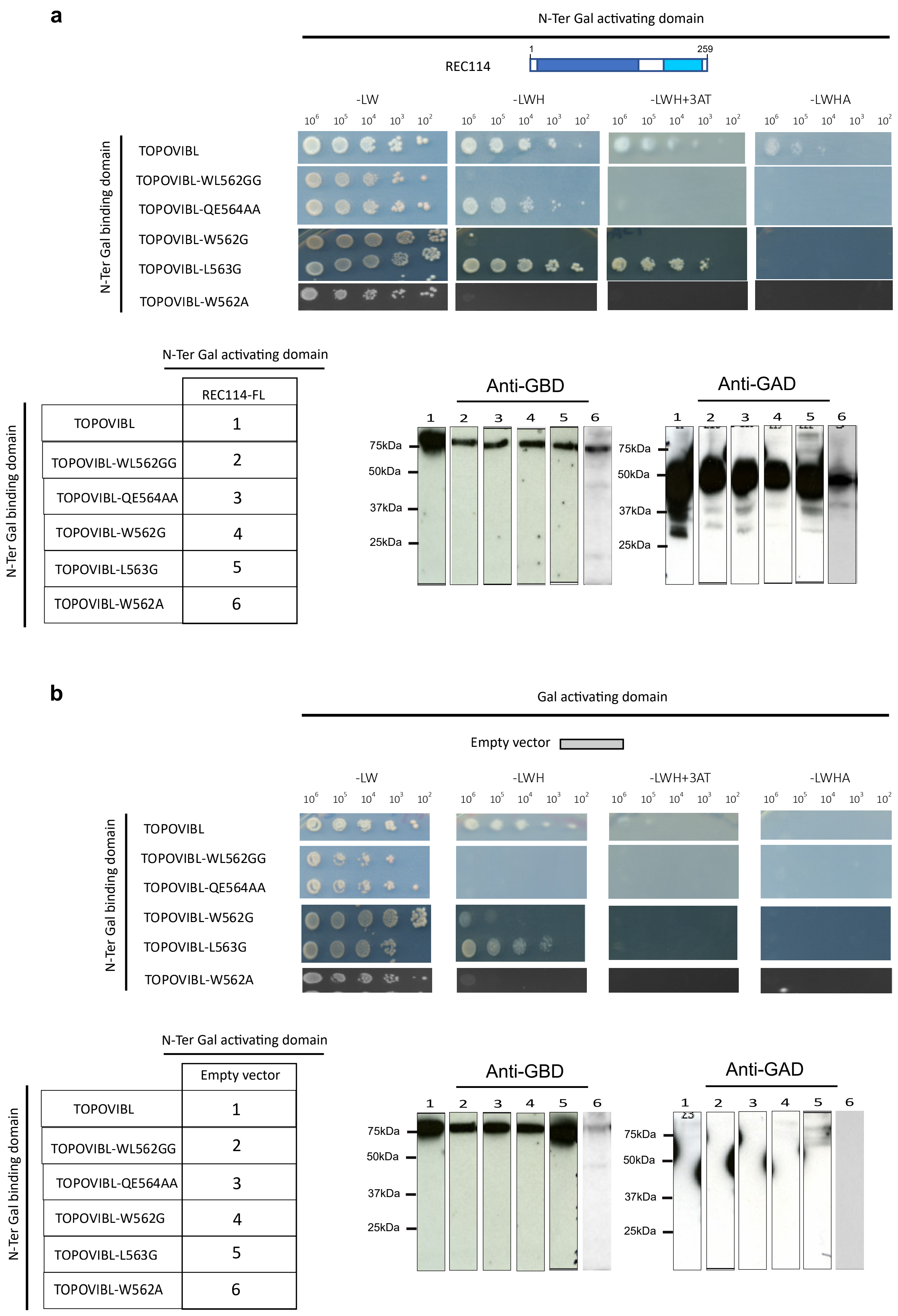

**Supplementary Figure 5-1**

**c**

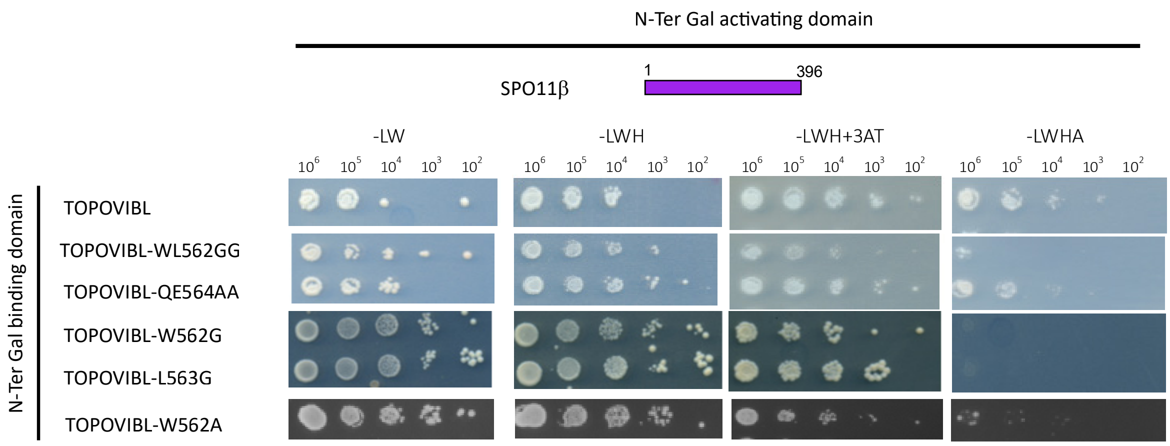

N-Ter Gal activating domain / N-Ter Gal binding domain

| | SPO11B |
|---|---|
| TOPOVIBL | 1 |
| TOPOVIBL-WL562GG | 2 |
| TOPOVIBL-QE564AA | 3 |
| TOPOVIBL-W562G | 4 |
| TOPOVIBL-L563G | 5 |
| TOPOVIBL-W562A | 6 |

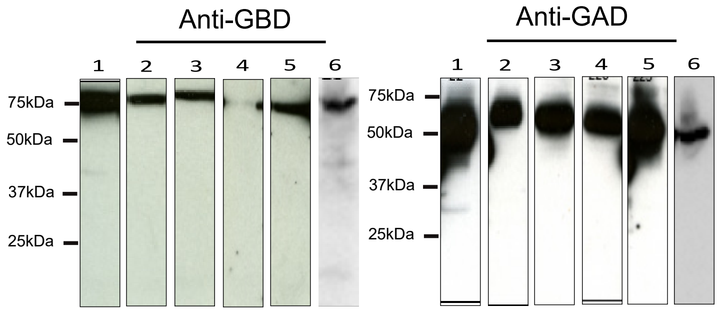

**Supplementary Figure 5-2**

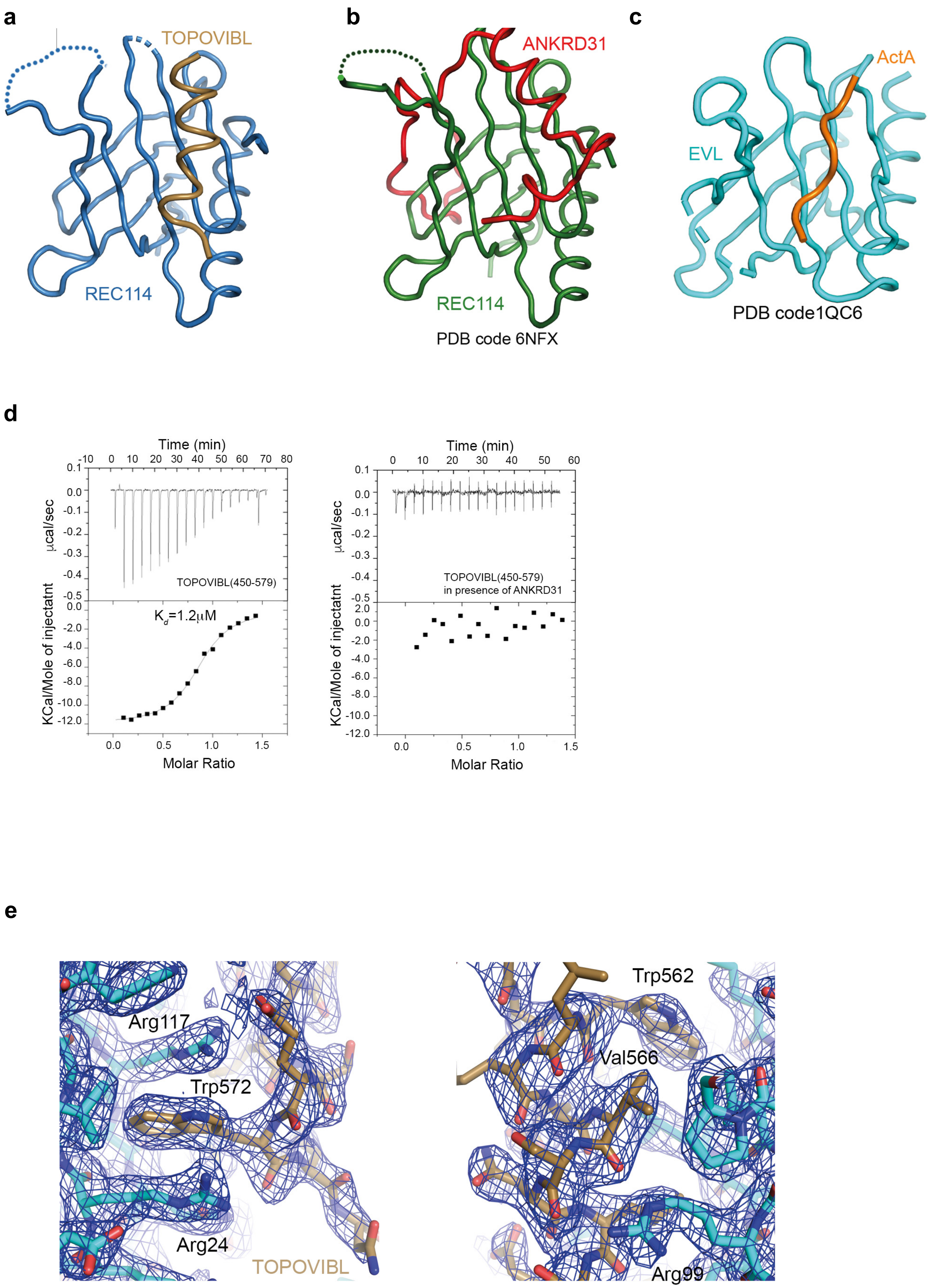

**Supplementary Figure 6**

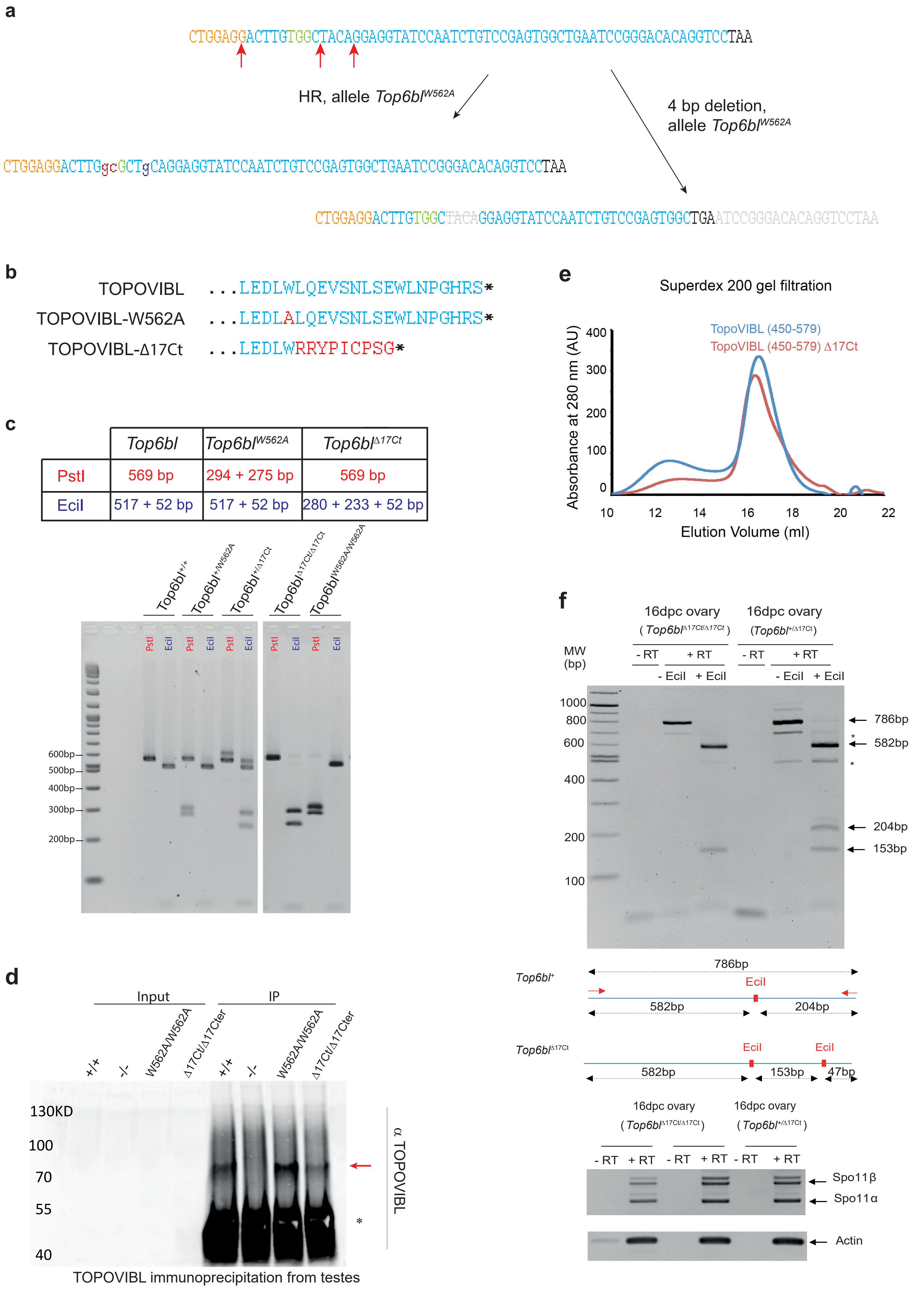

**Supplementary Figure 7**

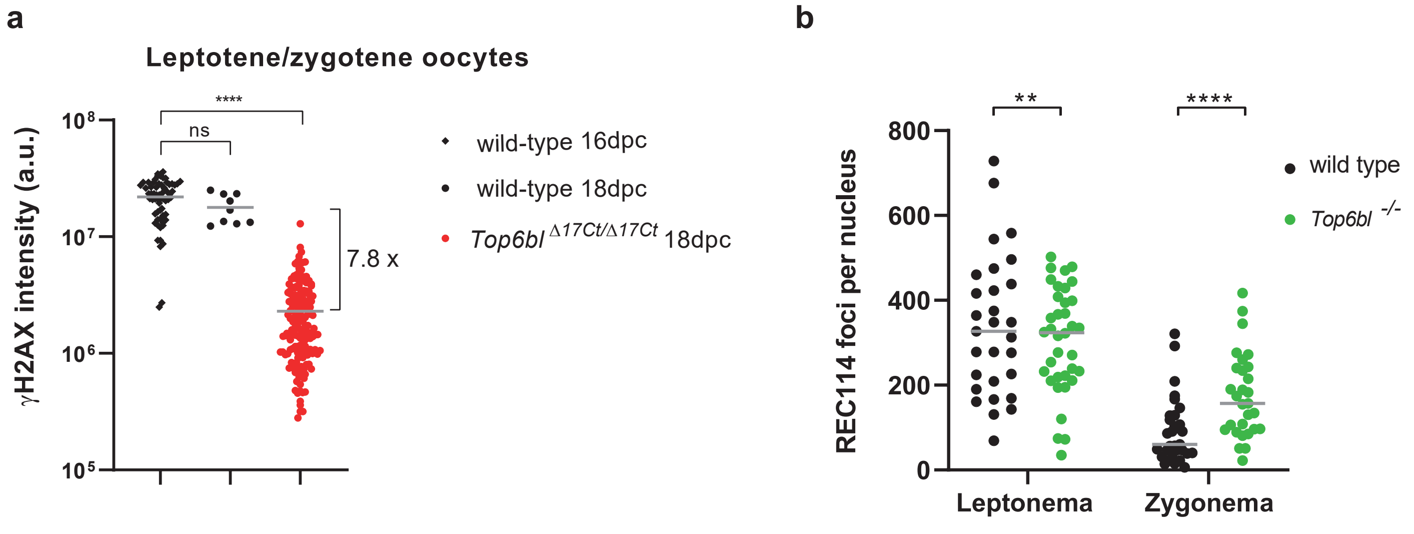

**Supplementary Figure 8**

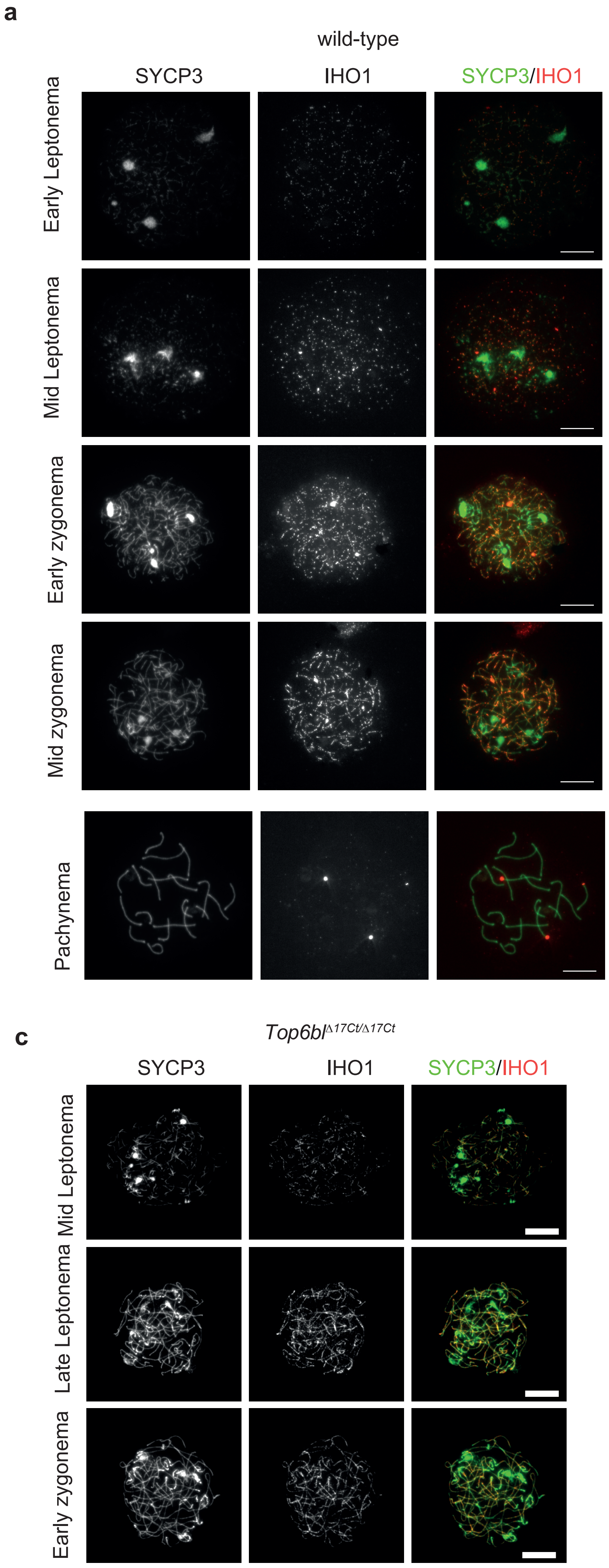

**Supplementary Figure 9**

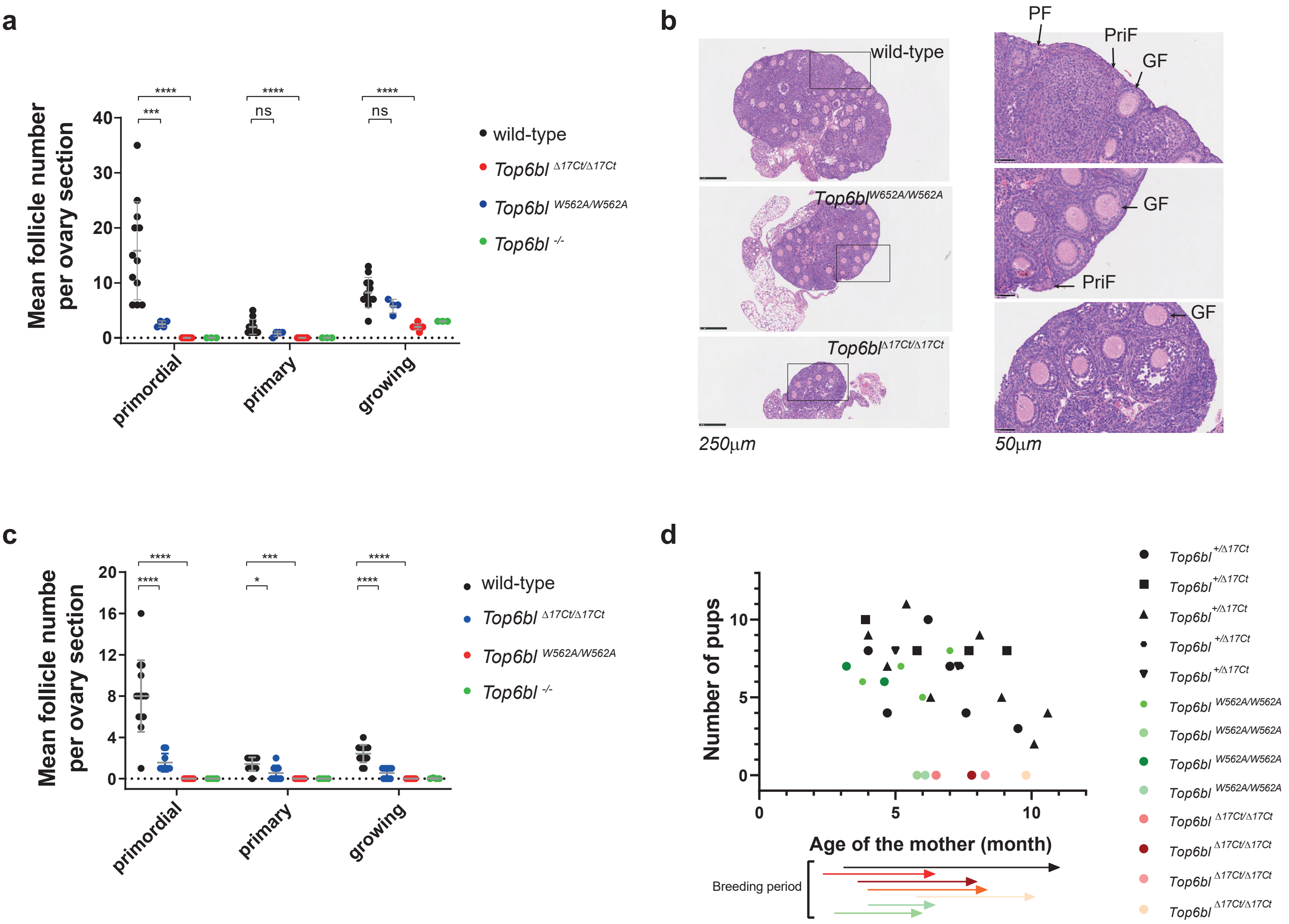

#### Supplementary Figure 10

**Supplementary Figure 11**

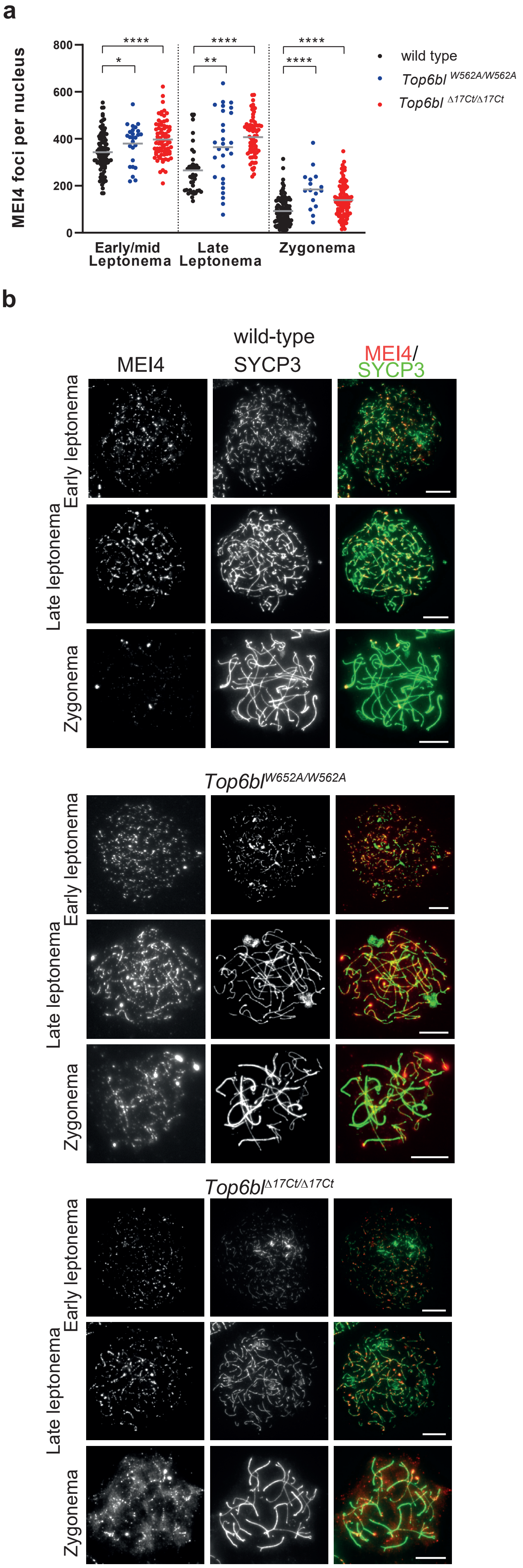

### Supplementary Figure 12

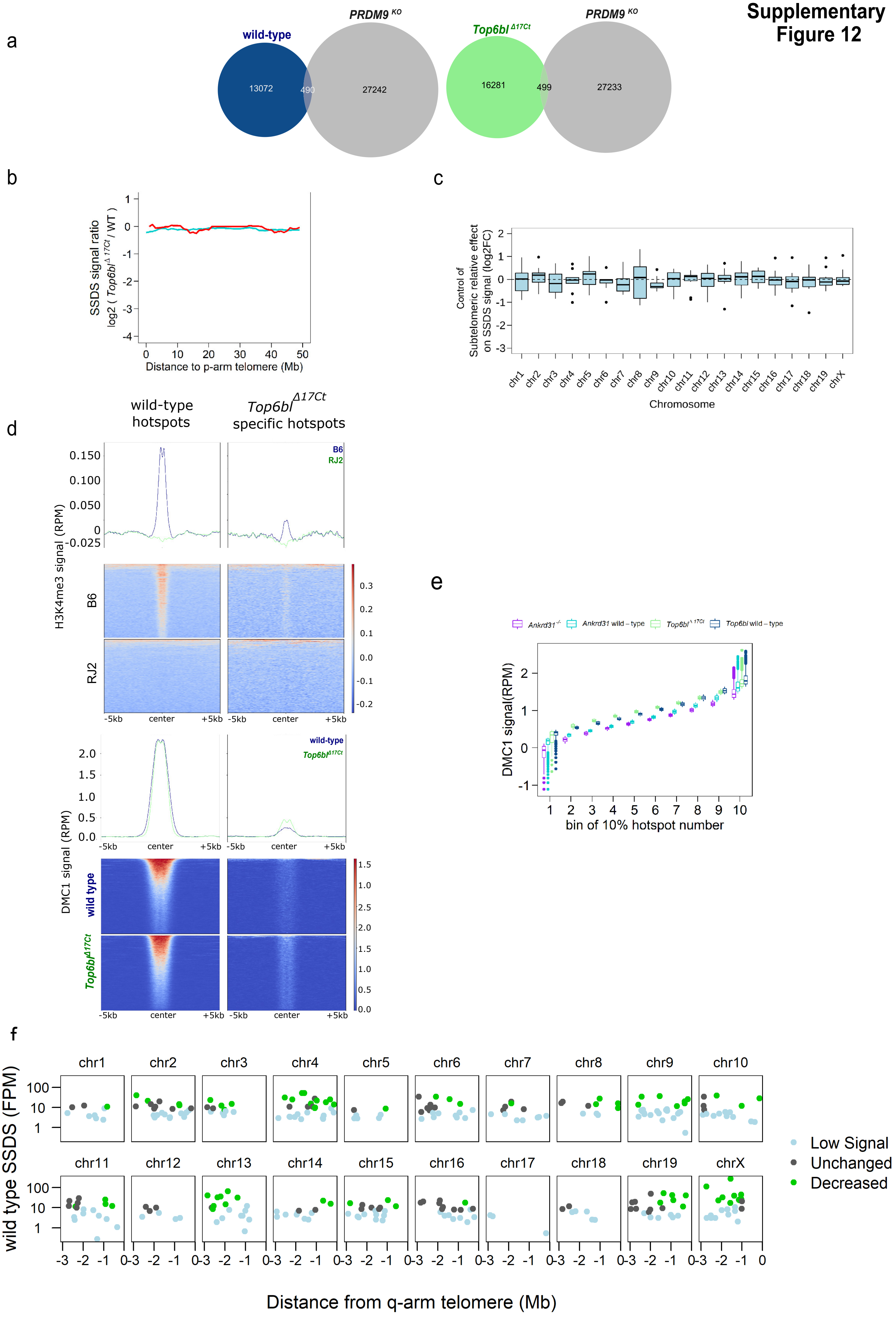

### Supplementary Figure 13

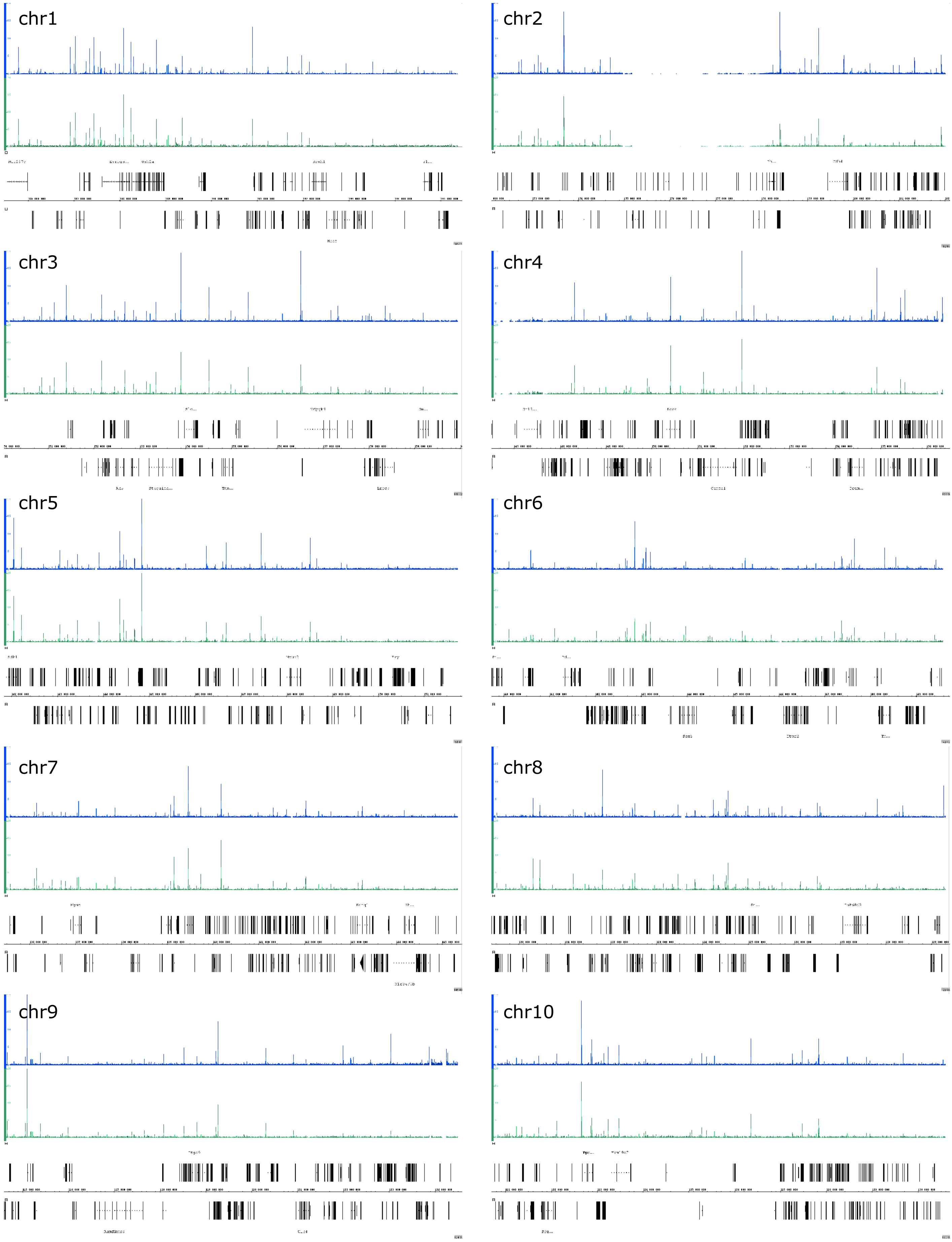

### Supplementary Figure 14

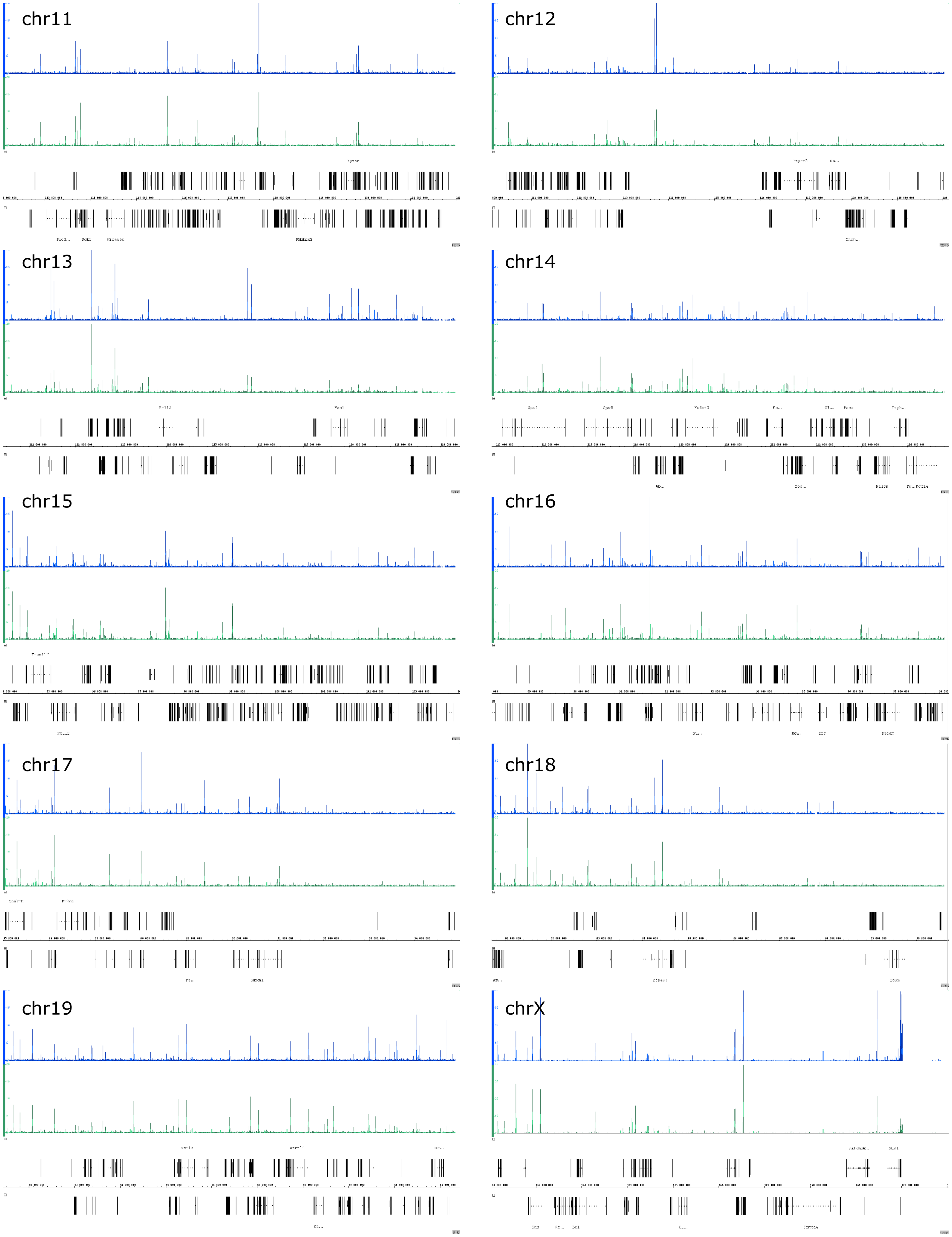

### Supplemental Figure S15

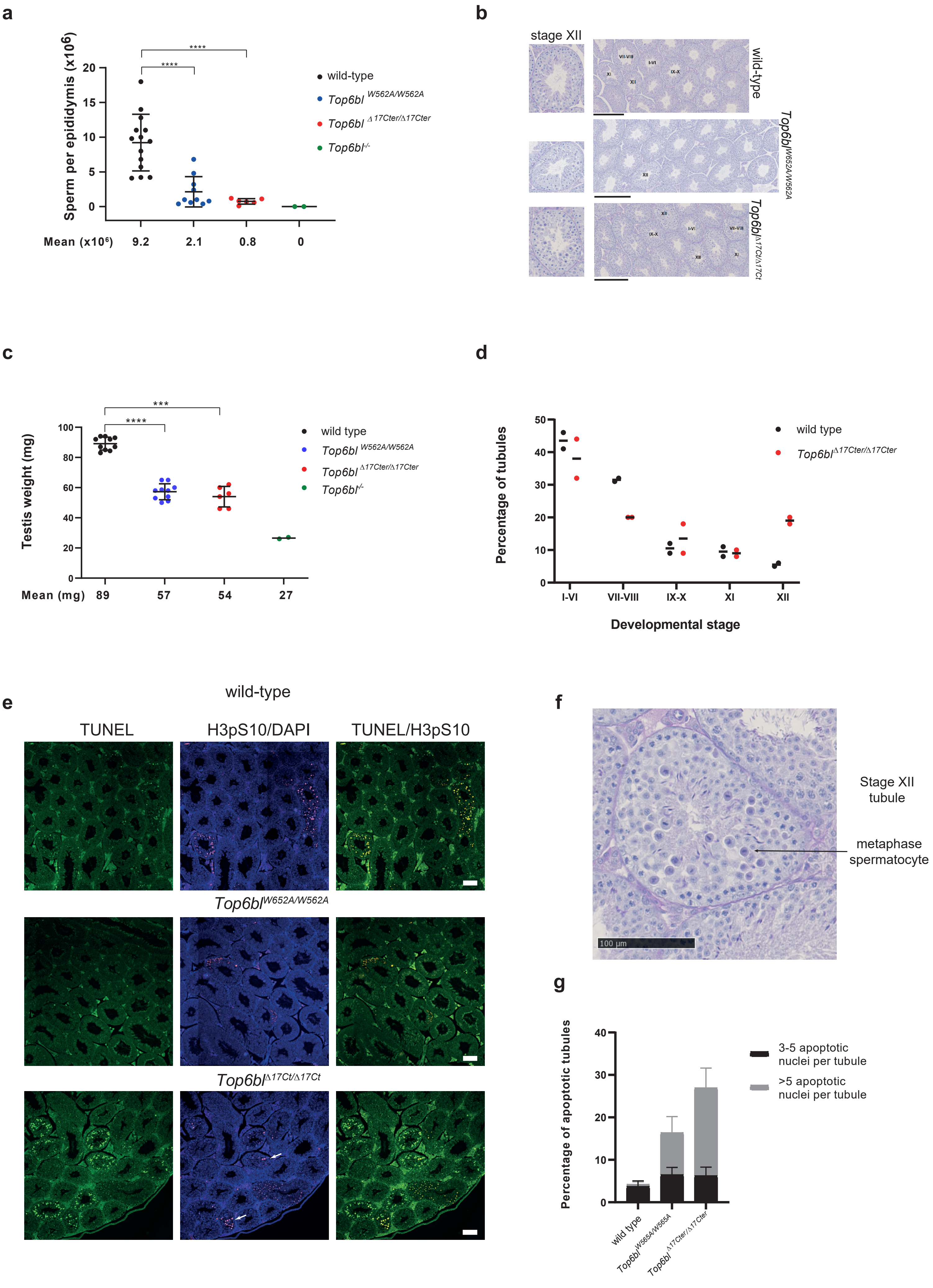

#### Supplementary Figure S16

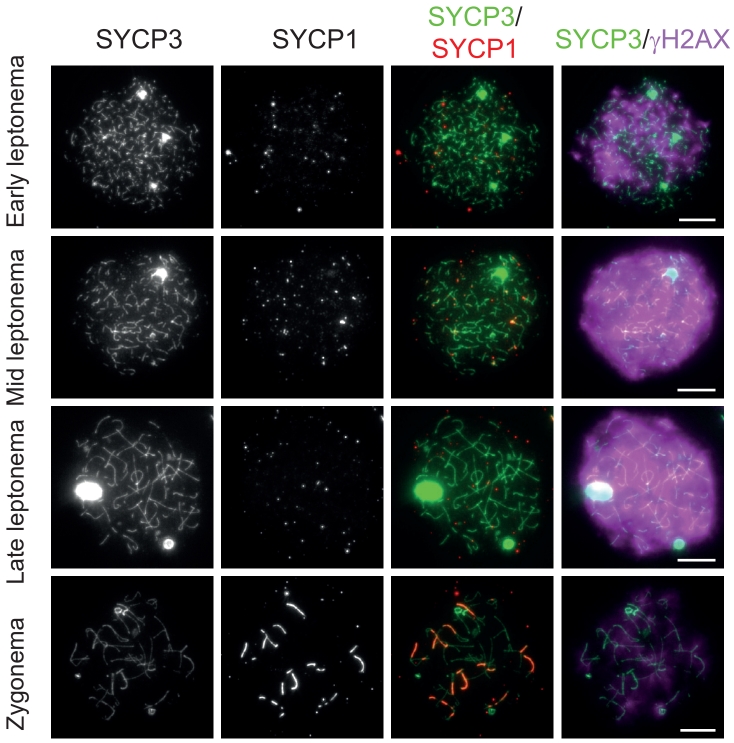

#### Supplementary Figure 17

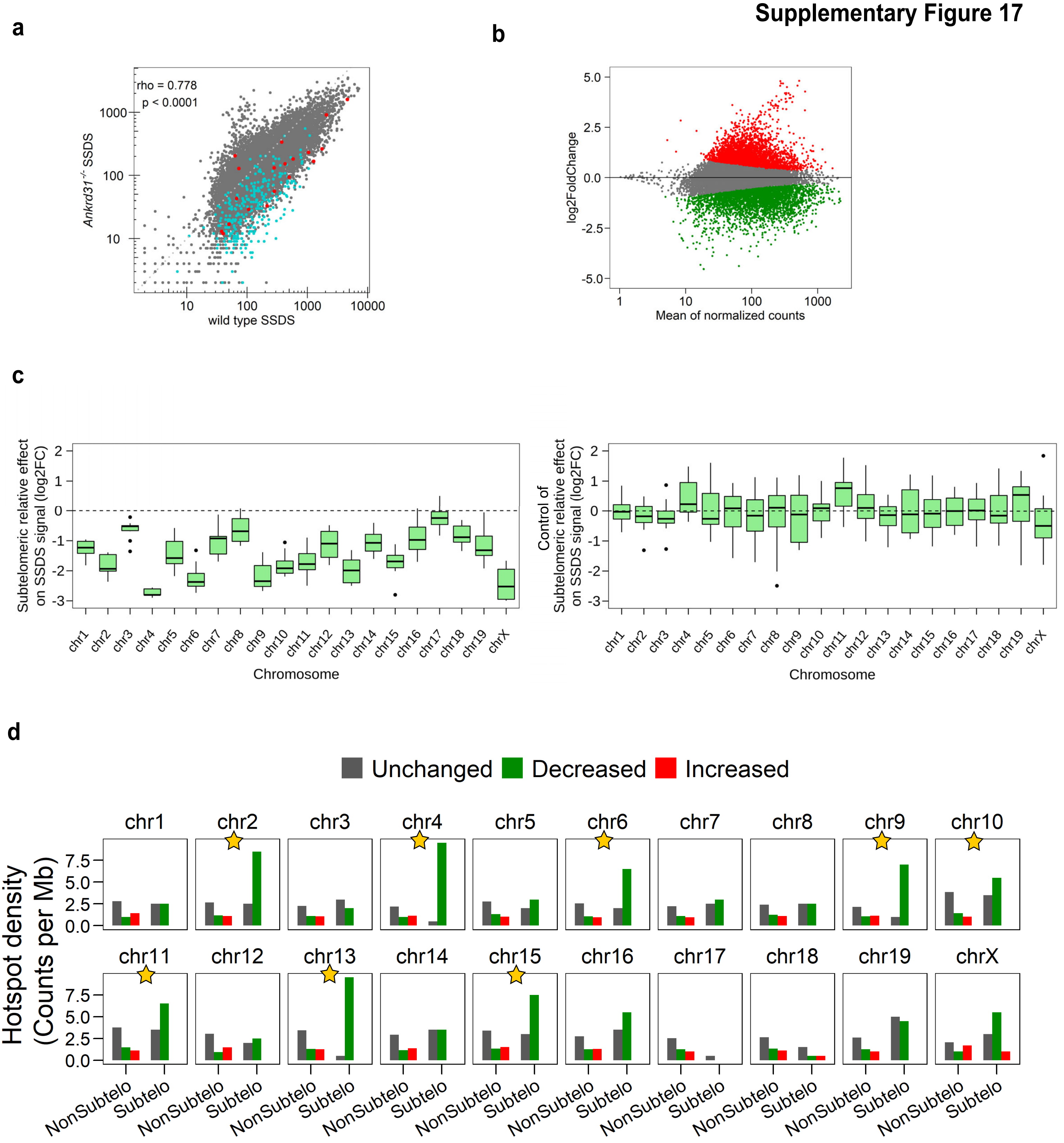
